# A complex view of GPCR signal transduction: Molecular dynamics of the histamine H3 membrane receptor

**DOI:** 10.1101/604793

**Authors:** L. D. Herrera-Zúñiga, L. M. Moreno-Vargas, L. Ballaud, J. Correa-Basurto, D. Prada-Gracia, D. Pastré, P. A. Curmi, J. M. Arrang, R. C. Maroun

**Author notes:** LDHZ: Tecnológico de Estudios Superiores de Chicoloapan, Loma de Guadalupe., 56380 Ejido de Chicoloapan, Mexico State, MEXICO. LMMV: Computational Biology and Drug Design Research Unit. Federico Gómez Children’s Hospital of Mexico City, MEXICO. JCB: Laboratory for the Design and Development of New Drugs and Biotechnological Innovation, SEPI-ESM, Mexico City, MEXICO.

## Abstract

In this work, we study the mechanisms of classical activation and inactivation of signal transduction by the histamine H3 receptor, a 7-helix transmembrane bundle G-Protein Coupled Receptor through long-time-scale molecular dynamics simulations of the receptor embedded in a hydrated double layer of dipalmitoyl phosphatidyl choline, a zwitterionic poly-saturated ordered lipid. Three systems were prepared: the apo receptor, representing the constitutively active receptor; and two holo-receptors -the receptor coupled to the antagonist/inverse agonist ciproxifan and representing the inactive state of the receptor, and the receptor coupled to the endogenous agonist histamine and representing the active state of the receptor.

An extensive analysis of the simulation shows that the three states of H3R present significant structural and dynamical differences, as well as a complex behavior given that the measured properties interact in multiple and inter-dependent ways. In addition, the simulations describe an unexpected escape of histamine from the orthosteric binding site, in agreement with the experimental modest affinities and rapid off-rates of agonists.

## INTRODUCTION

The G-protein coupled receptors (GPCR) is the largest family of integral-membrane signaling receptors and their topology consists of a bundle of seven transmembrane α-helices (TM1-TM7). The N-terminus (N-ter) is extracellular. Since the number of TM helices is uneven, the C-terminus (C-ter) is intracellular. Connecting the helices are three extracellular loops (EC1-ECL3) and three intracellular loops (ICL1-ICL3). The GPCRs belong to the superfamily of 7TM receptors.^1–3^ The histamine receptors (H1R, H2R, H3R and H4R) are members of the biogenic amine receptor subfamily of GPCRs and present, in addition to the TMs, a short C-ter amphipathic juxta-membrane helix 8 (H8) in the cytoplasmic side, oriented parallel to the membrane and making contacts with this one. The histamine receptors’ interaction with histamine (HSM; Fig. 1a), a neurotransmitter, elicits a variety of physiological effects, including allergic reactions (H1R),^4^ gastric acid secretion (H2R),^5^ mediation of neurotransmitter release and the inhibition of cAMP production (H3R),^6, 7^ and immunological response (H4R).^8^ HSM, the endogenous agonist of this family of receptors is a neurotransmitter synthetized and released by histaminergic neurons. It plays a major role in cognition and in other physiological functions such as vigilance, attention, impulsivity and feeding/weight regulation.^9^ It is stocked in vesicles and released after an electrical stimulus. HSM will bind pre- or post-synaptic receptors. Ciproxifan^10–12^ (CPX or FUB-359; CAS No. 184025-18-1; GRAC database, guidetopharmacology.org; Fig. 1b) is a highly potent and selective competitive H3-receptor antagonist/inverse agonist with pro-cognitive properties and a nanomolar affinity (for a given signaling assay used, inverse agonism refers to the ability of a compound to inhibit constitutive GPCR signaling, presenting thus negative efficacy).^13^

**Fig. 1.**
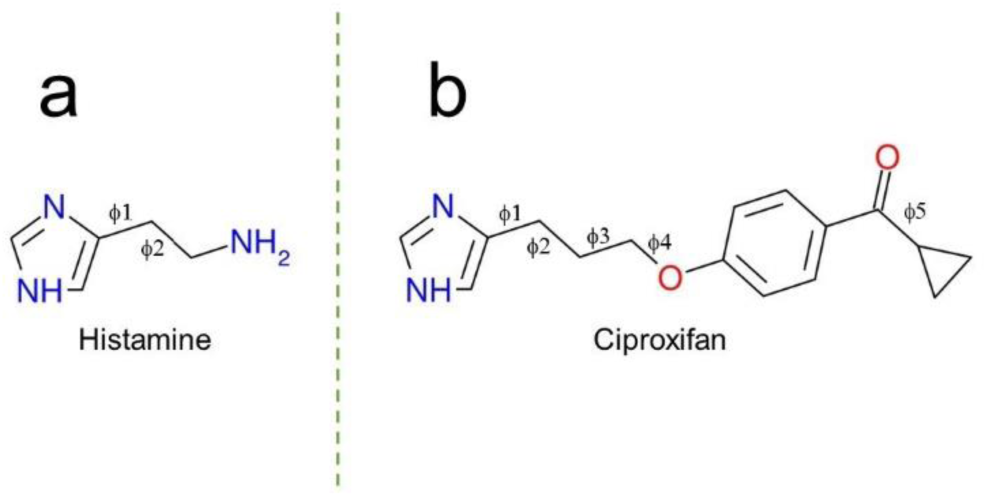
Chemical Structure of a) histamine and b) ciproxifan, with torsions around single bonds denoted ϕ.

The rat H3R (rH3R) was originally discovered in the brain on histaminergic neurons as a presynaptic auto-receptor and heteroreceptor inhibiting the synthesis and the depolarization-induced release of HSM.^7^ H3R is predominantly expressed in the central nervous system (CNS) and to a lesser extent in the peripheral nervous system.^14^

The cloning of the histamine H3R cDNA in 1999^6^ allowed detailed studies of its molecular aspects and indicated that H3R can activate several signal transduction pathways. The H3R is regarded as a potential therapeutic target because of its location in the CNS and for the modulation of a variety of functions such as cognitive processes, epilepsy, food intake and sleep-wakefulness.^15, 16^ The transmembrane region of H3R is often the site of ligand and drug interaction. Several H3R antagonists/inverse agonists appear to be promising drug candidates.^17–19^ Three-dimensional (3D) atomistic models of antagonist-receptor complexes have been used to investigate the details of ligand and drug interactions with H3R and have been successful in providing important insights regarding their binding; additionally, several groups have reported the features of the general H3R pharmacophore. This approach has been particularly successful for investigating GPCR/ligand binding modes and is complementary to 3D receptor/ligand modeling. The features of this antagonist pharmacophore are a primary basic group, either a piperidine or pyrrolidine, which is connected by an alkyl linkage to a second group; other groups observed that the addition of a second basic group increases the binding affinity.^20^ Across the superfamily of GPCRs, there exist many residues that have been conserved throughout evolution and are thus thought to play key roles in receptor structure and/or function. Site-directed mutagenesis has demonstrated the importance of many of these residues in several different biogenic amine receptors, including some of the HSM receptors^21^ (and references therein). Human H3R is sensitive to monovalent cations such as sodium.^22^ The interaction of the ligand with Asp 2.50 facilitates binding to residues in other TM domains. The critical role of TM5 has been demonstrated in many receptors (β2-adrenergic, H1, H2, etc.) including the H3R.^23^ On another hand, the existence of distinct active and inactive conformations of the H3R has been established *in vitro* and *in* vivo via the pharmacological concept of protean agonism.^24, 25^

In the absence of the experimental structure of H3R, several computational studies have been carried out for determination of the binding of several antagonists to H3R using a homology model for the receptor and employing the continuum dielectric approximation for the surrounding bilayer environment. Thus, based on the crystals of rhodopsin,^26^ 3D *in silico* models for H3R have been obtained in the past.^20, 23, 27–37^ Nevertheless, these models suffer from a variety of imperfections during their building, such as use of incorrect or absent alignments; manual adjustments and manipulations; energy refinement *in vacuo*; short molecular dynamics (MD) trajectories (1-10 ns); non consideration of the state of the template structure (active/inactive). Other models do not contain any ligands.^38^ In other instances, modeling of GPCRs (class A, B and C) was performed on a high-throughput basis, but the models only pretended to represent TM domains of receptor structures aimed at studying receptor-antagonist interactions and not their activated states.^39^ In the MemProtMD database,^40^ the structures of membrane proteins in the PDB are inserted in an explicit lipid bilayer and MD simulations in a coarse-grained (CG) representation are undertaken. But the structures are taken from the PDB as they are, with missing loops and/or segments, so that the reported analyses (contacts, displacements) might not be accurate. As expected, significant divergences among the models exist.

In this work, we generated curated *in silico* 3D structural models of H3R in the active, inactive and constitutive states, this latter being represented by a ligand-devoid receptor in an active state. We then proceeded to embed the receptor models in a hydrated, ionized and electrically neutral DPPC phospholipid bilayer for studying its behavior. Our approach consisted in simulating the binding of known H3 ligand compounds to the receptor model for studying and analyzing the spatiotemporal behavior of the resulting H3R-ligand complexes through MD simulations totaling more than 3 µs. An extensive analysis of the results of the trajectories indicated that each state of the receptor is different and is described by many structural and dynamical properties that are interdependent, showing an intricate network of short- and long-distance crosstalk between distinct regions of the receptor. This is a fingerprint of a complex behavior st*ricto sensu*. In addition, the MD simulations showed a spontaneous escape of HSM from the orthosteric binding site, preceded by a short binding step in the extracellular vestibule during the unbinding pathway.

In establishing the signaling mechanism of H3R, identifying the large-scale movements taking place, the structural changes conveying from the ligand binding pocket to the G protein binding site, the interaction networks taking place and how those changes propagate, is of utmost importance and should help to develop efficient effectors of this receptor.

## RESULTS

### Free energy landscapes suggest a multiplicity of conformations for each state of the receptor and between receptor states

Fig. 2a shows the PC1-PC2 2D plot of the cartesian Principal Component Analysis (PCA) obtained for the protein structure of the **antagonist-H3R** complex. In the presence of CPX, the plot shows a V-shaped energy landscape with the vortex to the left and with four populations: clusters C1, C2, C3 and C4, located in quadrants 1 to 4 (Q1-Q4). The V-shape shows a large span of about 65 Å over PC1 and a smaller one of about 43 Å over PC2. Fig. 2b shows the PC1-PC2 plot for the structure of H3R in the **agonist-H3R** complex when the agonist is still bound to H3R. It is composed of three major clusters -C1, C2 and C3- and a minor one (C4 in Q3). The PC1-PC2 plot for the **apo** receptor is in Fig. 2c. The occupied region in the map is rather large and diffuse, presenting several energy clusters with a marked cluster at the Q1/Q4 border (C2). Other energy maps (PC1 vs. PC3, PC2 vs. PC3) are in the SI section.

**Fig. 2.**
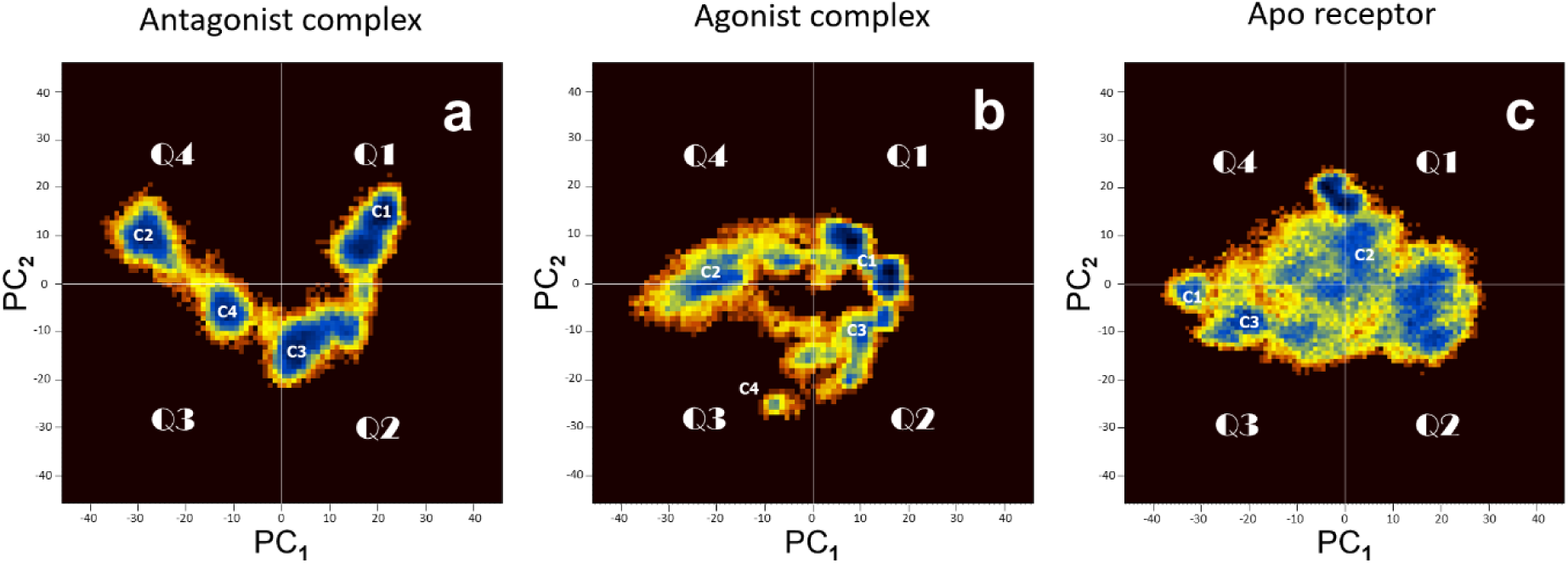
PCA-based free energy landscape of the structures of the three systems identified by cartesian Principal Component Analysis. The panels a), b) and c) show a pseudo color representation of the distribution of the first two principal components PC1, PC2 obtained from a ∼ 1 µs MD simulation for the antagonist, agonist and apo structures, respectively. Blue color represents energy wells (C1, C2, etc.). The map is divided into four quadrants (Q1-Q4).

Even though the principal component maps cannot be interpreted directly in physical terms, it is noteworthy to observe that, in general, the maps for the different systems show distinct morphologies, and number and depths of the conformation populations. This is due solely to the presence or absence of ligand -holo or apo- and to its nature –agonist or antagonist and reflects the conformational heterogeneity taking place for each given state of the receptor. In addition, the PCA clusters are structurally consistent with the metastable conformational states evidenced through the RMSD matrices (SI).

### Eigenvectors generated by the PCA show intra-molecular correlated movements and inter-molecular differential movements

The eigenvectors represent the global displacement of each residue in the system for all the trajectory and are used to generate new conformations. The images in Fig. 3 show the superposition of these conformations for each of the four clusters (C1-C4) generated by the PCA for the **antagonist-H3R** complex. The ensemble of conformations in Fig. 3a corresponds to cluster 1 (C1) and shows that this mode involves essentially large movements of the N-ter extracellular head of the receptor and its ECL2, which act as a flexible lid that opens and closes on top of the ligand-binding cavity. Loops ICL2 and ECL2 exert a push-n-pull motion on TM4 as a rigid body. For TM5, we observe a compression-extension movement of the fragment above Pro 5.50 and a kink movement around this residue. We also observe kink movements around Pro 6.50 and Pro 7.50 of the conserved ^6.47^CWXP^6.50^ and ^7.49^NPXXY^7.53^ motifs, respectively. All these movements are correlated. No other important movements are detected in the other helices. C2 in Fig. 3b shows large correlated movements of ECL2, ICL2 and to a lesser extent ICL3, with slight kink motions of TM1, TM5-TM7 around their middle-helix proline residues. In addition, TM5-TM7 move collectively in a breathing motion along the plane of the membrane. The mode corresponding to C3 shows large amplitude motions for ECL2, and a pendulum-like rigid-body motion of the lower zone of TM5-7 with their IC loops (Fig. 3c). Fig. 3d shows the collective modes of movement corresponding to the last cluster, C4. First is an angular N-ter to C-ter motion of H8, whose C-ter gets close and far from the membrane, accompanied by a concerted motion of the N-ter, ECL3 and ICL2, indicating long-range communication between the extra- and intra-cellular regions of the receptor, as in activation of the µ-opioid receptor.^41^ With regards to the **agonist-H3R** complex, Fig. 4a shows the conformations of C1. We can observe big correlated motions of the N-ter and ECL2 extra-cellular segments of the receptor. Along with large conformational fluctuations in ICL2, this latter motion leads to an opening and closing of the far-away intra-cellular domain of the receptor, in agreement with experimental 3D structures of active state GPCRs, such as β2-AR.^42–45^ Fig. 4bcd show essentially motions of the N-ter of the receptor. Fig. 5a shows the eigenvectors of C1 for the **apo** receptor. The N-ter, ECL2 and ECL3 undergo large concerted unfolding-folding motions, with ECL2 exerting a hinge-bending motion on TM4 perpendicular to the plane of the membrane. For C2 (Fig. 5b), the motions are much reduced. For C3 (Fig. 5c), large motions of N-ter and ECL2 are observed, accompanied of an important movement of the ICL3 loop. The large motions detected in C1 and C3 may be attributed to the absence of a bound ligand.

**Fig. 3.**
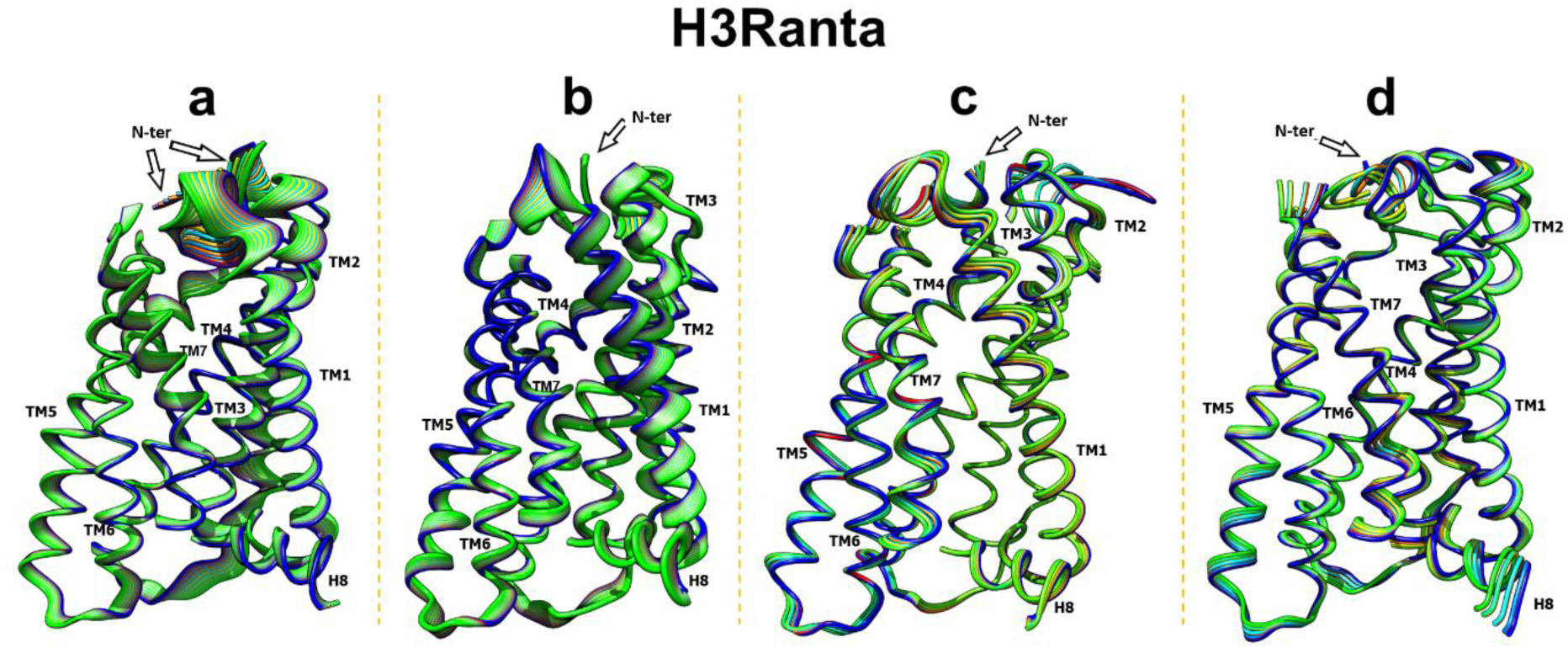
Clustering of the antagonist state structures. Visualization of the movements of the first principal component PC1 in each of the four clusters, from green to blue, depicting low to high atomic displacements. Panels a) cluster C1, b) cluster C2, c) cluster C3, and d) cluster C4 as determined with the Carma package.

**Fig. 4.**
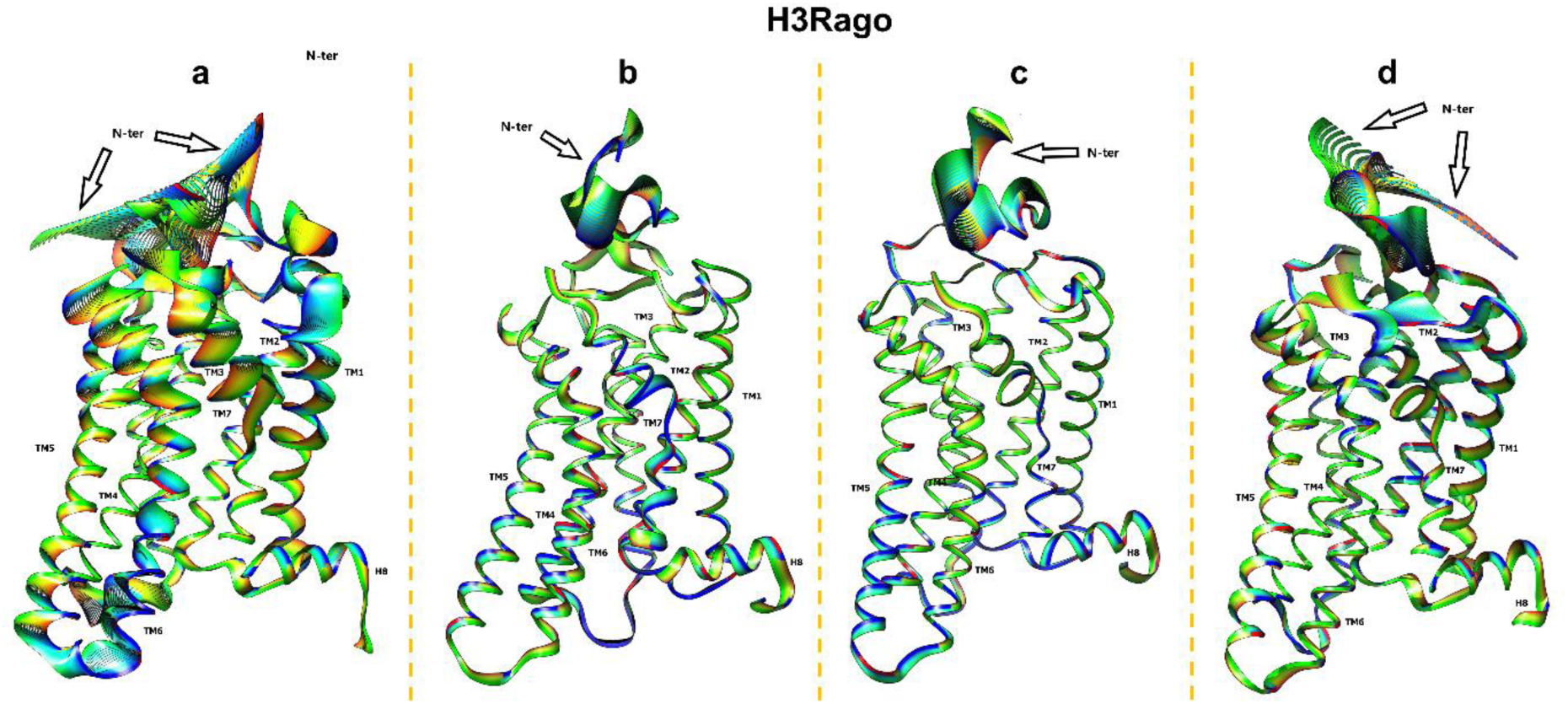
Clustering of the agonist state structures. Visualization of the movements of the first principal component PC1 in the four clusters. Color scale as in Fig. 3. Panels a) cluster C1, b) cluster C2, c) cluster C3, and d) cluster C4 as determined with the Carma package.

**Fig. 5.**
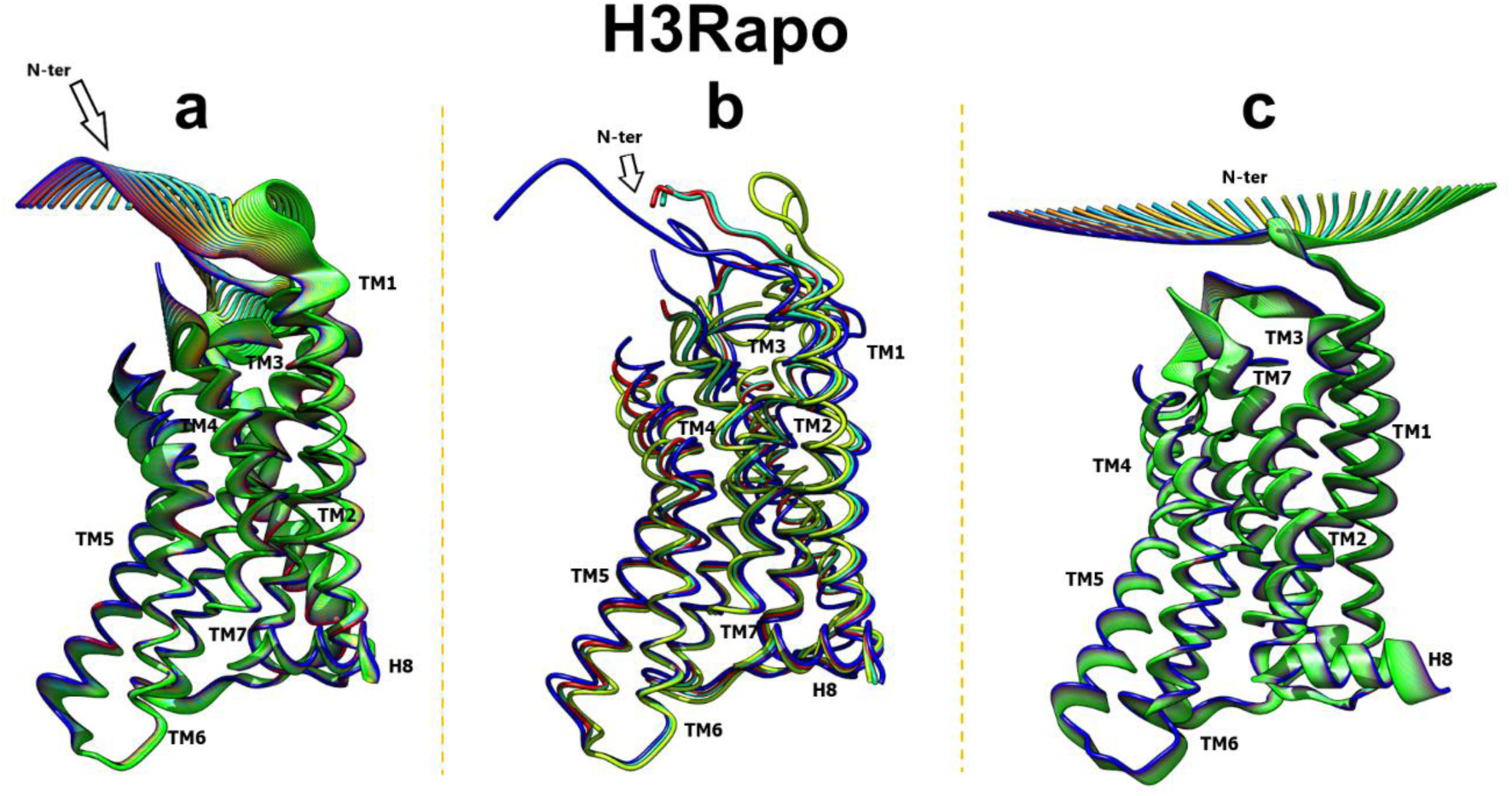
Clustering of the apo state structure. Visualization of the movements of the first principal component PC1 of the three clusters C1, C2 and C3 as determined by the Carma package. Color scale as in Fig. 3. Panels, a) cluster C1, b) cluster C2, and c) cluster C3.

Therefore, the differences in morphology of the PCA maps and clusters of the holo-receptor (Fig. 2ab, 3, and 4) with respect to those of the apo receptor (Fig. 2c and 5) suggest that receptor conformations are ligand induced.

### Agonist and inverse agonist establish differential interactions with the receptor

For **CPX**, the carbonyl oxygen between the cyclopropane and the phenoxy ring is in a long-lasting (>70% of residence time) bifurcated bond with the N atoms of the main chains of Glu 45.53 and Phe 45.54 (ECL2) (Fig. 6a). Whereas, due to the mobility that the single bonds give to the imidazole ring, this one interacts at one time or another with either the internal solvent molecules or different neighboring residues; none of these interactions is significantly populated. As shown in the LigPlot+ diagram^46^ of Fig. 6a, the imidazole moiety is associated to three water molecules. In Fig. 6b, this moiety is H-bonded to Asp 3.32 and a water molecule, whereas in Fig. 6cdef, the imidazole ring shows no H-bonds to water molecules. In Fig. 6e, the imidazole is H-bonded to the N3 of the indole of Trp 7.43. Other contacts along the trajectory are listed in Table 1 and include all residues within 4 Å of the CPX ligand with large residence times. The residues with 70% or more in contact with CPX include three Cys, two Phe and one of each Leu, Ala, Val, Tyr, Trp, Asp, Glu, and Gly. Notice the presence of two acidic residues - Asp and Glu.

**Fig. 6.**
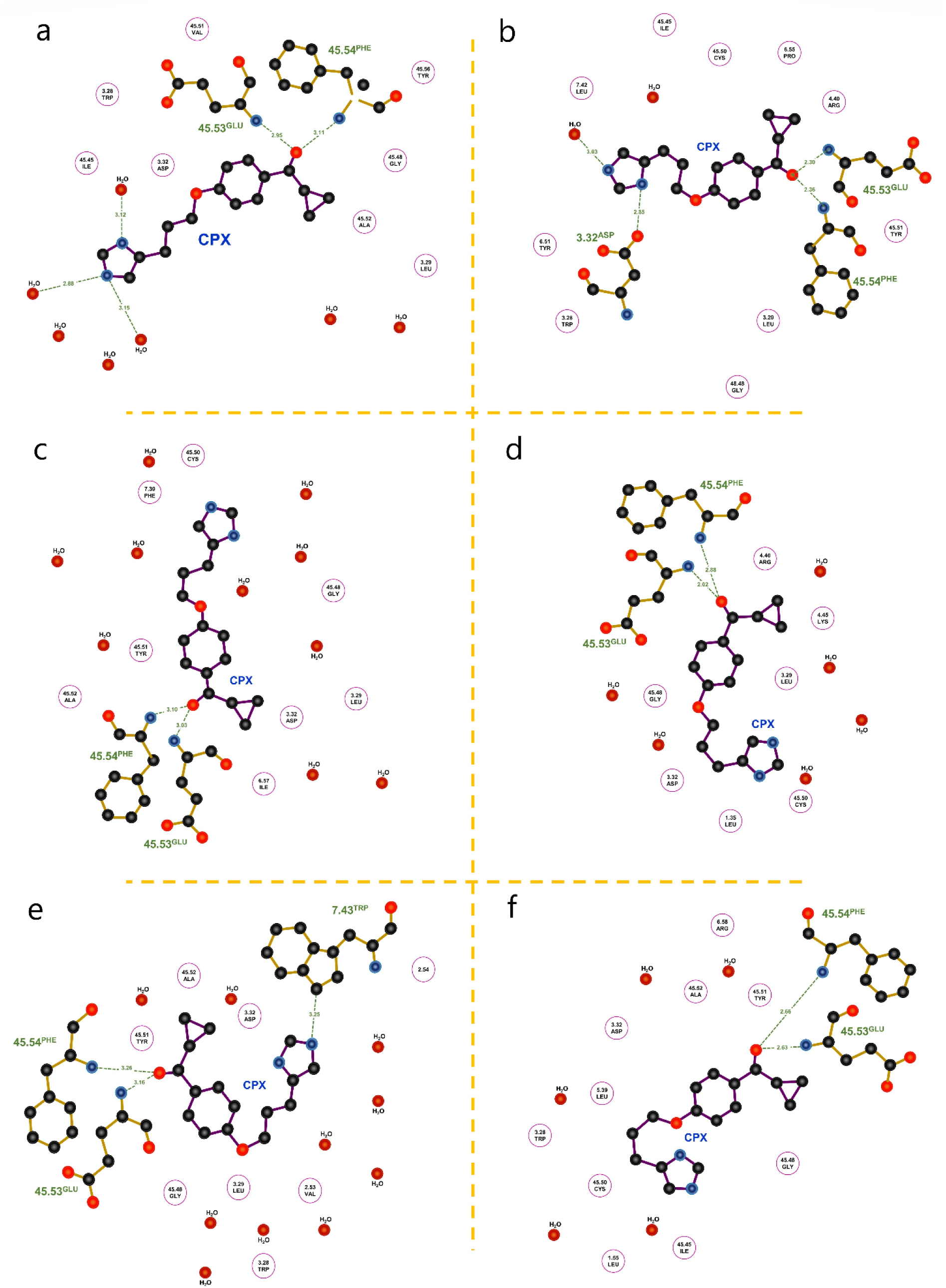
2D-binding mode of ciproxifan within the H3 receptor. Polar interactions and H-bonds (green lines) between the amino acids (in circles), water (red dots) and ciproxifan ligand (CPX) can be appreciated. The a) to f) panels show the most relevant configurations and bioconformations of CPX throughout the simulation. For torsion angles ϕ1-ϕ4, the bioconformations include: a) an extended or all-*trans* chain; b) a “zig-zag” chain; c) a “paddle” chain; d) a “cyclic” conformation that approaches Cδ2 of the imidazole with C11 (the carbon bound to the ether oxygen), forming a virtual 5-membered ring; e) and f) two “closed” or *cis* conformations. In the last frame of the trajectory, CPX adopts the conformation in a). The dihedral connecting the carbonyl oxygen and the cyclopropane, ϕ5, adopts two states: perpendicular (−90 to −110°) or co-planar (±180°) to the aromatic ring.

**Table 1.**
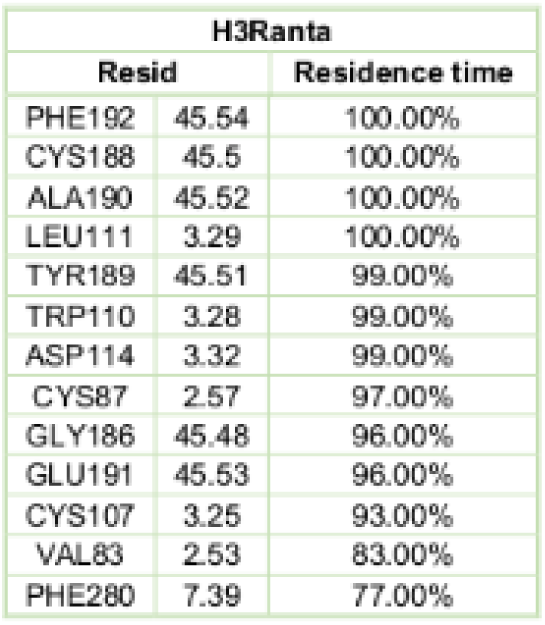
Interactions between the ciproxifan antagonist/inverse agonist and the receptor for residence times greater than 70%.

Thus, the LigPlot+ plots in Fig. 6 show the flexibility of the imidazole-containing moiety of CPX, reflected in the wide range of values the different single-bond torsion angles adopt during the MD trajectory (t, g+, g-, −120°), leading to essentially six bioactive conformations (a-f).

For the endogenous ligand **histamine**, the number of residues most in contact are Asp 3.32, Trp 3.28 and Trp7.43, and Phe 45.54 (ECL2) (Table 2; Fig. 7, section 1). Fig. 8b shows an HSM binding mode corresponding to section 1. The binding pocket of the apo receptor, along with internal water molecules is in Fig. 8c.

**Fig. 7.**
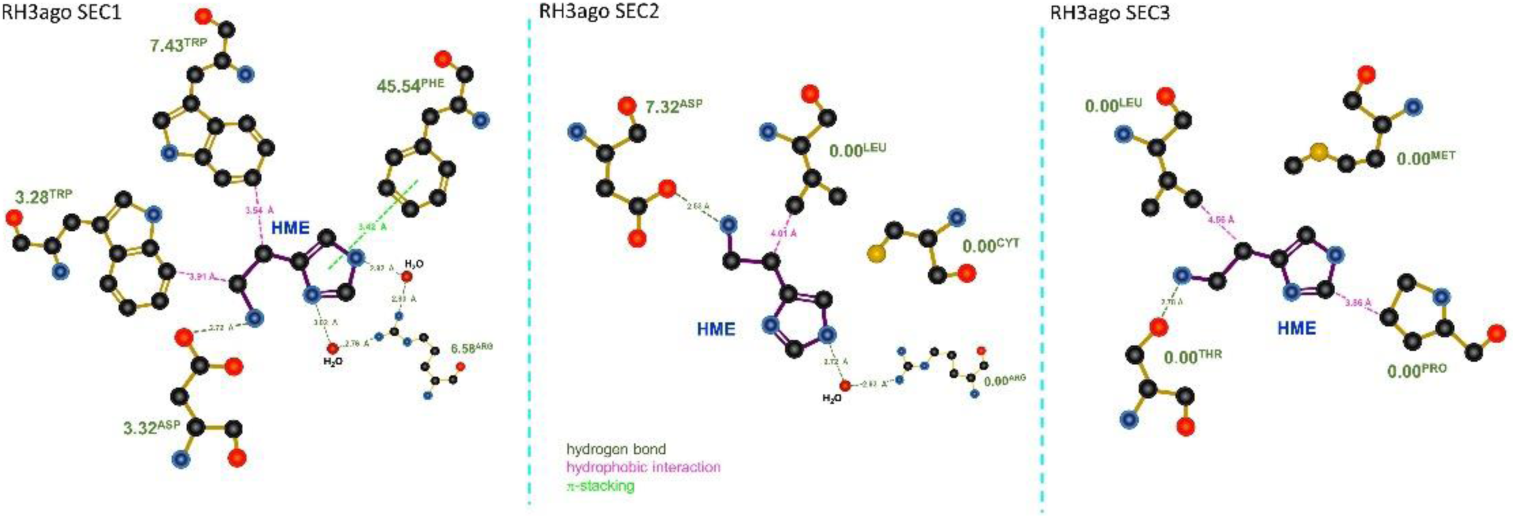
2D-binding mode of histamine within the H3 receptor. Polar interactions and H-bonds between the amino acids, water and histamine ligand (green lines); hydrophobic interactions (magenta lines); and π staking (light green lines) can be appreciated. The three panels correspond to sections 1-3 in the trajectory and show the most relevant configurations and conformations of histamine throughout the simulation.

**Fig. 8.**
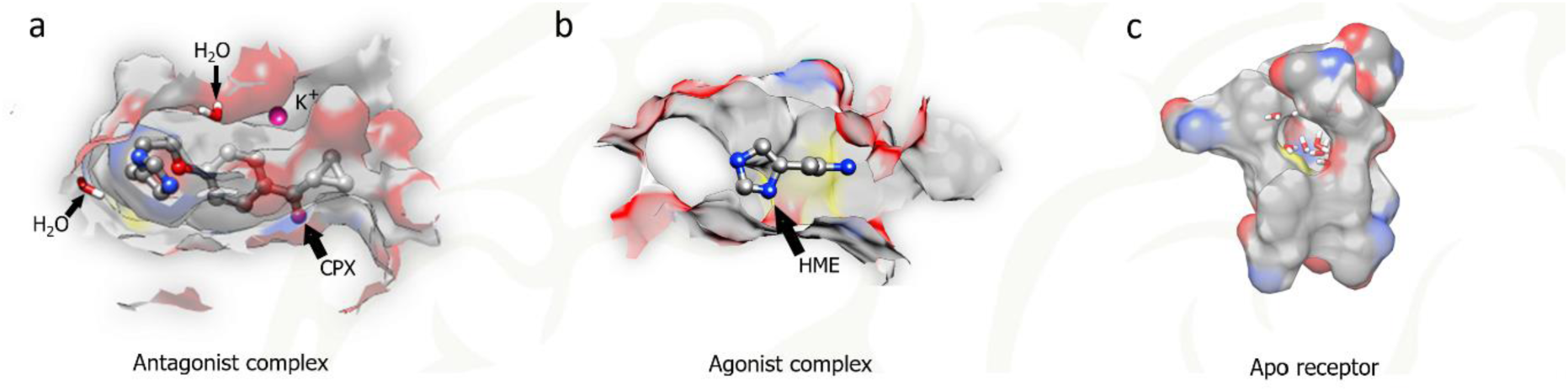
Side view of the H3R ligand binding pocket for a) the antagonist complex, b) the agonist complex and c) the apo receptor (devoid of ligand), respectively, as resulting from an optimal structural alignment of the TM helices of each system. The average number of water molecules in the orthosteric site is of 61, 35, and 49, respectively. Binding pose f (Fig. 6f) is shown in a), as well as the K^+^ ion.

**Table 2.**
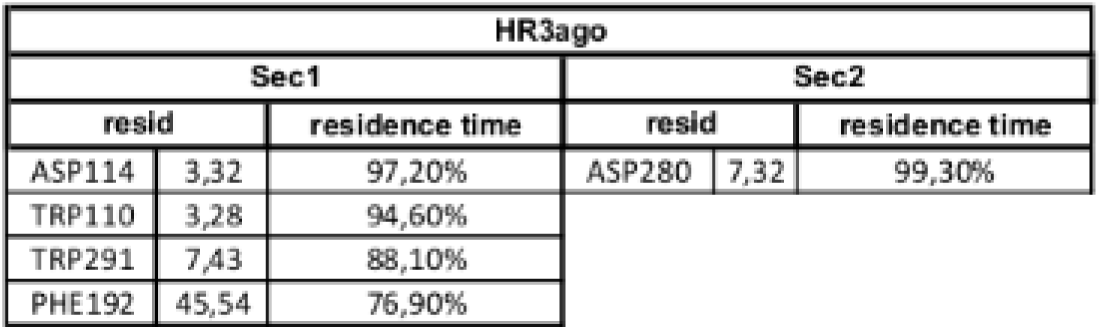
Interactions between the histamine agonist and the receptor for the first two periods of the production trajectory. Residence times greater than 70%.

As for the effects of the Glu 5.46 to Ala mutation,^23^ instead of a long-lasting direct interaction between the imidazole ring of either ligand and Glu 5.46, we suggest rather an indirect/allosteric interaction involving the sodium allosteric binding site.

### K^+^-dependent conformational changes: diffusion into the antagonist complex sodium allosteric site

During the production run, a potassium monocation from the bulk solvent spontaneously diffused into the sodium allosteric pocket of the antagonist-H3R complex, near Asp2.50. As the ion appears only in an extracellular vestibule of the receptor and then in the cavity, the diffusion of the ion is extremely fast (∼100 fs; Fig. S47). The cation is hydrated by five water molecules while in the aqueous milieu, three during its path to the interior of the receptor, and five in the final stage, when in the Na^+^ allosteric site. The presence of the potassium cation in the antagonist-H3R complex argues for a role equivalent to that of the sodium cation in the stabilization of the inactive state.^47^ By analogy, we assume that a Na^+^ follows the same insertion pathway as the K^+^.

As far as HSM is concerned, if there is no hysteresis, i.e., the HSM entrance and exit pathways are the same. On another hand, the HSM and K^+^ pathways are different (Fig. 9 vs. Fig. S47).

**Fig. 9.**
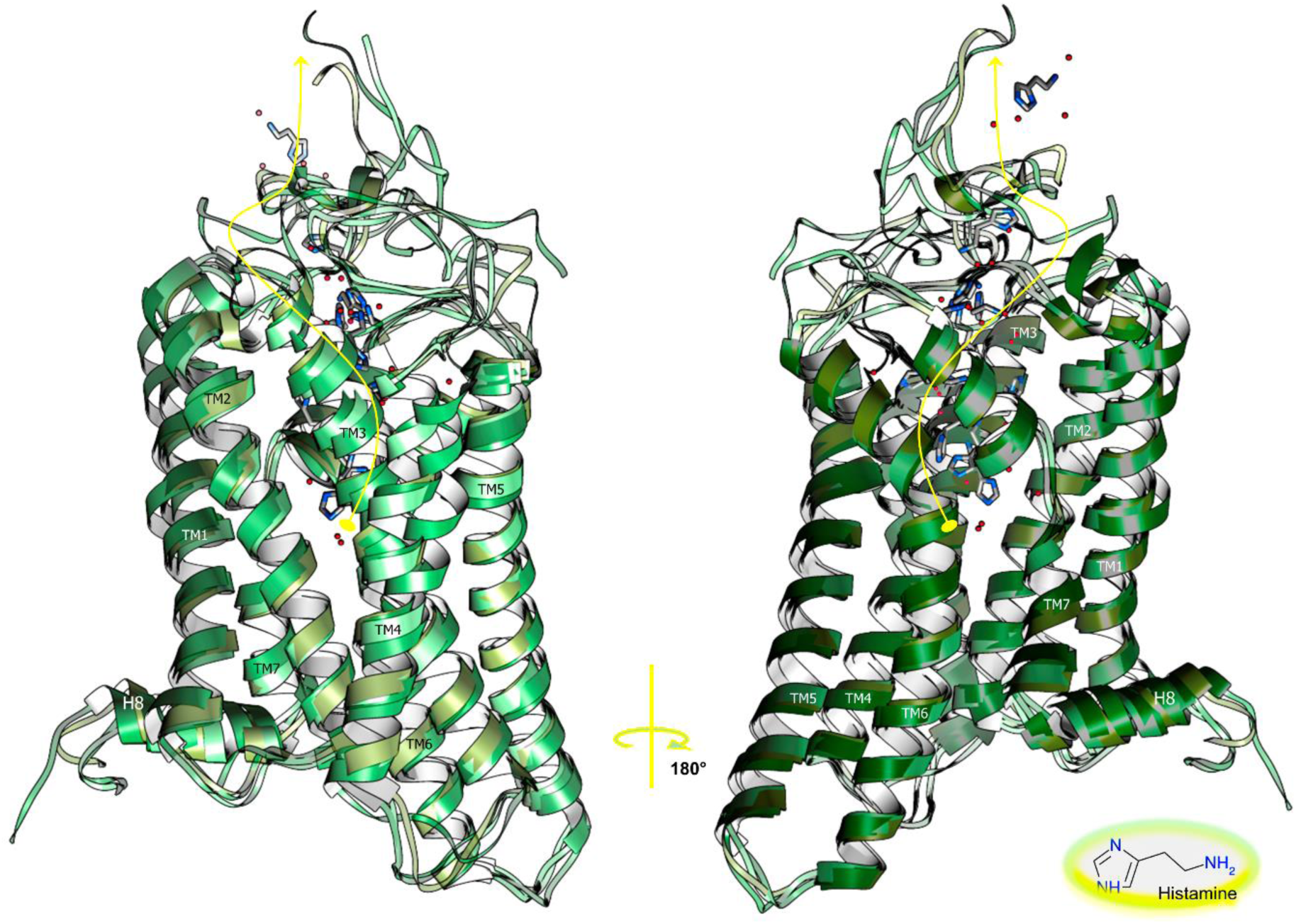
Schematic illustration of the four phases of the spontaneous unbinding and exit pathway of the HSM ligand (curve in yellow), showing its lodging in the binding cavity (section 1), with interactions of either the imidazol or the amino group with Asp 2.50, Leu 2.60, Tyr 2.61, Trp 23.50 (ECL1), Trp 3.28, Asp 3.32, Tyr 3.33, Ser 3.39, Phe 45.54 (ECL2), Tyr 6.51, Trp 6.48, Leu 6.54, Met 6.55, Arg 6.58, Tyr 7.35, Trp 7.43 and Ser 7.46; its disposal from it (section 2) with interactions with Ser13, Gly14, Ala15, Gly25, Ala26, Arg27, Gly28, Phe29, Ser30 of the N-ter, Trp 3.28, Leu 3.29; Ala 45.52, Glu 45.53, Phe 45.54 of ECL2, Arg 6.58, Asp 7.32, Tyr 7.35, Glu 7.36 and Trp 7.43; its small residence time in the vestibular area formed by Met1, Asn11, Ala12, Ala15, Arg27, Trp33 of the N-ter, Trp 3.28, Glu 45.53 (ECL2) and Asp7.32 (section 3); and its escape into the solvent (section 4). Several of the previous interactions are mediated by water.

Finally, Arrang and co-workers^48^ observed that the presence of Ca^2+^ down-regulated the H3R receptor. We attribute this effect to an allosteric binding site for Ca^2+^ at the extra-cytoplasmic region and/or at the sodium binding site (their ionic radius −114 and 116 pm, respectively-are very similar).

### CPX forms a stable complex in the orthosteric site, but HSM shows several binding modes and eventually quits the site spontaneously in a short time interval

We obtained the initial structures for the two holo-systems, i.e., the HSM-H3R complex and the CPX-H3R complex through ligand docking to the orthosteric site of the receptor.

We found that there were multiple non-covalently bound states of HSM corresponding to the activated state of H3R due to the small size of the agonist and to its ability to establish stabilizing non-bonded interactions in distinctly different binding orientations with surrounding residues. Indeed, during the docking simulation HSM showed several major binding modes in several binding subpockets in the orthosteric site. For the starting structure, we chose the one compatible with experimental results with H2R in which the quaternary Nς of HSM interacts with Asp 3.32 (Asp98 of H2R; Asp114 of H3R), highly conserved through class A GPCRs and essential for HSM binding and action, serving as a counter-anion to the cationic amine moiety of HSM and agonist and antagonist binding.^49, 50^ The found multiple poses are compatible with the experimental findings of Gantz et al.^49^ in which removal of the negatively charged amino acid abolished HSM stimulated increases in cellular cAMP, but not HSM binding to the receptor, supporting HSM’s multiple binding poses, but suggesting that the one in which it interacts with Asp3.32 leads to activation of the receptor. For CPX, we chose a binding pose analogous to that of HSM in which the imidazole moiety is in contact with Asp 3.32. The binding pockets of the two ligands are thus partially shared. There may be a secondary allosteric low-affinity binding site for agonists as seen by binding kinetics,^48, 51^ but in this study we focus on the orthosteric site only.

Interestingly, during the MD simulation the agonist-H3R complex showed an unexpected behavior in which the HSM ligand, even though at given times deep in the binding cleft, eventually escaped spontaneously from the binding site. We thus defined four intervals to better describe the unbinding process. The first interval (section 1) during which the ligand is bound to the orthosteric binding cavity, comprises from nanosecond 112 of the MD production phase, when the RMSD of Cα atoms attains a plateau, to nanosecond 590. In this interval, the HSM displays, nevertheless, a very high mobility despite the solvent molecules filling the cleft. The N-ter folds over ECL2, results in a salt bridge between Arg27 and Glu 45.53 (ECL2), respectively, with ECL2 folding over the cleft, capping the binding pocket. The second interval (section 2), a transition, includes the beginning of the ligand out of the site but still bound to the transmembrane part of the receptor-it lasts 1.15 ns. The N-ter begins then to unfold and ECL2 moves aside following the disruption of the salt bridge. The third interval (section 3) comprises the positively charged ligand trapped between the extra-membrane N-ter and ECL2 regions and the anionic heads of the upper leaflet of the phospholipids; it lasts almost 1 ns. Thus, it involves the extracellular vestibule, but it does not seem to represent a metastable binding site.^52^ The binding cleft is then open to the extra-cellular space. The fourth and last interval (section 4; 592 ns − 1.02 µs) in the unbinding pathway defines the exit of the ligand to join the bulk of the solvent. The N-ter now forms the lid of the cleft, with ECL2 remaining “aside”. In the bulk, HSM shows a hydration shell of about 30 water molecules and interacts eventually with a chloride ion with its alkyl amine. HSM binds from time to time to the extracellular vestibule. The exit pathway of HSM is illustrated in Fig. 9 and involves TM2, ECL1 TM3 ECL2 TM6 and TM7 for section 1; N-ter, TM3, ECL2, TM6 TM7 for section 2; N-ter, TM3 ECL2 and TM7 for section 3; the solvent for section 4.

## DISCUSSION and CONCLUSIONS

GPCR dynamics are important from a structural and functional viewpoint, including conformational changes and rearrangements of the transmembrane helices.^53^ In this work we have explored with extensive MD simulations many microscopic spatial and temporal properties of the classical activation and inactivation pathways of the second step (see below) of signal transduction by the H3 receptor that are not easily accessible by experiment, including new undocumented interactions. Of course, other activation mechanisms exist.^54^

The average values of several indicative properties, such as the RMSD and the RMSD matrix (Fig. S1, S2), remain roughly constant with increased sampling, even though the production run temperature was relatively high (323.15 K), leading us to believe that the 1 µs MD simulations for each of the three systems sample satisfactorily the phase space.^55^

Furthermore, we ran two other parallel small trajectories that pointed to the general character of our observations (not shown). An apparent paradox is that we can observe significant phenomena during MD simulations that span 1 µs, whereas the time constant for the activation switch of a GPCR can be in the tens to hundreds of ms. For example, for α2A-AR in living cells the rate constant was < 40 ms;^56^ these authors also refer to other results that indicate that the <u>entire</u> GPCR-signaling chain can be activated within 200-500 ms. The time-scale difference between theory and experiment is due, in our view, to the fact that we have studied only a confined time interval of the entire GPCR signaling cascade whose individual elements comprise at least (1) ligand binding, (2) conformational change of the receptor (preliminary activation or inactivation), (3) interaction between the ligand-complex receptor and the G-protein (full activation), (4) G-protein conformational changes including GDP release and GTP binding, (5) G protein-effector interaction, (6) change in effector activity, and (7) the resulting ion conductance or second messenger concentration changes.^57^ Steps 1-3 are dovetails with the ternary complex model for GPCRs^58, 59^ and correspond to the Ri ⇔ ARi ⇔ ARa schemes of the extended or cubic complex models (Ri, initial state unbound receptor; ARi, ligand-complexed receptor in the initial state; ARa, ligand-complexed receptor in the active/inactive state). We may add receptor oligomerization between steps 2 and 3 when applicable. But our simulations do not deal with the whole activation process, rather with step 2 only-the initial structures represent already the docked complexes- and can be considered to span it well, even though the H3R in any of the two ligand-bound described states is not in a fully active or inactive state, rather a “pre-active” state since it is not bound to its G-protein (Gi/o) to form the ternary complex, but in which the receptor is able to interact with it.^60, 61^

An illustration of this second step in which there is conformational fluctuation of the receptor due to ligand binding is the case of β2-AR, whose agonist-induced conformational changes include a large outward movement at the cytoplasmic end of TM6 and an α-helical extension of the cytoplasmic end of TM5.^43, 44^ This is observed also in the collective motions of the TM5, 6 and 7 helices of H3R (Fig. 10), supporting the view that several receptor collective movements are common throughout class A GPCRs. In addition, we can observe in the extracellular face view of Fig. 10 that the next exposed TM helix TM5 is that of H3R in the apo state, whereas the inner most TM5 is that of the antagonist complex. Instead, peripheral TM4 is more eccentric in the order antagonist complex > apo receptor > agonist complex.

**Fig. 10.**
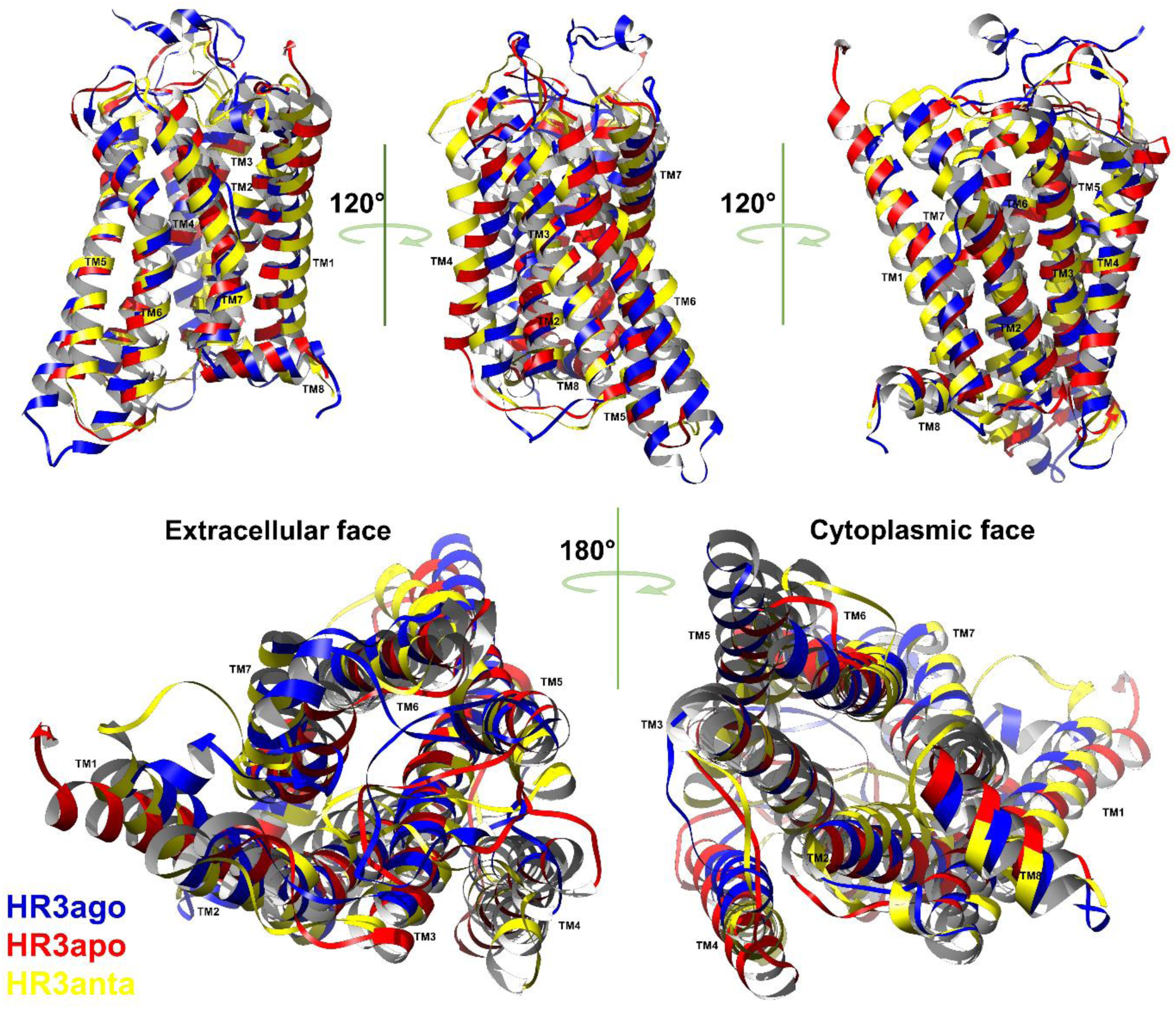
3D superposition of the production-phase average representative structure of each system. In yellow, the antagonist-H3R complex; in blue the agonist-H3R complex; in red the apo receptor.

Ghamari et al.^62^ built a 3D model of human and rat H3R (with N-ter absent and with ICL3 modeled as the lysozyme fusion protein in the crystal structure) by the homology comparative approach using the M3 muscarinic acetylcholine receptor in complex with tiotropium, a potent muscarinic inverse agonist (PDB Code 4DAJ). Thus, the model, just like the template receptor, is in the inactivated state. For the structurally conserved segments of the molecule, i.e. the TM helices, the RMSD (as estimated by PyMOL’s Align function) with our CPX-bound H3R is of 3.2 Å, the main deviations located at TM4 and 5. Over the years, Stark, Sippl and coworkers have published several *in silico* 3D models of H3R for different purposes.^34, 63^ The 2001 publication, a study of receptor–antagonist interactions, presented the first models of rH3R and hH3R and was based on the rhodopsin structure. In the 2005 work,^63^ the rhodopsin-based homology model of hH3R focused only in the hH3R binding pocket suitable for virtual drug design. The 2007 homology model of hH3R was also based on the crystal structure of bovine rhodopsin and was simulated by small MD trajectories in a DPPC/water membrane mimic, in view of obtaining binding sites not overfitted to a given ligand. Diverse binding modes with different ligands were obtained by allowing side-chain rotamerism, but CPX was not among the ligands probed. Yet, we found several features in common with our rH3R binding site, including the presence of important residues in the binding pocket, such as Asp 3.32, Tyr 3.33 and Glu 5.46. Levoin et al.^29^ generated several hH3R models based on the crystal structure of hH1R (PDB Code 3RZE) and described four distinct binding modes out of the published models of H3R. A binding mode of our HSM agonist corresponds to their β-binding mode (perpendicular to the membrane plane and bridging Asp 3.32 (114) and Asp 2.50 (80)). Another of our HSM binding modes is represented by mode γ, implying an interaction between Asp 3.32 and the amine of the ligand. Nevertheless, the ligand can be found in head-to-tail or tail-to-head orientation. The binding mode of the CPX inverse agonist in our complex is oblique to the membrane plane and has some features with mode δ. Finally, Kong, Y. and Karplus, M.^64^ studied the signal transduction mechanism of rhodopsin by MD simulations of the high resolution, inactive structure in an explicit membrane environment. Even though we are dealing with H3R and not rhodopsin, and that the ligands are not the same, several of our results are analogous to those results, indicating common structural features in the general mechanism of class A GPCR activation.^65^ Thus, we observe correlated movements of TM6 and TM7 around the class A GPCR highly conserved Pro 6.50, and Pro 7.50 of the CWXP and NPXXY motifs. In agreement, Kong and Karplus find that the major signal-transduction pathway involves the interdigitating side chains of helices VI and VII.

Several new interactions are observed to contribute to the mechanism of signal propagation from the binding pocket to the G-protein binding sites in the cytoplasmic domain. Our results show that the biological state of the receptor is closely linked to its conformational dynamics and that the binding of a given ligand results in a variety of states, indicating the existence of ligand-induced heterogeneous receptor conformations, in coherence with experimental data.^25, 66–70^ Thus, as observed in this work for H3R, functionally different ligands induce and stabilize distinct conformations in the receptor (Fig. 11), as opposed to the existence in dynamic equilibrium of inactive- and active-state conformations. Indeed, the RMSD matrices and the free energy landscapes among others illustrate this phenomenon and show how subfamilies of conformations interconvert for a given state of the receptor.

**Fig. 11.**
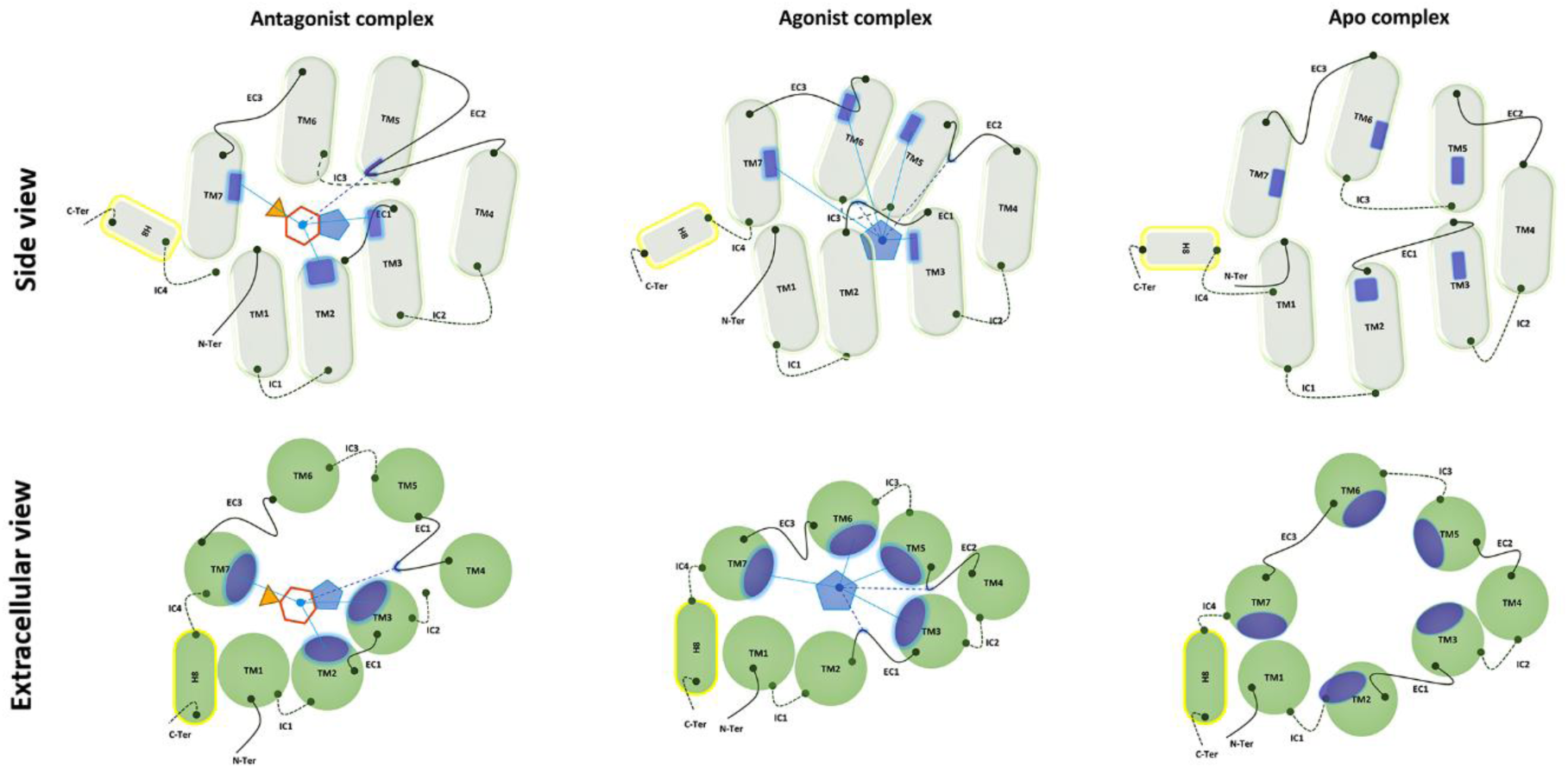
Schematic representation of the structural differences between the inactivated, the activated and the constitutive-activity H3R. The blue patches represent the contacts between CPX (multi-polygon), HSM (pentagon) and the TM α-helices (cylinders or circles). They show the relative displacement and rotation of the helices between the different states. Continuous curves at the helix extremes show the extracellular loops and the dotted curves the intracellular loops. Straight (dotted) lines represent interactions between the ligand and the α-helices (loops). HME binding carries with it a reduction of the size of the site. Approximate tilts in the TM helices are shown in the side view. The relative orientation of the receptor among the three systems is the result of an optimal superposition of the TM helices.

The present work confirms other findings reported in the literature and describes new phenomena that contribute to the mechanism of signal transduction: (i) diverse events leading to signal propagation from the binding pocket to the G-protein binding sites in the cytoplasmic domain upon binding by the endogenous agonist HSM, by the CPX inverse agonist, and in absence of ligand binding; (ii) specific lipid-binding sites on the receptor (Fig. S15, Tables S5-S7).

At constant number, the water molecules are very mobile in the orthosteric cavity. Notwithstanding, this number depends on the state of the receptor (Table S2, upper left-hand box), and of course on the size and the flexibility of the molecule bound to it. Several of these water molecules mediate H-bonds between the receptor and the ligands and play important roles not only in the activation processes, but also in inactivation and spontaneous activation (Fig. S10).

We made other non-reported compelling observations. The high mobility in the binding site of the small endogenous agonist ligand HSM leads to several poses and eventually to it leaving the binding site after a certain time lag (Fig. 9). Before escaping into the solvent, it interacts with the N-ter and ECL2, consistent with the reports that ECL2 seems to be involved in ligand binding, selectivity and activation of GPCRs,^71–73^ assuming the entry and exit pathways are similar. The pathway of ligand unbinding involves unfolding of the N-ter of the receptor. This very short-termed transition between the agonist-bound state and the constitutively activated state implies that the HSM-H3R complex is thus not a stable one and suggests that this orthosteric ligand can also act as an allosteric modulator. Experimentally, the agonist efficacy and the dissociation rate constant are highly correlated,^74^ explaining the high efficacy of the HSM endogenous agonist. The time interval of the unbinding of HSM, ∼2 ns, is very fast and fits together with the experimental rapid off-rates of agonists and modest affinities.^43, 51^ HSM binding is thus transient, and a short MD trajectory would not have witnessed this phenomenon. In contrast, the binding of the antagonist was stable all along the trajectory.

This dovetails with the experimental affinities measured for HSM and CPX, which indicate that the affinity of this latter for the receptor is at least an order of magnitude larger at the human H3R.^13^ The unbinding pathway observed for HSM in this simulation suggests a plausible binding pathway in which HSM would bind to an extracellular vestibule of the receptor,^52^ leading to transient activation, followed by “pre-activation,” in a stepwise binding mode as part of a sequential activation.^75^ Moreover, HSM bound to the extracellular vestibule of the receptor several times during its perambulation in the solvent bulk. And one binding mode was equivalent to the binding mode of section 3 of the getaway. Otherwise, the H3R orthosteric site experiences a significant plasticity since it can accommodate the conformations induced by functionally different ligands, in coherence with what is observed experimentally and computationally for many GPCR receptors.^61, 76^ Thus, the helices that are in contact with CPX are TM2, TM3 and TM7, as well as ECL2 (Fig. 6, Table 1); for HSM, the corresponding helices are TM3 and TM7, as well as the ECL2 (Fig. 7, Table 2).

Arrang and co-workers^13^ compared potencies of H3-receptor ligands on inhibition of [^125^I]-iodoproxyfan binding to rat H3R and human H3R. CPX displayed significantly higher potency at the rH3R when compared to the hH3R. Production of a partially humanized chimeric rH3R allowed them to identify two residues responsible for this heterogeneity: Ala 3.37 (119) and Val 3.40 (122), both from TM3. In our structures of the CPX-rH3R complex, those residues are not in a direct contact with CPX (Fig. 6), suggesting short-range allosteric effects behind the experimental observations and an increased, favorable hydrophobic environment for rH3R, as compared to hH3R showing instead Val and Thr at positions 3.37 and 3.40, respectively.

We have also seen that the lipid-binding sites in the receptor will depend on the state of the receptor. In other words, the location, number and structure of the lipid-binding site will recognize the state of the receptor. Tellingly, the lipids will bind to specific TM helices and will be localized in different regions-upper and lower leaflets-of the bilayer (Fig. S12-S14). Our analyses highlight also the presence of distinctly different lipid allosteric regulatory sites according to the state of the receptor^77^ and point to the mechanism of modulation of lipid on receptor function^78–80^-each specific protein-lipid interaction stabilizes the corresponding state of the receptor (Fig. S15). Therefore, just like cholesterol binding to the β2-AR, there are specific binding sites for DPPC in the H3R receptor, but specific to the state of the receptor.

Like palmitic acid and cholesterol in the β2-AR-carazolol crystal structure (PDB code 2RH1), the contacts between the lipids’ hydrocarbon chains and hydrophobic receptor residues follow the hydrophobic matching principle. But, as we found, the specific lipid-binding motifs on the membrane protein surface consist also of positively charged residues that specifically interact with the negatively charged phosphodiester groups (Fig. S12, S14). Our findings are thus consistent with the experimentally reported lipid binding sites in membrane proteins,^81, 82^ and with the protein-head and protein-tail contacts for the H1R (PDB ID 3RZE), as reported in the MemProtMD database.^40^

The complexity of the atomic phenomena involved in the structure and dynamics of H3R makes the design of ligands for the selective activation or inactivation of the receptor to produce the desired molecular and conformational effects difficult. Thus, the full amount of our results and the corresponding analyses lead us to believe that unraveling the mechanisms of signal transduction-activation, inactivation, constitutive activity-cannot be based on fluctuations in a single microscopic feature or a small number of them. These mechanisms are complex, as revealed in this work, and need multiple descriptors for a better understanding. The inter-dependent, epistatic descriptors involve rigid-body motions of helices, along with different variations in their mechanical properties (bend, wobble and winding-unwinding for TM5-TM7, Fig. S43-S45; tilt, rotation, turn angle per residue, Fig. S19-S42; helix displacement and shift, Fig. S46; multiple simultaneous rotamer toggle switches, Fig. S16-S18; ionic locks and inter-residue contacts, Table S10). We have observed variations as well in other physicochemical characteristics, such as the formation and rearrangement of networks of charged residues (Fig. S7-S9) and of H-bonds between amino acid residues (Table S1) and between amino acid residues and internal water molecules (Fig. S10); the differential composition in amino acid residues of the internal cavity of the receptor (Table S2); the presence of mobile networks of internal water molecules and their numbers; the presence of hydrophobic and aromatic clusters (Table S3 and S4, respectively); and specific lipid binding sites on the receptor (Tables S5-S7, S8). These complex properties may be the result of the size and multiple composition of the receptor and its environment (membrane, solvent, ions) and modulate its activity state.^83^

More realistic simulations could now include cholesterol, a specific and abundant membrane component, in addition to the major DPPC lipid component, and study its diffusion to the receptor surface, as well as the competitive effects between cholesterol and phospholipid.^84^ The dynamic properties of GPCRs may be modulated by the coupling to the membrane environment and its multiple structural states.^85, 86^ Finally, even though we observed distortions of the lipid bilayer for each system, their study was out of the scope of this work.

Altogether, we hope that our extensive study of the histamine H3 receptor will contribute to a better understanding of signal transduction’s molecular events.

## METHODS and MATERIALS

We use throughout this paper the Ballesteros-Weinstein generic numbering scheme for the amino acid residues of class A GPCRs.^87, 88^ The numbers attributed correspond to frame 0 of the trajectory. There is no dynamical reattribution of numbers along the trajectory.

### Homology modeling

We built the 3D structure of the transmembrane regions of the *R. norvegicus* histamine H3 receptor (Hrh3 gene, UniProt Q9QYN8, isoform 2 or rH3R(413) or short isoform H3S) with MODELLER’s sequence homology method^89^ by taking advantage of the diverse experimental X-ray structures of several homologous class A 7TM receptors that have become available in the last years, such as the β2-AR (PDB codes 2RH1, 2R4R, 2R4S, etc.). We thus chose templates that corresponded to the wished states-activated, inactivated or apo. Despite the lower-than-the-threshold (30%) overall sequence identity between H3R and other receptors needed to generate reliable 3D models, the high sequence homology per helix plus the pattern of highly conserved residues on each of the TM helices of GPCRs of class A allows us to “anchor” the H3R sequence to the template sequences and thus use homology modeling with high confidence as we are not sampling the folding of the protein. The *R. norvegicus* sequence is 93.7% identical to *H. sapiens*’; we took the short isoform of the former because its pharmacological characterization is more complete. We paid attention by choosing active state structures as templates for the HSM-H3R and H3R apo homology models, and inactive state structure templates for the CPX-H3R homology model. Thus, for the **apo** form of H3R, we used the ligand-free native opsin from bovine retinal rod cells (PDB 3CAP) whose structural features are attributed to an active GPCR state, representing the constitutive activity of the apo H1R. For the **agonist** complex, we chose the same activated state structure for docking the HSM agonist. For the **inverse agonist** complex, we used as template human β2-AR in complex with inverse agonist carazolol (PDB 2RH1). We removed the native ligands, antibodies and any other foreign molecules. The whole-sequence sequence identity between hH3R and hβ2-AR is of ∼22%, including ICL3 (∼25% excluding ICL3), below the threshold generally used for homology modeling. But, as a matter of fact, we do not consider the overall homology of the entire sequence, but rather local TM alpha-helix sequence homologies only, as the large structural differences are localized in the extramembrane ends and in the loops. Thus, for TM1, for example, the local sequence identity between hH3R and hβ2-AR is of 36%, large enough for modeling, as opposed to the 22% for the whole sequence. The TM1-TM1 values of the rH3R:hβ2-AR pair are of 47.4% identity/89.5% similar (LALIGN software, EMBnet Server).

As far as the loops connecting the TM helices is concerned, we employed a truncated form of the receptor by reducing the size of the IC3 loop (∼140 residues), given its conformational heterogeneity and the fact that its long sequence shows no homology to any experimental 3D structure. A Blast search of the 140-amino acid length of ICL3 against the PDB gives no significant results that would help for its modeling and its *ab initio* modeling is not possible due to its size. Even though we have analyzed the sequence of ICL3 and found several low-complexity regions, such as poly-Pro and poly-Ser, and other sequence fingerprints or motifs, we were not able to model this protein domain. ICL3 was then the only loop modified. Thus, when necessary, we replaced the fusion lysozyme from the template structures with several residues from the N-terminus of ICL3 and several residues from the C-terminus of ICL3, resulting in a short loop. Our use of a short isoform H3R is justified by the finding that the HSM auto-receptor is reported to be a short isoform of the H3R, H3R(413).^90^ ICL3 has been often replaced through protein engineering to produce stabilized versions of the receptor for structure determination. Some teams have tried to study the effects of ICL3 on the β2-AR.^91^ The lengths of all other loops in our models were those of the native sequence. Since they are small-sized, we were in no need to generate their conformations by other methods. The starting conformation for ECL2, which contains about 22 residues was unstructured, but it folded unto specific conformations as the simulations evolved. In all models obtained, the disulfide bridge between Cys 3.25 of TM3 and Cys 45.50 of ECL2 was satisfied. In addition, H8 in the C-ter was stable throughout the trajectory and remained in a cytoplasmic juxta-membrane-bound position, despite the lack of palmitoylation at Cys 8.59. All these features are an indication that our 3D homology models present a correct folding and are thermodynamically stable. For a more realistic and complete model, we added for the first time the N-ter of the H3R, not reported in experimental structures of homologous receptors. We achieved this by constructing first the corresponding peptide in extended conformation and then fusing it to the N-ter of the crystal structure with PyMOL. Preliminary internal energy minimizations led the N-ter to fold back from the solvent in a stable conformation. At the end, we selected for each system the model with the lowest value of the MODELLER objective function.

### Docking simulations

We generated three systems: two holo-receptors - an agonist-receptor complex and an antagonist-receptor complex; and an apo receptor, i.e. with no ligand bound to it. For the first complex, we used as the prototype ligand the endogenous agonist HSM in the (major) tele-tautomeric form; the quaternary amine at nitrogen Nς carries a positive charge. For the second complex, we took CPX as the prototype antagonist/inverse agonist. The structures of the agonist-receptor and antagonist-receptor complexes were obtained after steered molecular docking calculations in the putative orthosteric binding site with the AutoDock Vina program.^92^ For that purpose, we centered the grid box of about 30 Å of side length in the ligand pocket of the receptor, amid the TMs, ensuring thus that the search space was large enough for the ligand to translate and rotate; values for the exhaustiveness, num_modes and energy_range variables are of 12, 10 and 10, respectively. All single bonds in the ligands were free to rotate (two in HSM, and five in CPX). After the docking of the ligand to the receptor, the most representative complexes were further optimized by means of energy minimization. The stereochemical quality of the 3D models was verified with the ProCheck^93^ and WhatIf^94^ programs. Given the significative sequence homologies of H3R to other GPCRs used as templates as far as the transmembrane regions is concerned, we estimate our 3D models to be of high accuracy and corresponding to a crystal structure of ∼3 Å resolution.

### Molecular dynamics simulations

We used the Membrane Builder tool in the CHARMM-GUI server^95, 96^ for construction of the membrane and for immersing the receptor in a pre-equilibrated symmetric lipid bilayer with the Insertion method. For the protein-membrane-water system, a 1,2-Dipalmitoylphosphatidylcholine (DPPC)-based bilayer (PubChem CID 6138) was generated in a rectangular water box in which the ionic strength was kept at 0.15 M by KCl. We used this salt as we wanted to see the effects of the monovalent potassium on the receptor, as opposed to sodium. In addition, the relative number of K^+^ and Cl^-^ counterions allowed to obtain an electrically-neutral system. The cubic box lipid bilayer consisted of about 188 phospholipids solvated with a shell of water molecules and ions placed around the bilayer with the program Solvate (https://www.mpibpc.mpg.de/grubmueller/solvate). We also included buried waters in the internal cavity of the receptor with the program Dowser (http://danger.med.unc.edu/hermans/dowser/dowser.htm). The space surrounding the bilayer was cropped to a rectangular, periodic simulation box with periodic boundary conditions. This configuration allows the receptor, placed in the center of the layer, and its periodic image, to be separated by a significant distance, avoiding thus unwanted receptor-receptor interactions. The dimensions of the box were of about 80 nm by 80 nm by 100 nm and depended on the system at hand. In addition to the receptor, the explicit water solvent, the phospholipids and the K^+^ and Cl^-^ counterions, the box contained, if appropriate, the ligand. The resulting systems contained 70 000-100 000 atoms. Among the several output files generated by the Charmm-gui.org server, we used the topology files for the subsequent MD simulations. The dielectric constant of the interior of the receptor is high and like that of the aqueous solvent (∼78) due to the presence of internal waters.

The Propka^97^ program was utilized to assign the protonation states of the titratable groups of H3R at the value pH of 7.4. We payed attention to the fact that the side chains exposed to the midst of the membrane are embedded in a low-dielectric, hydrophobic environment. The ionization states of ionizable side chains in the interior of the receptor (since it is saturated with water), in the phosphatidylcholine head-group zone and in the extra- and intra-cellular zones were assigned as those corresponding to exposure to a high-dielectric surrounding. DPPC is a phospholipid that is well characterized physicochemically and whose experimental average surface area per headgroup in the gel phase is known^98^ (63±1 A^2^ at 323K). We performed MD experiments on a test membrane and noticed that it was not necessary to apply an external surface tension in the calculations as we reproduced the experimental DPPC area per phospholipid in its absence. In addition, certain GPCRs have been shown to function in cholesterol-free membranes.^77^

We employed the Nanoscale Molecular Dynamics software package (NAMD), version 2.7b1, using the CHARMM-22/CMAP force field for proteins for the simulations. HSM and CPX were parametrized with the CHARMM General Force Field (CGenFF) program of the ParamChem initiative (https://www.paramchem.org) or with the CHARMM “patch” (https://www.charmm.org/charmm/documentation/by-version/c40b1/params/doc/struct/#Patch) command that allowed us to obtain the topology and parameter files.

We used the following parameters for the molecular simulations: Leapfrog Verlet algorithm for Newton’s equation integration; integration step of 2 fs allowing the SHAKE algorithm for keeping fixed all bonds involving hydrogen atoms; update of the lists of non-covalent pairs of atoms every 20 fs; long-range electrostatic interactions with Particle Mesh Ewald algorithm, spacing 1 Å, 4 fs update; non-covalent cut-off radius of 10 Å and the non-bonded pair-list distance of 15 Å; microcanonical/NPT ensemble during equilibration and production runs; TIP3P model of water; temperature and pressure-coupled Langevin baths to ensure an isotherm and isobaric ensemble, with T=323.15 K and P = 1.013 bar. The water thickness on either side of the membrane could be up to 35 Å, depending on the system.

The simulation of the membrane-water system was performed in three stages. During the first stage, restraints on all non-solvent heavy atoms were applied; the system was energy-minimized for several hundred thousand steps using steepest-descent first and then conjugated gradient. Afterwards, several hundred thousand steps of conjugate gradient were used to minimize the side chains of the receptor and the aliphatic chains of DPPC, keeping the heads of the phospholipids and the backbone of the receptor fixed. Lastly, other several hundred thousand steps were used to minimize the free system with harmonic constraints on protein atoms, then by freeing all atoms. This procedure was followed by a slow warming-up of the system for 50 ps. In the second stage, full equilibration of the system was achieved by using the NPT ensemble for 100 ns, smoothly removing the applied constraints. The third stage of the simulation consisted of 800-900 ns production run under the NPT ensemble. We performed the equilibration of the system in the absence of external pressure on the system, allowing energy dissipation without restriction.

We took the starting frame for all our analyses to be the frame at which the RMSD reaches a first plateau. For the antagonist complex, this corresponds to nanosecond 150, and for the agonist and apo receptors to nanosecond 112. Given the behavior of HSM, only the first segment of the agonist complex trajectory, when the ligand is well bound to the receptor, is considered for the analyzes (112-590 ns).

We ran most of the calculations in the massively parallel IBM Blue Gene of the HPC center IDRIS (http://www.idris.fr) in France.

## Acknowledgments and funding sources

This study was supported by grants from:

- ECOS Nord-ANUIES Mexique and UMR-S 1204 INSERM/Université d’Evry-Val-d’Essonne/Université Paris-Saclay for visiting scholarships to LDHZ.
- Institut Servier for a postdoctoral fellowship to LMMV.
- Université d’Evry-Val d’Essonne/Université Paris Saclay for a visiting professorship to JCB.

## Author contributions

Conceptualization of the study, RCM; Methodology, RCM & LDHZ; Software, LDHZ; Formal analysis, RCM & LDHZ; Investigation RCM, LMMV, JCB & DP; Resources, JMA, PAC; Writing - original draft, writing - review and editing, RCM, JMA, DP; Visualization, LDHZ; Supervision, RCM; Project administration, RCM; Funding acquisition, RCM, JMA, PAC.

