## Supplemental data for "A complex view of GPCR signal transduction: Molecular dynamics of the histamine H3 membrane receptor"

TITLE

Short title: Molecular dynamics of H3R signal transduction

AUTHORS

L.D. Herrera-Zúñiga,[1,3,4] L.M. Moreno-Vargas,[3] L. Ballaud,[3] J. Correa-Basurto,[1] D. Prada-Gracia,[2] D. Pastré,[1] P. A. Curmi,[1] J.M. Arrang,[3] R.C. Maroun[1,3*]

[1] Structure et activité de biomolecules normales et pathologiques, UMR-S U1204, INSERM/Université d'Evry-Val d'Essonne/Université Paris-Saclay, Evry, FRANCE

[2] Computational Biology and Drug Design Research Unit. Federico Gómez Children's Hospital of Mexico City, MEXICO

[3] Laboratoire de Neurobiologie et Pharmacologie Moléculaire, Centre de Psychiatrie et Neurosciences, INSERM U894, Paris, FRANCE

[4] Área de Biofísicoquímica, Departamento de Química, Universidad Autónoma Metropolitana, Unidad Iztapalapa, Mexico City, MEXICO

Current addresses:

LDHZ: Tecnológico de Estudios Superiores de Chicoloapan, Loma de Guadalupe., 56380 Ejido de Chicoloapan, Mexico State, MEXICO

LMMV: Computational Biology and Drug Design Research Unit. Federico Gómez Children's Hospital of Mexico City, MEXICO

JCB: Laboratory for the Design and Development of New Drugs and Biotechnological Innovation, SEPI-ESM, Mexico City, MEXICO

General physical properties of the system and their definitions

- The **root mean-square deviation** (RMSD) of the cartesian coordinates of the Cα atoms measures the structural drift of a molecule. In the absence of determining free energy landscapes for the membrane-protein system, the time dependence of the RMSD is a good indicator of the convergence of the 3D structure of the protein along the simulation to a stable state.

- The **RMSD matrices** contain the RMSDs using Cα atoms only between all possible pairs of structures from the trajectory. For generating the matrices, we used every 500th structure for the calculation. The RMSD color gradient goes from dark blue for small values, through yellow for medium values, to dark red for large values, with the origin at the upper left-hand corner. The different blue squares around the diagonal show time periods within which the structures resemble each other more than to frames outside these intervals. The blue squares may be taken to represent subfamilies or ensembles of conformations. The passage from one square to the other indicates different conformer families. The local conformational transitions involved in these changes in RMSD are reflected in the high number of blue regions around the diagonal of the corresponding RMSD matrix.

- The **2D distance or contact maps** for the average structure represent the Cα atom distance between all possible amino acid residue pairs in a 3D protein structure. They can be used to describe similarity between protein structures and to represent characteristic patterns of secondary structure. The origin, i.e., residue 1 (N-ter) for the x- and y-axes is at the upper-left corner. The last residue (C-ter) is at the lower-right corner. Red zones denote proximities between atoms, and thus between helices.

- The **RMSF** as a function of the residue number is a measure of the thermal mobility or structural heterogeneity.

- **Principal Component Analysis** (PCA) represents a classic dimension reduction approach by constructing orthogonal linear combinations of many properties (in this case conformations), called principal components (PC). The greatest variance of the data lies on the first component, the second greatest variance on the second component, and so on. The PCA allows the identification of conformational communities or clusters, dividing the conformations into several populations and covering in this way a well-defined region in diversity space given by the principal components. These essential degrees of freedom describe major collective modes of fluctuation that are relevant for the function of the protein.[1] PCs may be referred here to as 'meta- or super-conformations' and are very useful in investigating the molecular motions of proteins.[2,3] Through the PCA, molecular dynamics simulations (MD) allow the sampling and the grouping of similar conformational states.

- **Mechanical properties of the helices**

Characterization of the structural changes of the α-helices, such as tilt, rotation, bending, wobble, winding-unwinding, compression-extension and displacement is important for clarifying the contribution of helices to the mechanical properties of proteins, contribution that can be important for understanding the mechanics of signal transduction in the case of 7TMs.

  - **Global tilt** defines the orientation of each helix in a protein. It is the angle that the helix axis forms with the laboratory frame's coordinate axes. The evolution of this variable is shown in dial plots in which the concentric circles represent the passage of time.

- The **global helix rotation**[4] is a measure of the rotation of the helix as a rigid body around its own axis.
- The **local tilt**, which defines the orientation of the helical axis with respect to the reference helix configuration, shows the extent of change in helix direction.
- The **bend** (kink) angle measures the angle between the axis of the helix before and after a proline residue. The closer to 0° the kink angle is, the smaller the kink. The **wobble** angle is close to zero when the post-proline helix is bent, so that its axis is moved towards the proline Cα, and close to positive or negative values when this axis is moved away from the Cα. The **face-shift** angle is a measure of the winding or unwinding of the helix. When the face-shift angle adopts positive values, it means that the helices are underwound. The converse is true, i.e., when the angle is negative, the helices are overwound.[5]
- During MD simulations, the membrane-spanning helices can undergo **compression-extension** movements beyond just random fluctuations. The measurement of the length of the helix serves to estimate this variable.

- The total displacement of the **center of mass** (COM) of the helix is used to characterize its position. The total displacement gives the contribution from the X- and Y-components, i.e. along the membrane plane. The displacement of each helix in the Z-direction, perpendicular to the plane of the membrane, can also be calculated.[4]

For the analysis of the data generated by the MD production trajectories, we used the following programs:

- EUCB[6] for the computation of the following properties, with 10 ns jumps and a switch distance of 5 Å.
    - Root mean-square fluctuations (RMSF) of each amino acid residue
    - H-bonds
    - Ionic pair matrices or charged-residue clusters
    - Conformational switches
    - Side chain rotamer angles
    - Ionic bridges
- SIMULAID[4,7]
    - Mechanical properties of the helices (TRAJHELIX)
    - Analysis of proline residues with kinks (PROKINK)
    - H-bonds between protein and solvent
- CARMA[8]
    - Root mean-square deviation (RMSD) of the cartesian coordinates of Cα atoms with respect to the initial frame
    - RMSD matrices between structures
    - Distance maps

- The principal component analysis (PCA) of the Cα atoms of the protein
- PCA-based free energy landscapes or ΔG plots obtained from the equation $\Delta pG = -kB\ T\ ln(P/Pmax)$, where P and Pmax are probabilities obtained from the distribution of the principal components for each structure (frame) of the trajectory.

- ProKink[5]
  - Bend angles
  - Wobble angles
  - Winding-unwinding angles
- VMD[9] and Tcl/TK home-brewed scripts run in VMD
  - Distances between amino acid residues
  - Detection of the residues of the internal cavity of the receptor
  - Non-covalent contacts between cavity residues and ligand
  - Distribution and residence times of lipids around the protein
  - Lipid binding sites on the protein
  - Positions and residence times of water molecules in the interior of the protein
  - H-bonds between residues
  - Visualization of the trajectories
  - Production of molecular figures

- PyMOL (PyMOL Molecular Graphics System, Version 2.0 Schrödinger, LLC.)
    - Molecular graphics and visualization

The residues composing the internal cavity of the representative structures of the receptor produced by CARMA were defined by finding the "wet" residues, i.e. those residues that were in contact (3 Å or less) with an internal water for more than 80% of the trajectory. The "wet" atoms of the ligands were determined in the same fashion. This approach determined the residues that had an interaction with a water molecule that had itself an interaction with the ligand. On another hand, pocket detection, ligand binding site and analysis and visualization of tunnels and channels were performed with the Fpocket,[10] SiteHound[11] and Caver software tools.[12] Those residues found in at least three out of four searches were considered cavity residues.

We did not deem necessary to energy minimize the complexes at the frames we looked at during the analysis. Instead, we selected ten frames at regular intervals of the MD trajectory to obtain the interactions between the ligand and the receptor, since we compiled the long-lasting neighbors of the former all along the trajectory.

When dealing with the lipid components of the membrane, we wished to find out whether there were any DPPC-specific binding sites on the receptor. For this purpose, we found out the amino acids that are in contact at least 80% of the time with at least one atom of a lipid molecule defining high-affinity, specific binding sites with the membrane protein.

We performed the computations in a Linux cluster with one master node of 8 CPUs, 10 To stocking; and 192 CPUs (16 nodes of 12 Intel XEON E5630@2.53GHz, 24 GB RAM, CPUs each). The NAMD parallel jobs were executed through the mpiexec application.

RESULTS

Selected results

In this section, we describe the outcome of a multidimensional analysis of many microscopic properties of each of the three systems dealt with in this work. We mention below some selected results.

- RMSD of Cα atoms and RMSD matrices

o The MD trajectories lead to physically stable systems (Fig. S1).

o For each of the three systems, subsets of similar conformations appear during the trajectory, indicating structural variations (Fig. S2).

- Cα-Cα distance maps

o The overall relative positions of TM helices in the apo receptor and the agonist-H3R complex resemble each other (Fig. S3).

- RMSF

o The overall RMSF values are in the order antagonist complex > agonist complex >> apo receptor (Fig. S4).

- Principal Component Analysis (PCA)

o Eigenvectors

Presence of long-range communication between the extra- and intra-cellular regions of the two receptor complexes through their membrane region (Fig. 3, 4 of main text). Correlation movements between the N-ter and H8 for the apo receptor (Fig. 5 of main text).

Coordinates along the paths connecting the clusters represent transition state conformations.

- Characterization of the internal cavity of the receptor

o Charged residue clusters; H-bonded clusters

There are two charged amino acid clusters in the two receptor complexes (Fig. S7, S8), as compared to four clusters in the apo receptor (Fig. S9).

The antagonist-bound receptor shows the largest number of water-mediated H-bonds (Fig. S10).

In all three systems, water occupancies in the binding cavity are characterized by very low residence times and exchange only with the bulk solvent in the extra-cellular region. The antagonist complex lodges the largest number of water molecules in the binding cavity (Table S2, upper-left box).

- Hydrophobic clusters

o In addition to the existing H-bonds, differential non-covalent interactions like hydrophobic clusters of residues contribute to stabilize the overall structure of the receptor in each one of its states (Table S3).

- Mechanical properties of the helices

o Global X-, Y- and Z-tilt

The average values of the global X-, Y- and Z-tilt angles for each TM helix of each of the three systems show rather small dispersions around each average value. As the trajectory evolves, the corresponding values for H8 have larger variations (Fig. S19abc-S26abc for the antagonist complex; S27abc to S34abc for the agonist complex; S35abc to S42abc for the apo receptor).

o Global helix rotations

The antagonist receptor and the apo form have in common a negative value for the average rotation of the eccentric TM4 helix. The rotations of the helices seem to be correlated (Fig. S19d-S26d for the antagonist complex; S27d-S34d for the agonist complex; S35d-S42d, for the apo receptor).

- o Local helix tilt

For all three systems, all TM helices show average values of 0-10° in helix direction, with little dispersion about these values (Fig. S19e to S26e for the antagonist complex; S27e to S34e for the agonist complex; S35e to S42e for the apo receptor); helix direction and fluctuations in direction are larger for H8.

- o Turn angle per residue (TPR)

Large variation of H8 of the ago receptor (Fig. S34f).

- o Bend (Kink), Wobble and Face-shift angles

Section 1 of the **agonist-H3R** complex shows rather large variations in the bend angle values of TM5-TM7, especially TM7 (Fig. S44g), as a function of time. As for the wobble angle, TM5 shows variations around ±125°, with values around ±π for TM6 and TM7 (Fig. S44beh). The face-shift values for the three helices are of ±π (Fig. S44cfi); whereas those for the **antagonist-H3R** complex are of 25°-75° (Fig. S43cfi); for the **apo receptor** these values are of 25°-85°(Fig. S45cfi). Thus, helices TM5 to TM7 of the agonist-H3R complex tend to unwind, whereas the same helices in the antagonist-H3R complex and the apo receptor are underwound.

- o Helix length (compression-extension)

In all three systems, TM3 (36 residues), TM5 (24 residues) and TM6 (33-34 residues) are the helices whose average length is the largest and protrude into the aqueous phase of the intracellular side of the membrane.

o Total helix displacement

The eccentric TM4 helix may play a role in the inactivation mechanism of H3R by the antagonist through its total displacement (6 Å) on the plane of the membrane and its shift in the Z direction (7 Å) (Fig. S46a TM4).

o Met side-chain switches

The conformational states of the methionine residues remain unchanged between the three states - there are thus no conformational changes of the methionines upon activation in H3R.

- Lipid binding sites

o The antagonist-H3R complex shows the most lipid binding sites and involves upper- and lower-leaflet lipids (Fig. S15).

#### General results

- RMSD plateaus of the MD trajectories imply physically stable systems

Fig. S1 shows the RMSDs of the Cα with respect to the initial structure. For the **antagonist-H3R** complex, the RMSD attains a plateau at about 150 ns, remains constant at 4.2 Å until ~550 ns and then rises slightly to 4.8 Å. The **agonist-H3R** complex shows an RMSD that attains values ~5.0 Å at 600 ns until the end of section 1. This observation is consistent with the fact that agonist ligands tend to be less efficient in the stabilization of the structure of the protein.[13] The **apo** receptor shows considerable changes in the RMSD, especially in the middle of the trajectory, just before 600 ns, when it reaches almost 5 Å of RMSD, to fall afterwards to around 3.5 Å at the end of the 930 ns trajectory. All in all, the observed fluctuations suggest a good packing quality of each model.

For the structurally-conserved TM helices of the receptor, the RMSDs of each of the three systems with respect to the crystal structure of the H1R-doxepin complex at 3.1 Å resolution[14] (PDB code 3RZE) are as follows: 3.1 Å for the last frame of the antagonist-H3R complex, 2.5 Å for the last frame of the agonist-H3R complex, and 2.0 Å for the apo receptor. These data indicate that the MD trajectories lead to physically stable systems.

- RMSD matrices point to high structural variations

Fig. S2a for the **antagonist-H3R** complex shows a tendency for the blue areas to get darker as the trajectory unfolds, indicating that subsets of similar conformations persist. For the first period (section 1) of the **agonist-H3R** complex (Fig. S2b), i.e. when histamine is bound to the receptor (up until nanosecond 590), the structures up to 328 ns resemble each other a lot (small RMSD values) at the beginning of the first phase of the period. Then about five subsets of conformations appear, of which the last one is maintained sometime after histamine unbinding. Then another subset appears centered around 726 ns. For the **apo** receptor (Fig. S2c), the RMSD matrix is rather sparse and weak in blue regions, i.e. in small values of RMSD, indicating higher structural variations.

- Cα-Cα distance maps indicate the overall relative positions of TM helices in the apo receptor and the agonist-H3R complex are equivalent

The Cα-Cα distance map of the average structure of the **antagonist-H3R** complex shows that CPX (represented in the map after all the amino acid residues) is in contact with TM2, the N-ter of TM3, ECL2, the C-ter of TM6 and the N-ter of TM7 (Fig. S3a). For the first section of the trajectory of the **agonist-H3R** complex, the proximity relationships are in Fig. S3b and for the **apo** receptor in Fig. S3c. All relative positions of the helices are shown in the form of a matrix for each of the three systems in Fig. S3d, in which an x represents the mentioned proximities between TM helices. The matrices are symmetrical. We can see that the relative TM helix positions of the average structure of the agonist-H3R complex and apo receptor are similar, with a clear off-diagonal line showing a sequential array of proximities: TM1-TM2-TM3-TM4-TM5-TM6-TM7. The inter-helix distribution of spatial distances for the antagonist-H3R complex shows no proximity between TM2 and TM4, TM3 and TM7, and TM4 and TM5; instead, this complex shows a proximity between TM2 and TM6. As mentioned in the Introduction, H3R possesses a high constitutive activity in the absence of agonist. Our results show indeed that the overall relative positions of TM helices in the apo receptor and the agonist-H3R complex resemble each other. In addition, ECL2 is in contact with the antagonist, in conformity with the importance of this loop in ligand binding, selectivity and (in)activation,[15] as well as stabilization of the inactive state of the receptor.[16]

- RMSF values highlight structural heterogeneity

Fig. S4 shows the RMSF values as a function of residue number. For the **antagonist-H3R** complex, the graphic shows that the highest values come from TM3 (16 Å) and TM5 (17 Å).

The other trans-membrane helices show a decrease in RMSF when going from the N- to the C-ter. Instead, TM3 and TM5 show large increases towards the C-ter. This amounts to larger structural fluctuations for the fragments of the TM2- TM5 in the lower leaflet of the membrane, and for the segments of TM1 and TM7 in the upper leaflet of the membrane. As far as the first section of the **agonist-H3R** complex is concerned, the fluctuations are more homogeneous between the ends of the helices. Astonishingly, the values of the fluctuations of the **apo** receptor are much lower, in the 7 to 9 Å range. Like the antagonist-H3R complex, the structural heterogeneity is predominantly localized to the lower leaflet of the lipid bilayer, although of about half the magnitude.

In conclusion, the overall values of the RMSFs are in the order antagonist complex > agonist complex >> apo receptor.

With regards to the ligands, the RMSF average values of HSM and CPX are 13.5 and 40.2 Å, respectively, indicating the increased flexibility of CPX with respect to HSM.

- Free energy landscapes are different for each receptor state

For the **antagonist-RH3 complex**, the PC1-PC3 map (Fig. S5a) has a laid S-shape and shows almost no energy barriers among the clusters in Q1. There are also about four clusters. The PC2-PC3 map (Fig. S6a) shows a free energy landscape in which the four clusters are centered around the origin and fusion into essentially two wide clusters positioned at Q1-Q2 and Q3-Q4. Fig. S5b shows the corresponding PC analysis for PC1 and PC3 of the **agonist-H3R complex**, with a double energy well at Q1-Q2. The corresponding PC2-PC3 map (Fig. S6b) shows several clusters in a C-shape at Q1-Q2. The PC1-PC3 map for the **apo** receptor shows a deep cluster in the origin of the plot (Fig. S5c). The energy wells in the PC2-PC3 map (Fig. S6c) are rather disperse, with a large "cluster of clusters" at Q3-Q4. Receptor conformations in-between the clusters correspond to transition states.

- Leu is the most frequently membrane-exposed residue

The fraction of membrane-exposed hydrophobic amino acid residues is of 71, 71 and 67% for the antagonist, agonist (section 1) and apo systems, respectively (red boxes in first column of Table S9), with Leu being the most frequently exposed residue, representing 27-29% of all membrane-exposed residues. With respect to all exposed residues, the exposed hydrophilic residues, represent 22%, 12%, and 18% for the antagonist complex, for the agonist complex and for the apo receptor, respectively. As far as the charged residues is concerned, their relative exposed populations are of 8%, 17% and 15%, for the antagonist complex, for the agonist complex and for the apo receptor, respectively.

- Clusters of charged residues get reconfigured from one state of the receptor to another

Electrostatic interactions are of special importance for membrane proteins because of the low dielectric environment in membranes. We thus proceeded to detect clusters of charged residues using a distance criterion of 12 Å between COMs of the side chains of amino acid residues.

For the **antagonist-H3R** complex, there are two charged amino acid clusters that persist 90% of the time of the production trajectory. One cluster is in the extracellular region of the receptor and the other one in the intracellular region. The two clusters are represented in Fig. S7. The amino acid residues for the first cluster belong to TM3, ECL2, TM5-TM7, and those of the second cluster to TM1, ICL1, TM3- TM6, ECL1, ICL2, ICL3 and H8. For the **agonist-H3R** complex, section 1, two independent clusters persist while HSM is bound to the receptor (Fig. S8). They involve N-ter, TM3, ECL2, TM6 and TM7 for the first network; and TM3- TM6 for the second. One residue of the first cluster, Asp 3.32, is in contact with the protonated amine (pKa ~9.4) of the aliphatic amino group of HSM, consistent with experimental data.[17] Four clusters of charged residues can be found for the **apo** receptor (Fig. S9). Amino acid residues from TM1, TM3 and ECL2 compose the first cluster; one residue from each TM3, 5 and 6 the second cluster; residues from the N-ter, ECL1 and TM6 the third cluster; and residues from TM1, ICL1, ICL2, TM3-TM5, ICL3, TM7 and H8 the fourth cluster. Many of these residues are also involved in long-lasting H-bonds, i.e. those prevailing more than 70% of the time.

- H-bonded networks between amino acid residues are a function of the state of the receptor

For the **antagonist-H3R** complex, inter-residue H-bonds with residence times greater than 70% connect ECL2 (TM4-TM5 loop) and ICL2 (TM3-TM4 loop) to TM3; N-ter to TM7; TM1 to H8; and ECL2 to ECL2 (Table S1). For the first segment of the production trajectory of the **agonist-H3R** complex, ECL2 is H-bonded to the N-ter, TM3 and TM4; and the N-ter of TM1 to H8 (Table S1). Finally, for the **apo** receptor, only one non-intra-helical H-bond persists (Table S1). Thus, it is interesting to notice that the antagonist-bound receptor shows the largest number of inter-residue H-bonds, providing the structure of this complex with an added energetic and structural stability.

- The composition of the internal cavity of the receptor abounds in Tyr and Leu

Table S2 shows that there are 38 residues that form the orthosteric cavity of the CPX-bound receptor. These residues are contributed by TM2, TM3, ECL2, and TM5-TM7. The environment of the ligand is rather hydrophobic with the three aromatic amino acid side chains being represented. Twenty-six residues compose the internal cavity of the agonist-H3R complex, with contributions coming this time from TM1, TM3, ECL2, and TM5-TM7. For the apo receptor, the cavity is composed of only 24 residues. As expected, this cavity is the smallest one, given the absence of ligand and the subsequent contraction of the orthosteric cavity. Secondary structures contributing to it are TM2, TM3, ECL2, and TM5-TM7. In all three systems, TM4 being eccentric, it contributes with no residues to the receptor's cavity, just like all the loops (except ECL2) and, of course, H8. Notice the contribution of ECL2 to the morphology of the binding cavity.

The residues that compose the internal cavity of H3R in its different states are of diverse types, with Tyr and Leu being the most abundant, followed by Phe and Ser. Residues Pro, Gly, Gln, Lys, Thr and His are absent (Table S2). Interestingly, the antagonist and agonist pockets show a net negative charge of 4, whereas the net charge of the pocket of the apo receptor is neutral (Asp 3.32 + Arg 6.58). The cationic ligands HSM and CPX thus induce changes in the pocket that bring into play acidic residues.

- Water molecules in the internal cavity and their H-bonded networks are persistent

A persistent dynamic network of water molecules is observed in the interior of the receptor, with waters penetrating and exiting the receptor from the extracellular region. These waters establish several H-bonded networks; nevertheless, the receptor is not a water channel since it presents a molecular plug formed by the lower-leaflet internal walls of TM2, TM3, TM6 and TM7, and the TM7-H8 loop that keeps the internal water molecules from exiting to the cytoplasm or cytoplasmic bulk waters from entering the receptor. About 61 water molecules are constantly present in the internal cavity of the **antagonist**-bound receptor during the trajectory, 35 for the **agonist** one, and 49 for the **apo** receptor(upper left of Table S2), so that water penetration to the receptor is largest upon antagonist binding, and smallest for the agonist-H3R complex, just like in the high-resolution structure of α2A-AR that reveals about 60 internal waters.[18] In addition, Yuan et al.[19] mention an increased penetration of water into the receptor cavity of μ- and κ-opioid receptors, which has been linked to the activation mechanism upon agonist binding. Therefore, the presence of water in the binding site clearly demonstrates its influence in the dynamics and conformation of the receptor.

We determined the H-bonded networks established by water molecules in the cavity with occupancy greater than 75%. In all three systems, these occupancies are characterized by very low residence times, of the order of 20% maximum, indicating a high fluidity of water molecules. Thus, even if a site may be continually hydrated, it is not so by the same water molecule 4/5 of the time. Again, the observation of the itinerary of the water molecules indicates that these enter and exit the internal cavity of the receptor through the extra-cytoplasmic region of the receptor only, no water molecules neither entering nor exiting the receptor through the cytoplasmic region. The internal cavity of the receptor has the shape of a bent funnel with a flexible lid at the top composed of the N-ter and the ECLs, especially ECL2.

For the **antagonist-H3R** complex, residues that contribute with their side chains to H-bonded networks in the cavity zone are Asn 7.45; Asp 2.50 and Asp 3.32; Glu 5.46 and Glu 7.36; Ser 7.46; Thr 6.52; Tyr 2.61; and Trp 7.43 and Trp 23.50 (Fig. S10). Of these, Asp 2.50, Glu 5.46 and

Ser 7.46 form two or more H-bonded bridges. The two largest networks involve five waters, Asp 2.50 and Ser 7.46 (Network 5); and four waters, Glu 5.46 and Thr 6.52 (Network 6, Fig. S10). Finally, the interaction of CPX with two residues in the protein, Tyr 2.61 and Asp 3.32, is mediated by two water molecules for the former, and by one for the latter (Netwk3, Fig. S10). Notice that TM1 and, of course TM4 and H8, do not participate to the H-bond network during CPX binding. The only charged amino acids in the interior of the receptor are in the binding cavity and are Asp 2.50 and 3.32, and Glu 5.46 (Fig. S10). The **agonist-H3R** complex shows two networks of H-bonds, the first of which includes HSM bonded to a water molecule and to Asp 3.32 (Fig. S10). The two networks of the **apo** receptor involve, each one, two internal water molecules (Fig. S10).

- Hydrophobic clusters may contain up to seven residues

We report in Table S3 the hydrophobic clusters at 90% occupancy formed by at least three side chains for the antagonist-H3R complex, the agonist-H3R complex and the apo receptor, respectively. The cutoff used is of 6 Å between the COMs of each amino acid residue; the set of residues is composed of Ile, Leu, Val, Phe, Met, Cys, Trp, Pro and Ala.

For the **antagonist-H3R** complex, there are six clusters. The distribution of clusters is as follows: two clusters of three residues, two clusters of four residues, one cluster of six residues and one cluster of seven residues. One cluster, implying the N-ter and ECL1 contains 7 residues (Ala 23 from the N-ter, Phe 2.56, Cys 2.57, Leu 2.60, Trp 23.50 (ECL1), Leu 3.24 and Trp 3.28). For the **agonist-H3R** complex, there are six clusters, and for the **apo** receptor there are five clusters. These non-covalent interactions, just like the H-bond networks, depend on the state of the receptor.

As for the clusters containing aromatic side chains only, the **antagonist-H3R** complex shows a 3-residue cluster; the **agonist-H3R** complex trajectory shows none; and the **apo** receptor show two clusters of 3 residues each (Table S4).

Moreover, we decided to detect the interactions between the aromatic side chains belonging to hydrophobic clusters to analyze their evolution during the MD simulation. For this purpose, we proceeded to use the COMs of the multi-atomic functional groups. The pairs of side chains studied were obtained from the hydrophobic clusters just described. For the **antagonist-H3R** complex, Fig. S11a shows an aromatic cluster composed of Phe 2.56 and Tyr 2.64 of the C-terminus of TM2, Trp 23.50 and Phe 23.52 (ECL1), and Trp 3.28. This cluster remains stable throughout the trajectory and locks the conformation between the C-terminus of TM2 and ECL1. The Tyr aromatic cycle is perpendicular to the Trp cycle, which in turn is perpendicular to the Phe cycle. This type of non-covalent interaction involving aromatic rings ($\pi$-$\pi$ or $\pi$-stacking) has been previously documented.[20] The second cluster is composed of Tyr 7.53 of the NPXXY microdomain (C-ter of TM7), which establishes a long-lasting $\pi$ - $\pi$ interaction with the first residue of H8, Phe 8.50 (Fig. S11b). The distances between the COMs of the corresponding pairs are in the 4.5-8.5 Å range. For the **histamine** complex, we found no such aromatic ring clusters. Lastly, the **apo** receptor presents two clusters. The first one is composed of Phe 6.44, Phe 7.39 and Trp 7.40; the second cluster is like the first cluster in the active complex (not shown).

- Side-chain switches for the Met residues are not detected

Kofuku et al.[21], by monitoring the NMR signals, investigated the role of Met residue 82 (2.53) in antagonist- and partial agonist-bound states of the β2-AR, which are correlated with conformational changes of the transmembrane regions upon activation. The corresponding residue in H3R is a valine and none of its neighbors is a methionine; nevertheless, we decided to monitor the conformational states of all Met residues for the three systems. We found that in all three systems, all but one of the methionine residues remain in the t or g- states throughout the trajectory. As opposed to the β2-AR then, the conformational states of the methionine residues remain unchanged between the apo receptor, and its antagonist- and agonist-bound states; there is thus no correlation of conformational changes of the methionines upon activation in H3R.

- Detection of lipid binding sites on the receptor leads to amino acid sequence motifs

We list in Table S5 the DPPC lipids attached to the receptor in the **antagonist** state and the amino acid residues they interact with. The residence time of a given lipid is listed in the first line of the “occupancy” column; the next lines show the residence times of the lipid with different amino acid residues of the receptor. We can see that one DPPC molecule (125) is permanently bound to the N-ter residues of TM6 Lys 6.32 and 6.35, Ser 6.36, Ile 6.39 and 6.43, and to the C-ter residues of TM7 (Leu 7.52 of the NPXXY motif and Leu 7.55; Cys 7.56 and His 78.00 belonging to loop TM7-H8). This lipid is part of the lower sheet of the membrane, on the luminal side. Another lipid binding site (lipid 110) is composed mostly by residues from TM3 (Val 3.40 and 3.44, Phe 3.41, Leu 3.45 and 3.52, and Tyr 3.48) and is an upper leaflet lipid. Lipid 161 lies proximal to residues from the N-ter of TM4, making it a lower layer lipid. An additional lipid in the upper leaflet (79) binds to two residues of the N-ter end of the receptor and to the N-ter of TM7. The first half of TM5 is occupied by an upper-leaflet lipid (31 or 33). A minor binding site is formed by the ICL3 loop and the N-ter of TM6 (lower-leaflet lipid 165).

Lipid 43's site is made up of N-ter TM3 residues and C-ter TM4 residues. Furthermore, a site composed by about 12 residues from the C-ter of TM7, the N-ter of H8 and the central region of TM1 binds a lower-leaflet lipid (149). An interhelical N-ter TM3 – C-ter TM4 site lodges an upper leaflet lipid. As can be seen from Table S6 for the **histamine** complex, lipid 2 is lodged in the middle section of TM6 and establishes one contact with one residue of H8; lipid 47 binds to the C-ter of TM5 and all along TM6; lipid 183 binds to the second half of TM2; lipid 7 interacts with H8; yet lipid 22 is in contact with ICL1, the last residue of TM1, and the first residue of TM2; lipid 59 is in contact with C-ter residues of TM5. In the case of the **apo** receptor, Table S7, the first lipid binds to the N-ter of TM1 and TM7, and to the C-ter of TM2; the 2$^{nd}$ lipid to the N-ter of TM3, to the C-ter of TM4; the 3$^{d}$ lipid to the N-ter of TM1; the phosphate head of the 4$^{th}$ lipid to an Arg of the N-ter; the 5$^{th}$ lipid to the C-ter of TM6.

In résumé, for the **antagonist-H3R** complex, the structures in contact with the 13 lipids are TM1, ECL2, TM3-TM6, ECL3, TM7, TM7-H8 loop, and H8 -TM2 not in contact (Fig. S15, Table S5). For the **agonist-H3R** complex, the following secondary structures participate to the binding of the seven lipids: ICL1, TM2, TM5, TM6 and the TM7-H8 loop (Fig. S15, Table S6). Interestingly, for the **apo** receptor the 5-lipid binding site involves all helices except TM5 (Fig. S15 and Table S7).

Table S8 shows the amino acid residues of the receptor grouped in four classes -non-polar aliphatic, aromatic, uncharged polar and charged- and their population in contact with lipids for each of the three systems. First, the fraction of non-polar residues (43-56%) is the largest, followed by the aromatic residues (15-26%), and the polar and charged classes. The most frequent residue of all groups and for all three states of the receptor is leucine. In the second class, Phe is the most abundant for the antagonist and agonist (section 1) systems, and Tyr and Trp for the apo system. In the polar group, Ser and Thr for the antagonist- and agonist-H3R complexes are the most represented residues. Finally, the antagonist-H3R complex is the system showing the most contacts of charged residues with lipids -eleven Arg and six Lys. The non-polar and aromatic class residues are in contact with the larger area acyl-tails of the lipids; whereas the basic residues interact with the negatively charged lipid head groups.

Table S5 for the **antagonist** complex shows 13 highest-occupancy lipid molecules associated to the receptor, of which four are in the upper-leaflet of the membrane and nine in the lower-leaflet (Fig. S15). As an illustration of the interaction of DPPC lipids with the inactivated H3R, Fig. S12 shows a 2D LigPlot+ diagram of lipids 125 and 149, showing H-bonds between Arg 8.51 and one oxygen atom from the phosphate head, and hydrophobic interactions with surrounding residues for lipid 149. Both lipids share Leu 7.55 as an interacting residue. In Fig. S13, extra- and intra-cytoplasmic views of the inactivated receptor for the 490 ns frame, we can see that DPPC molecules essentially bind to opposite sides of the exposed surfaces of the receptor. On one side of the receptor TM1, TM6 and TM7 lodge three lower-leaflet lipids and one upper-leaflet lipid. The other side, with TM3, TM4 and TM5 offers another binding site for two lower- and two upper-leaflet lipids. On that side, an upper-leaflet lipid finds a binding site provided by TM3 and TM4. Both leaflets contribute thus with lipids for receptor binding. For the **histamine** complex, all in all, there are five lipids in the lower part of the membrane, one in the upper part, and one in the middle (Fig. S15, Table S6). A LigPlot+ diagram illustrates the surroundings of lipid 2: lipids 31, 48 and 66, Lys 6.32, Lys 6.35, Ile 6.39, Gly 6.45, Leu 6.46, Val 7.48, Leu 7.55 and Cys 7.56 (Fig. S14a). The Lys residues interact with the phosphate moiety of the DPPC molecule, whereas the hydrophobic residues with its fatty acyl tail chains. For the **apo** receptor, there are five lipids binding to sites in the upper leaflet of the double bilayer (Fig. S15, Table S7). The LigPlot+ representation of Fig. S14b shows lipid 184 and its neighboring residues (Leu 1.42, Tyr 7.33, Glu 7.36, Thr 7.37, Trp 7.40, Ala 7.44 and Ala 7.47), all but one belonging to TM7. Nε1 of Trp 7.40 interacts with O22 of the DPPC lipid; Leu 1.42 and Ala 7.47 interact with one of the acyl tails of the lipid. Many water molecules surround the phosphate and choline moieties.

In order to extract the lipid-binding H3R amino acid sequence motifs, we selected those amino acid residues in Tables S5-S7 in contact with lipids that show residence times ≥ 90%. The resulting binding motifs are the following:

**Antagonist** complex

| DPPC | Motif | | Region |
|---|---|---|---|
| 33 | $N^{45.57}$ | $Y^{5.37}$ | ECL2, TM5 |
| 79 | $F^{0.00}$ | $^{7.33}YX_3TX_2W^{7.40}$ | N-ter, TM7 |
| 110 | $^{3.41}FX_3LX_2Y^{3.48}$ | $L^{5.45}$ | TM3, TM5 |
| 125 | $^{6.35}KX_3I^{6.39}$ | $^{7.55}LC^{7.56}$ | TM6, TM7 |
| 161 | $F^{3.41}$ | $^{4.41}RX_2RKX_2L^{4.48}$ | TM3, TM4 |

**Agonist** complex

| DPPC | Motif | Region |
|---|---|---|

| | | | | |
|---|---|---|---|---|
| 47 | $^{5.55}TX_2NLX_2Y^{5.63}$ | $^{6.31}KX_2AKX_2AX_2V^{6.41}$ | | TM5, TM6 |

**Apo** receptor

| DPPC | Motif | | | Region |
|---|---|---|---|---|
| 151 | $^{3.23}GX_2KLX_2VVX_2L^{3.34}$ | $^{4.53}AX_3Y^{4.57}$ | | TM3, TM4 |
| 184 | $^{1.35}LX_2LX_3L^{1.42}$ | $^{2.61}YX_3V^{2.65}$ | $^{7.33}YX_2ETX_2WX_3A^{7.44}$ | TM1, TM2, TM7 |

- Rotamer toggle switches are multiple and concerted

The goal of this section is to determine those amino acid side chains that undergo concomitant side-chain conformational changes. We focus on the aromatic side chains of Tyr, Trp and Phe, since several rotamer toggle switches dealing with those residues have been reported in the literature.[22]

In Fig. S16 is shown for the **antagonist-H3R** complex the time evolution of aromatic side chain dihedral angles $\chi_1$ (N-Cα-Cβ-Cγ) that undergo side chain conformational transitions. We can see that $\chi_1$ of Trp 6.48 of the CWXP ($^{6.47}$CysTrpXPro$^{6.50}$) motif adopts values of -60° (g-configuration) most of the time, until 665 ns, when it undergoes a transition to a trans conformation (±180°). Concomitantly, $\chi_1$ of Trp 1.31, Trp 45.36 and Phe 5.47 undergo correlated conformational transitions. Moreover, another concurrent $\chi_1$ change for Phe 3.51

and Trp 7.43 can be observed earlier, at around 615 ns (Fig. S16). Most of these residues are in the upper half of the receptor (towards the extra-cytoplasmic zone), suggesting that, at this phase of the mechanism, conformational changes in this zone may be enough to transmit the signal to the lower half of the receptor. Given that several of these side chains are rather far from each other, it seems that other side chains mediate the transmission across the membrane. As compared to the MD simulations of Nygaard et al.[23] (their Fig. 5a), Trp 6.48 (TrpVI:13) in our extended MD simulations adopts an intermediate conformation between the active and inactive states. For instance, starting at about half of the trajectory, the average distance between the center of masses (COMs) of the Phe 5.47 (Phe V:13) and Trp 6.48 side chains is ~13 Å. Tyr 7.53 of the NPXXY motif ($^{7.49}$AsnProXXTyr$^{7.53}$) is in the active conformer state throughout the trajectory (not shown), showing no switch; it engages in a hydrophobic interaction with Ile 6.40[23] (their Fig. 5b). Thus, in a remarkable fashion a multiple toggle rotamer switch of certain aromatic chains, along with bi-modal switches appears to be important for receptor H3R inactivation. For the **agonist-H3R** complex (section 1), the conformation of $\chi_1$ of Phe 5.47 remains constant at g- and Phe 3.41 undergoes a g- to trans transition in the last fourth of the trajectory (not shown). Concomitant transitions in $\chi_1$ of several aromatic side chains – Phe29 of the N-ter, Trp 3.28, Tyr 45.51 and Tyr 45.56 of ECL2, Trp 5.36, Phe 5.38 and Tyr 7.33- take place starting at about 0.5 µs (Fig. S17). Again, a multiple toggle rotamer switch of aromatic side chains takes place during H3R activation and involves several amino acid residues. Only Trp 6.48 is common to both mechanisms -activation, inactivation. All but one of the residues are in the upper zone of the receptor and imply the ECL1 and ECL2. The number of aromatic side chains that undergo conformational transitions in the **apo** receptor is much reduced and includes only Phe 45.54 and Tyr 45.56 of ECL2, and Tyr 7.33 and Phe 8.54 side chains, with the first and the third toggle switches interconverting between g+ (+60°), g- (-60°) and trans states, and the second and third toggle switches common to the activation mechanism (Fig. S18). The mentioned residues are in the upper and middle zones of the receptor, except for Phe 8.54, belonging to H8. None of the residues of the DRF ($^{3.49}$AspArgPhe$^{3.51}$) motif are thus involved in H3R activation or constitutive activity implicating that the ionic lock involving Arg3.50-(Asp 6.30 and Asn 5.64) is not conserved. This is consistent with the lack of direct evidence for a role of that motif in those two situations.[24]

In résumé, there are multiple concerted long-range rotamer switches for each system as the involved residues are several tens of angstroms away from each other. On the other hand, the number of aromatic residues undergoing simultaneous transitions upon activation is larger than in the antagonist-H3R complex and the apo receptor and involves residues from the N-ter, TM3, ECL2 (TM4-TM5 loop), TM5, TM6 and TM7. For the apo receptor, the aromatic side chain rotamer transitions involve just ECL2, TM7, plus the C-terminus fragment of helix H8. Finally, it is interesting to observe that aromatic side chains switches from ECL2 participate in the different mechanisms.

- Inter-residue contacts are unique to each state of the receptor; novel ionic locks are detected

We have measured many selected inter-residue distances using the COMs of the side chains to detect contacts. Those pairs of residues interacting according to the criteria described in the M&M section are in Table S10 for each of the three systems. Several of those interactions represent ionic locks.

For the **antagonist-H3R** complex, the conserved arginine of the ionic lock (motif DRF) among rhodopsin-like G protein-coupled receptors, Arg 3.50, interacts with Asp 3.49 and Asp 6.30 in a constant but dynamic fashion throughout the trajectory, in agreement with the behavior of the ionic lock in the MD simulations of the β2-AR receptor.[25] In addition, there is formation of a hydrophobic interaction between Met 1.39 and Trp 7.40/Trp 7.43, followed by disappearance of an initial Met 1.39-Tyr 2.61 interaction; the interaction between Met 6.55 and Tyr 3.33/Phe 7.39 is sporadic. Lastly, the distribution of the Met 6.55-Tyr 6.51 distance with a major peak at 5 Å and a minor one at 9 Å is analogous to the one described in the literature (Nygaard et al 2013, their Fig. 4c).[26] For the **agonist-H3R** complex, the following interactions emerge: Met 1.39-Trp 7.40, Met 6.55-Phe 5.48, while the Phe 5.47-Trp 6.48 distance becomes distended; Met 6.55-Tyr 6.51/Tyr 3.33, and Tyr 7.53-Phe 8.50 distances are constant throughout. For the **apo** receptor, are stable throughout the trajectory the following contacts: Asn 1.50-Pro 7.50, Asn 2.39-Asp 3.49/Arg 3.50, Asp 3.32-Trp 7.43, Asp 2.50-Pro 7.50, Met 6.55-Tyr 6.51, Met 1.39-Tyr 2.61/Trp 7.40/Trp 7.43, Met 1.54-Phe 2.51, and Trp 6.48-Ser 7.46. A Met 6.55-Tyr 3.33 interaction goes away gradually. Those distances that undergo a notable change include: formation of interactions Phe 5.47-Trp 6.48, and Tyr 7.53-Phe 8.50/Phe 8.54; sporadic interactions Arg 3.50-Asn 2.40/Asp 3.49/Asp 6.30, and the Met 4.46-Trp 4.50 couple.

In conclusion, only the apo receptor presents interactions between TM1 and TM2, and between TM2/TM3 and TM7. Interactions between TM3 and TM6 are unique to the agonist and antagonist-H3R complexes. Asp 3.32 in the apo receptor not being involved in interaction with a ligand interacts now with a Trp residue from TM7 (Trp 7.43). Notice that the Arg 3.50-Asp 3.49/Asp 6.30 H-bond and electrostatic interaction is unique to the antagonist-H3R complex. The Asn 2.39-Asp 3.49/Arg 3.50 interaction is absent in the agonist-H3R complex, just like interactions between TM7 and H8 are absent in the antagonist-H3R complex (not shown).

- Mechanical properties of the helices

Global tilt

Fig. S19abc to Fig. S26abc show the three components -X, Y and Z- of the global helix tilt angle for each of the eight helices of the **antagonist-H3R** complex. The X- and Y-components lie on the plane of the membrane whereas the Z-component is perpendicular to them. As can be seen, the X- and Y-components adopt values in the range 60-120° for all TM helices. The Z-tilt of TM4 is 10°, making of this the most perpendicular helix to the plane of the membrane. The axis of H8, being quasi parallel to the plane of the membrane, shows a global Z-tilt of 90°. Odd-numbered helices TM1, 3, 5 and 7 have their global Z-tilt angles in the range 145-160°, whereas even-numbered helices TM2, 4 and 6 have it in the range 10-30°. This is expected since helices whose N- to C-ter directionality is directed towards the cytoplasmic space have an opposite directionality to those ones pointing to the extracellular space. The resulting differential Z-tilt angle is of about 130° between the two subgroups and not of 180°, meaning that the change of orientation of one subgroup of helices with respect to the other subgroup is accompanied by rotations around the x- and y-axes. It can also be seen that, exception made of H8, the fluctuations in the values of the global helix tilt angles along the whole trajectory are rather small, implying a structural stability of the entire helical bundle. For the **agonist-H3R** complex, the mean values of the X- and Y- tilt angles of helices TM1-7 are between 60 and 120° (Fig. S27abc to Fig. S34abc), with minor variations of the orientation of the helices with respect to the plane of the membrane. TM4 and TM6 are the most perpendicular to the plane of the membrane. As for the **apo** receptor, the values of the mean X- and Y-tilt angles are between 90 and 120° (Fig. S35abc to Fig. S42abc), like for the antagonist-H3R complex.

Thus, the average values of the X- and Y-tilt angles for the three systems are pretty much in the same range (60-120°), with rather small dispersions around each average value.

Global helix rotation

Fig. S19d to Fig. S26d show the values adopted by the global helix rotation during the MD simulation for TM1-7 and H8 of the **antagonist-H3R** complex. In general, a certain degree of fluctuation around the helix rotation average values is found for all helices. The average rotation for TM1-3, 5-7 is between 0 and ±20°. That of the eccentric TM4 is of about -55°. Besides, the rotation of the latter undergoes a shift from -55 to -70° in the last 2/5 of the trajectory (Fig. S22d). TM6 undergoes a clear-cut transition (Fig. S24d) from ~5 to 30° in rotation also in the last 2/5 of the trajectory, making of this a concerted movement. For the **agonist-H3R** complex, the helix rotation value of TM4 is rather of 0° (Fig. S30d), which is reminiscent of the behavior of the global helix tilt angles. For the **apo** receptor, helix rotation is again "well-behaved" for all TM helices except for TM4, for which it is of about -40° at the end of the trajectory (Fig. S38d).

Thus, the antagonist receptor and the apo form have in common a negative value for the average rotation of the eccentric TM4. For all three systems, the global helix rotation of all helices seems to be correlated.

Local helix tilt and turn angle per residue (overall winding-unwinding)

The local helix tilt is shown in Fig. S19e-Fig. S26e for each helix of the **antagonist-H3R** complex, in Fig. S27e-S34e for the first period of the **agonist-H3R** complex, and in Fig. S35e-S42e for the **apo** receptor. For all three systems, all TM helices show average values of 0-10° with little dispersion about these values; helix direction and fluctuations in direction are larger for H8.

The overall winding-unwinding of a helix can be monitored using the turn angle per residue (TPR) and measuring its deviation from ideal values of an α-helix (99.4°).[7] Fig. S19f-S26f show the TPR for TM1-7 and H8 of the **antagonist-H3R** complex. We can see that in all cases the average values fall close to the ideal values, but that the dispersion around this value is higher for TM6, indicating wider ranges of winding-unwinding of the helix. These findings point to TM6 as having peculiar mechanical properties that differentiate it from the other TM helices. For the **agonist-H3R** complex (Fig. S27f-S34f), the TPR values are close to the ideal value, but TM6, TM7 and H8 show large fluctuations skewed towards angles greater than 90°. Finally, for the **apo** receptor, the average values are close to 99° and the fluctuations around it are rather of small magnitude for all helices.

In conclusion, it is the agonist-H3R complex that shows larger oscillations of the TPR values as compared to the antagonist-H3R complex and the apo receptor (Fig. S35f- S42f).

##### Bend (Kink), Wobble and Face-shift angles

Figures S43, S44 and S45 show the values of the bend, the wobble and the face-shift angles around proline residues Pro 5.50, 6.50, and 7.50 of helices TM5-TM7 for the three systems. As we can see from Fig. S43a for the **antagonist-H3R** complex, the values of bend for TM5 undergo several transitions during the trajectory but oscillate around 40° starting at about 630 ns. This implies a value of 140° between the helical fragments on each side of the proline kink. The wobble angle for TM5 in Fig. S43b decreases from about -70° to -120° at 560 ns to end-up close to π (±180°). The helix is underwound as the face-shift angle is always positive, residing in the 25° to 75° range (Fig. S43c). It is worthwhile noticing that the face-shift angle undergoes transitions at around 350, 490 and 560 ns, i.e. at the same frames in the trajectory as the bend and wobble angles undergo transitions, indicating the correlated behavior of these variables.

After some oscillatory behavior in the 210-630 ns interval, the average bend angle of TM6 (Pro 6.50) of the antagonist-H3R complex (Fig. S43d) attains a value of about 35°, just like in the beginning of the trajectory. The wobble angle attains stable values around -85°; the face-shift values are stable about 52° starting at 420 ns (Fig. S43e and Fig. S43f, respectively), indicating an underwound helix. For TM7, the bend undergoes transitions in the 630-700 ns interval and adopts a value of 50° at the end of the trajectory (Fig. S43g). The wobble angle begins at -130° and finishes at -130°, having a few local transitions in the π region (Fig. S43h). Finally, the face-shift values indicate also an underwound helix –final value around 50° (Fig. S43i). The bend angle of section 1 of the trajectory of the **agonist-H3R** complex Pro 5.50 of the TM5 helix reaches a plateau at around 70°, before the HSM ligand escapes (Fig. S44a); the wobble operates a change between local average values around -130° et 100°, with shifts to the π region (Fig. S44b); the face-shift angles remain constant at around π (Fig. S44c). For Pro 6.50, the bend begins around 30°, goes through 15° and ends up in 30°, with incursions in the π region (Fig. S44d). The wobble stays in the π region (Fig. S44e). The face-shift is ±π, indicating again a tendency to unwind (Fig. S44f). The mean value of the bend angle for Pro 7.50 is about 80°, spanning a 20 to 150° range (Fig. S44g). The corresponding wobble angle remains essentially in the π region (Fig. S44h), just like the face-shift (Fig. S44i). For the **apo** receptor, the bend angle of Pro 5.50 fluctuates slightly around 10°; the wobble, with a mean negative value is such that the post-proline axis moves towards the proline Cα; the face-shift is close to 80°, indicating an underwinding of the helix (Fig. S45abc, respectively). For TM6, the bend angle fluctuates around 40°, the behavior of the wobble begins and ends around -60°, with variations in the -50 to -150° region, and the face-shift values begin and end-up in 75° (Fig. S45def, respectively). For Pro 7.50, the mean bend value is around 35°, with little fluctuations during the trajectory. The wobble presents values around -150°, and the face-shift values correspond to an underwound helix (50°), just like in TM5 and TM6 (Fig. S45ghi, respectively).

In summary, helices TM5 to TM7 of the agonist-H3R complex unwind, whereas the same helices in the antagonist-H3R complex and the apo receptor are underwound. In addition, the agonist-H3R complex shows large variations in the bend angle values of TM5 to TM7, consistent with the global toggle switch activation model,[27] in which a seesaw movement of TM6 around a pivot corresponding to Pro 6.50 takes place. On the contrary, the antagonist-H3R complex and the apo receptor show very narrow distributions about a given bend value. As for the wobble angle, we notice that all three helices have negative values for the antagonist-H3R complex; a similar behavior is observed for the apo receptor; whereas TM6 and TM7 of the agonist complex lean towards unwinding.

##### Total helix displacement (THD). Helix shift in the z-direction.

In the second part of the trajectory, the **antagonist-H3R** complex experiences an increase of THD for TM3 and H8, and a decrease for TM7 (Fig. S46a). The **agonist-H3R** complex shows an increase in the THD of H8 at the end of section 1 (Fig. S46b). Interestingly, the **apo** receptor shows a correlated change in THD for TM3 and TM5 in the 620-830 ns range (Fig. S46c). The largest value of the total helix displacement with respect to the average structure corresponds to TM4 (6 Å) of the **antagonist-H3R** complex (Fig. S46a).

The helix shift in the Z direction, perpendicular to the plane of the membrane of the last frame of the trajectory with respect to the average structure for the **antagonist-H3R** complex attains the highest shift upward for TM4 (7 Å) (Fig. S46a). Like TM4, TM2 is shifted upward, while the Z-displacement of the rest of the TM helices is negligible and that of H8 is downward. For the **agonist-H3R** complex (Fig. S46b), the helix shift falls in the (-1.5, 1.5) Å range. For the **apo** receptor (Fig. S46c), the absolute values of the Z-shift of the last frame are much smaller than the ligand complexes, oscillating between -1.5 and 1.0 Å, but this time with TM6 showing the largest downward shift (-1.5 Å). We observe a correlated shift around 750 ns between TMs 1, 2, 3, 4 and 7 (Fig. S46c).

These last results suggest that TM4, even though eccentric with respect to the helix bundle formed by TM1-TM3, TM5-TM7 may play a role in the inactivation mechanism of H3R through its total displacement on the plane of the membrane and its upward shift in the Z direction.

- Diffusion of a potassium monocation into the antagonist complex sodium allosteric site.

$K^+$ begins its course towards the interior of the receptor between TM5 and TM6 and is in contact with Leu 5.39 and His 67.00 (ECL3); it is hydrated at this time by five water molecules. A short interval after, it interacts with Ser 5.43, Glu 5.46, Phe 5.57 and Met 6.55 from TM5 and TM6, and is coordinated by only two to three water molecules. Towards the end of its path, it establishes contacts with Asp 2.50, Cys 3.36, Ser 3.39, Glu 5.46 and Trp 6.48 (TM2, TM3, TM5 and TM6), several of which are mediated by water, like with Asp and Glu residues, to finally become rehydrated by five water molecules. From time to time, the monocation is in contact with the highly conserved Asp 3.32 of the binding pocket. In the last frame of the trajectory, the cation is coordinated by Asp 2.50, Asp 3.32 and by three structured water molecules (r<3.0 Å), just like in the $\alpha$2A-AR.[18] As opposed to HSM, its insertion in the pocket is irreversible in the time scale studied (Fig. S47, S48).

FIGURES

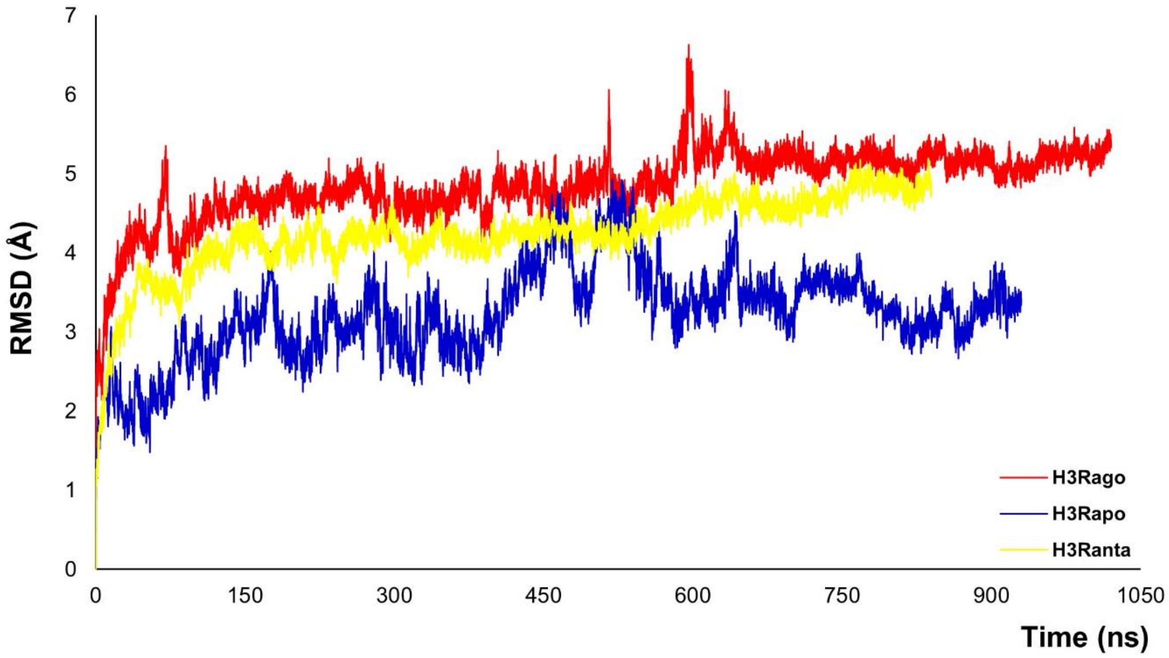


Fig. S1. Time-dependent Root Mean Square Deviation (RMSD) with respect to the initial structures for the antagonist (yellow), agonist (red) and apo (blue) structures.

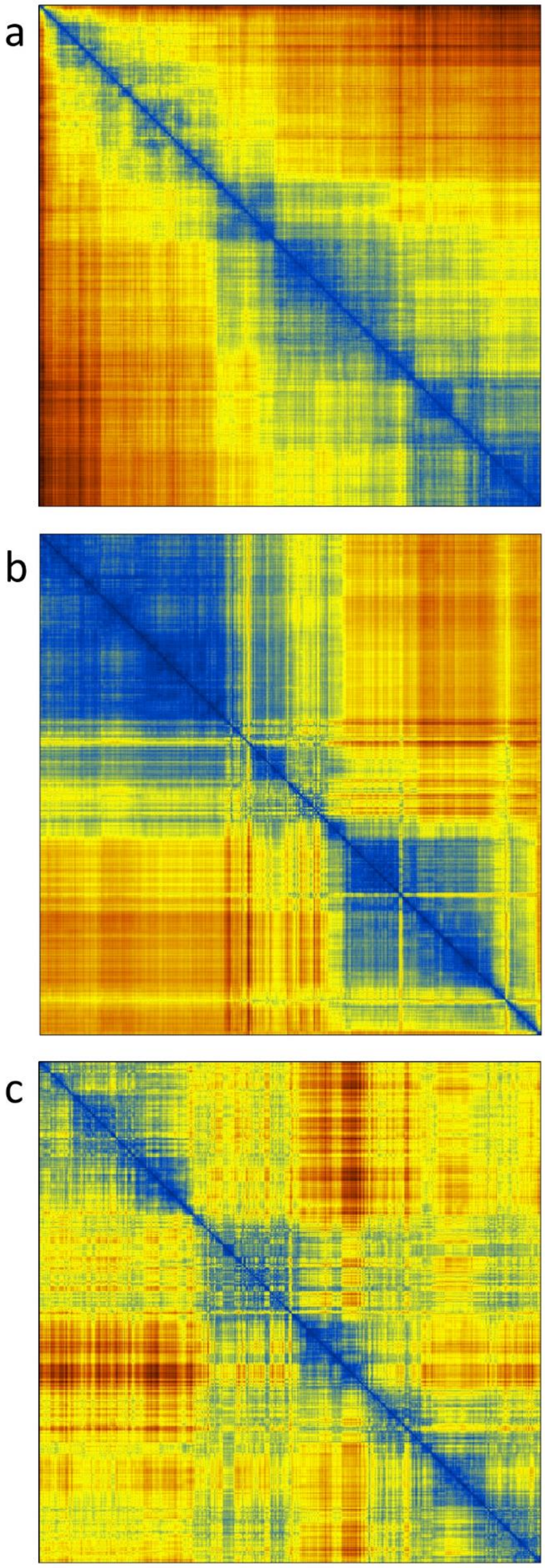


Fig. S2. The 2D RMSD matrix. Panels a) to c) correspond to the antagonist-complex, to the agonist-H3R complex, and to the apo receptor, respectively.

a

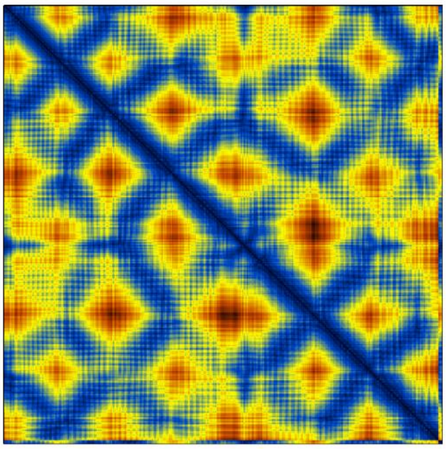

| TM | 1 | 2 | 3 | 4 | 5 | 6 | 7 |
|---|---|---|---|---|---|---|---|
| 1 | | x | | | | | x |
| 2 | | | x | | | x | x |
| 3 | | | | x | x | x | |
| 4 | | | | | | | |
| 5 | | | | | | x | |
| 6 | | | | | | | x |
| 7 | | | | | | | |

b

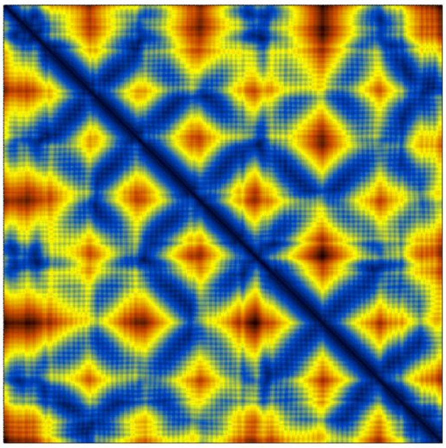

d

| TM | 1 | 2 | 3 | 4 | 5 | 6 | 7 |
|---|---|---|---|---|---|---|---|
| 1 | | x | | | | | x |
| 2 | | | x | x | | | x |
| 3 | | | | x | x | x | x |
| 4 | | | | | x | | |
| 5 | | | | | | x | |
| 6 | | | | | | | x |
| 7 | | | | | | | |

c

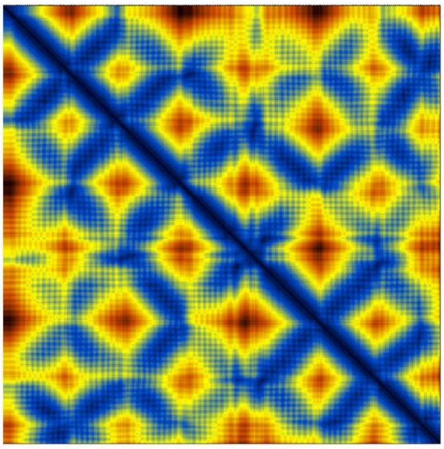

| TM | 1 | 2 | 3 | 4 | 5 | 6 | 7 |
|---|---|---|---|---|---|---|---|
| 1 | | x | | | | | x |
| 2 | | | x | x | | | x |
| 3 | | | | x | x | x | x |
| 4 | | | | | x | | |
| 5 | | | | | | x | |
| 6 | | | | | | | x |
| 7 | | | | | | | |

Fig. S3. The 2D contact maps. Panels a) to c) correspond to the antagonist- and agonist-H3R complexes, and to the apo receptor, respectively. Panel d) Relative positions of the TM helices in the form of a symmetric matrix as derived from panels a to c. A proximity between TM helices is represented by the presence of a matrix element.

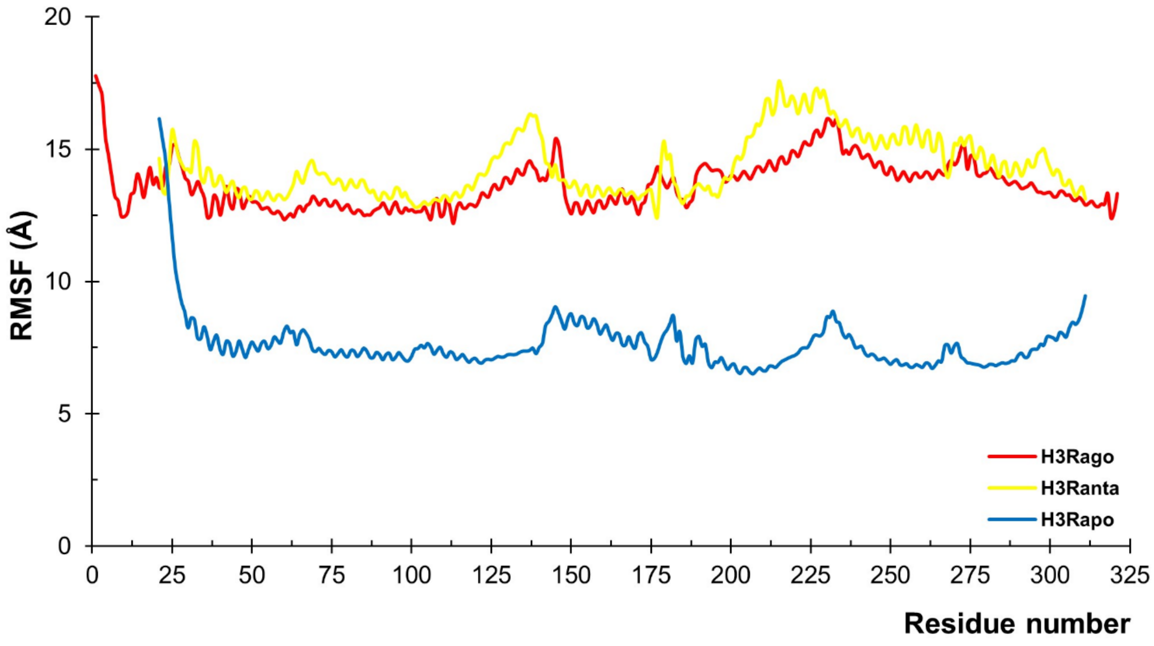


Fig. S4. Root mean square fluctuation (RMSF) of antagonist complex (yellow), agonist complex (red), and apo (blue) structures.

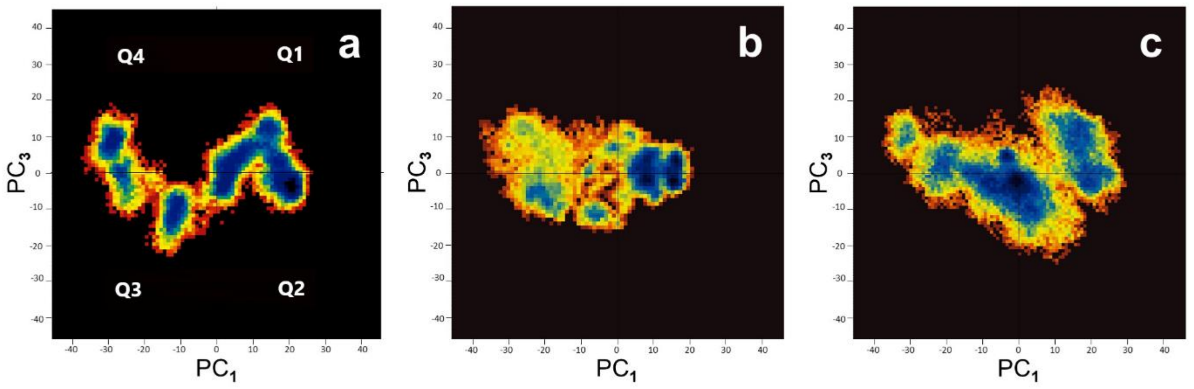


Fig. S5. Free energy landscape of the structures of the three systems identified by cartesian Principal Component Analysis. The panels a), b) and c) show a pseudo color representation of the distribution of the principal components PC1, PC3 obtained from a ~ 1 µs MD simulation for the antagonist, agonist and apo structures, respectively. Blue color represents energy wells (C1, C2, etc.). The map is divided into four quadrants (Q1-Q4).

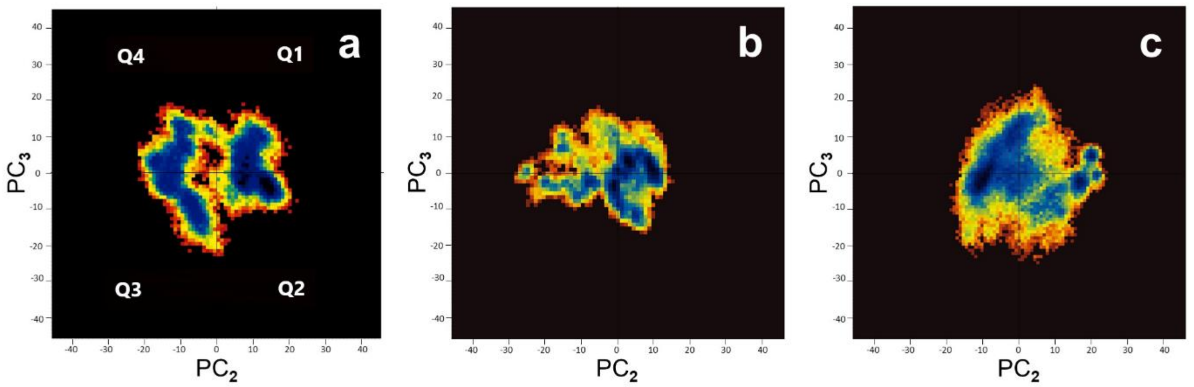


Fig. S6. Same as Fig. S5, but panels a), b) and c) correspond to the distribution of the principal components PC2, PC3 for the antagonist, agonist and apo structures, respectively.

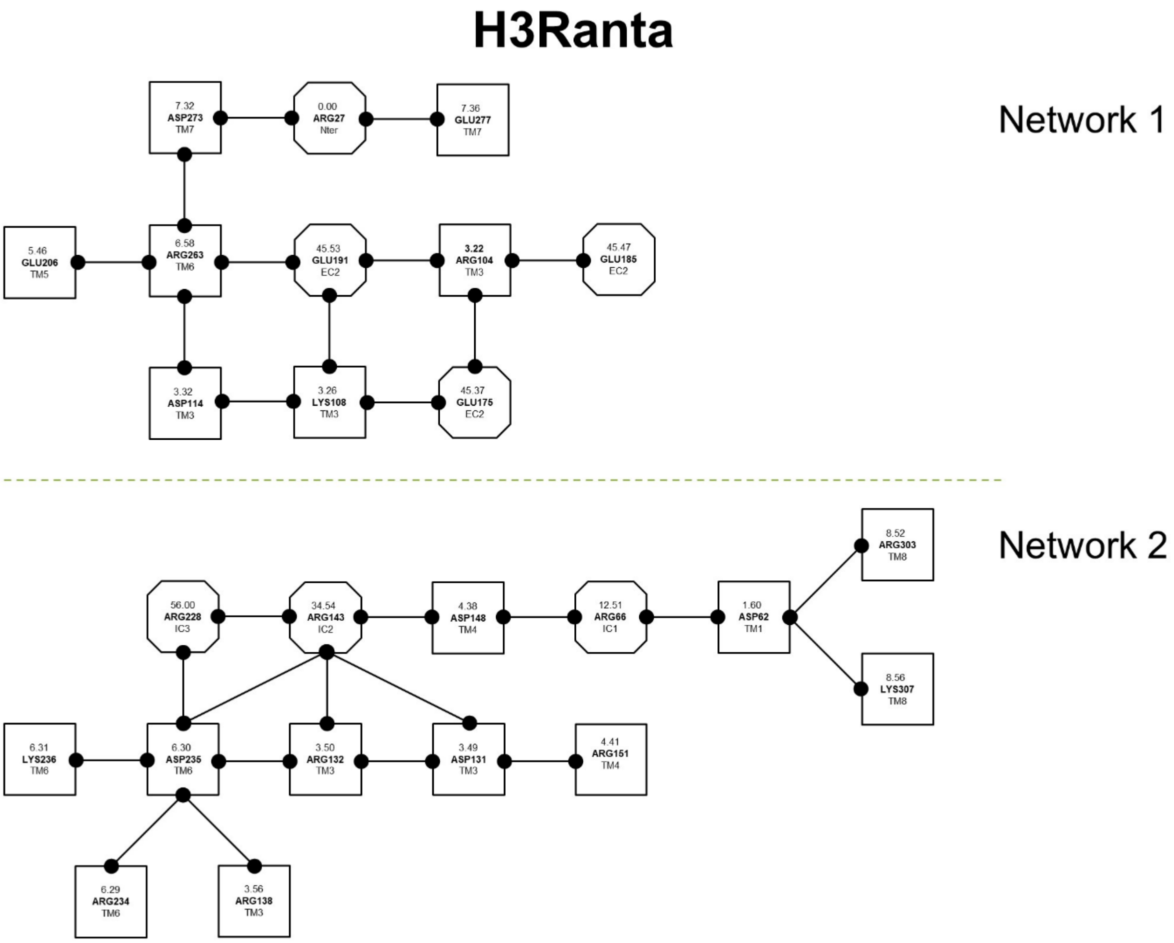

Fig. S7. Networks of charged residues for the antagonist-H3R complex. The two charged amino acids networks stabilize the structure of the receptor, important for the formation of an electrostatic clamp. The boxes correspond to the TM residues and the octagons to N-ter and loop residues.

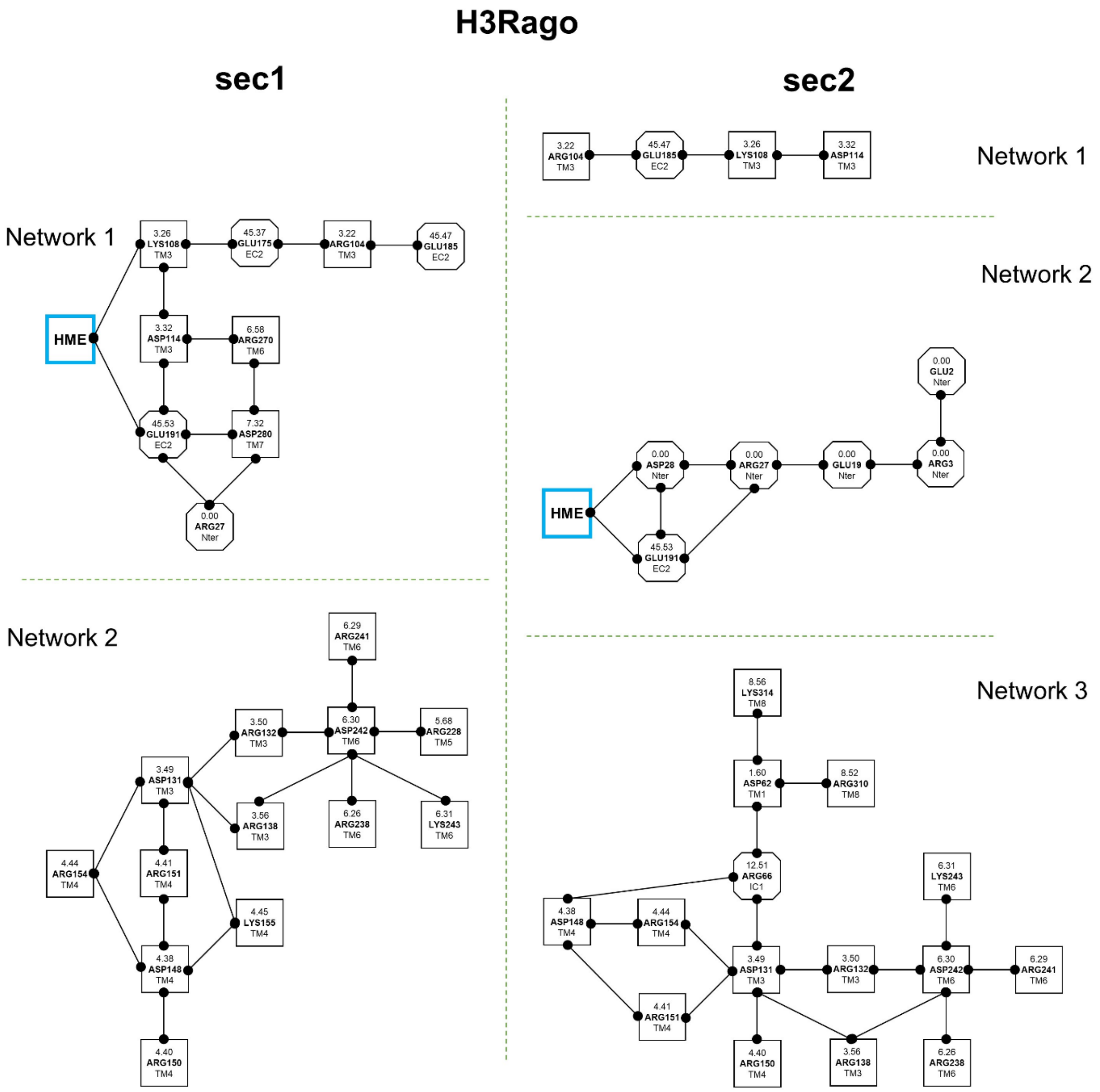


Fig. S8. Networks of charged residues for the agonist-H3R complex. In the two first sections at least one cluster interacts with the ligand. The first section in the trajectory shows two inner electrostatic networks, whereas the second section shows three. Shapes as in Fig. S7.

### H3Rapo

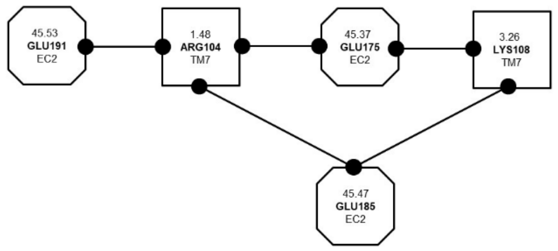


Network 1

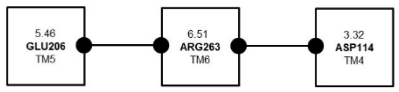


Network 2

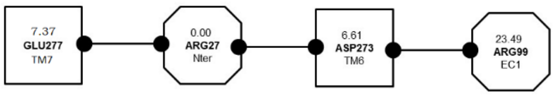


Network 3

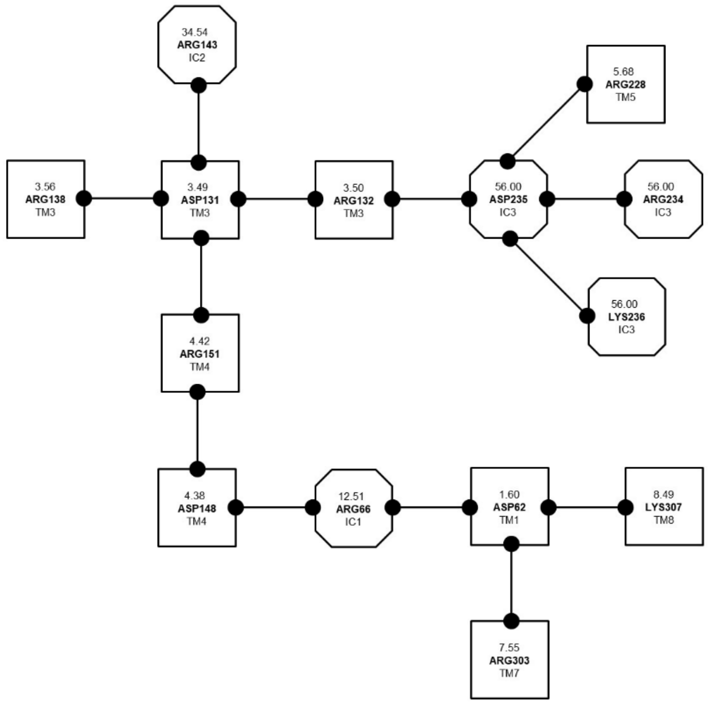


Network 4

Fig. S9. Networks of charged residues for the apo receptor. The four charged amino acids networks stabilize the apo structure. The first three networks involve less than six residues. Shapes as in Fig. S7.

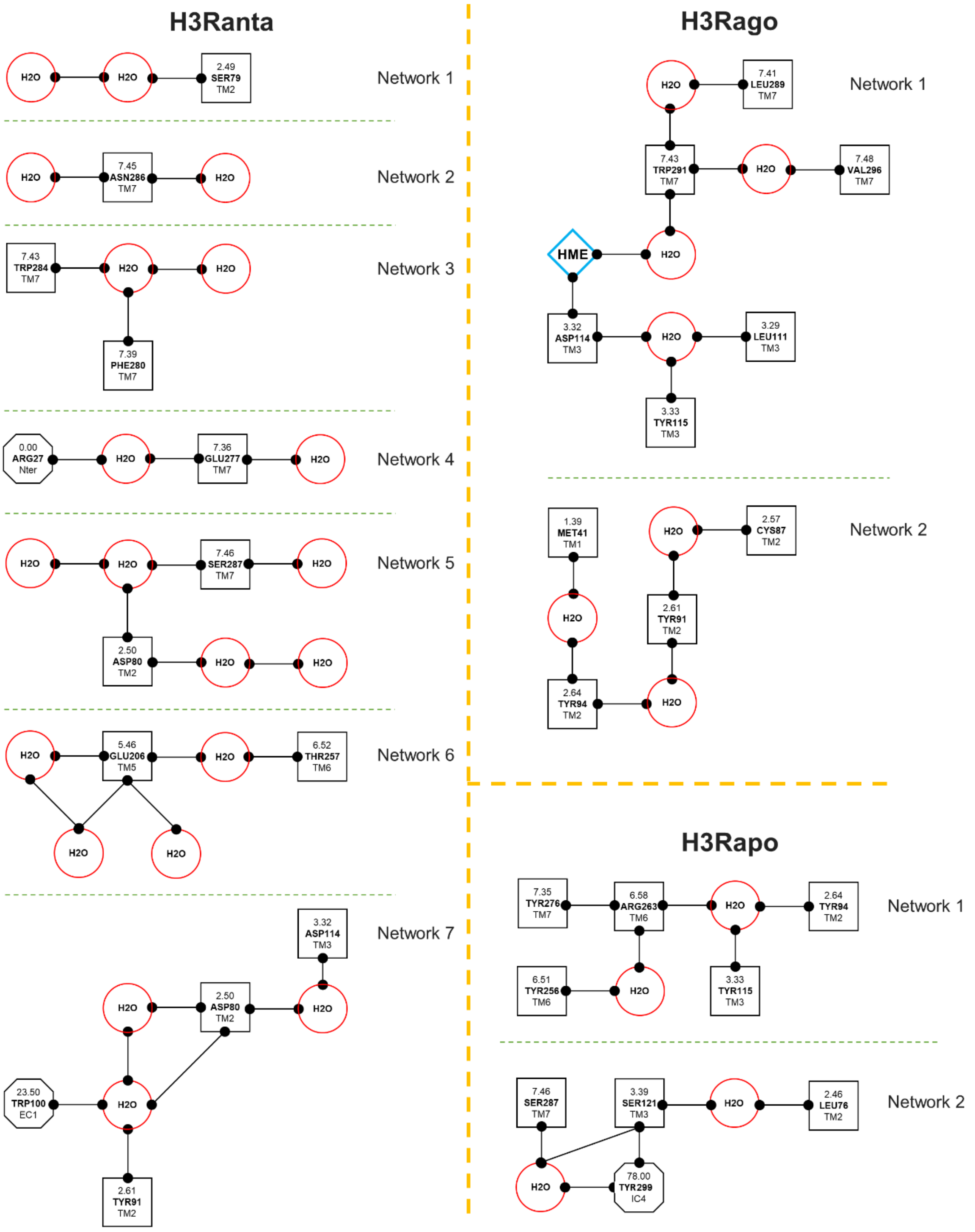

Fig. S10. H-bonded networks. a) Antagonist-H3R complex. The seven networks involve several internal water molecules and clusters, but not the CPX. b) Agonist-H3R complex. The first network shows a water molecule mediating the interaction between the HSM ligand and Trp7.43. c) Apo receptor. In all three systems, all but one are water-mediated interactions. Many of these interactions are polyvalent. The boxes correspond to the TM residues, the red circles to water molecules and the octagons to N-ter and loop residues.

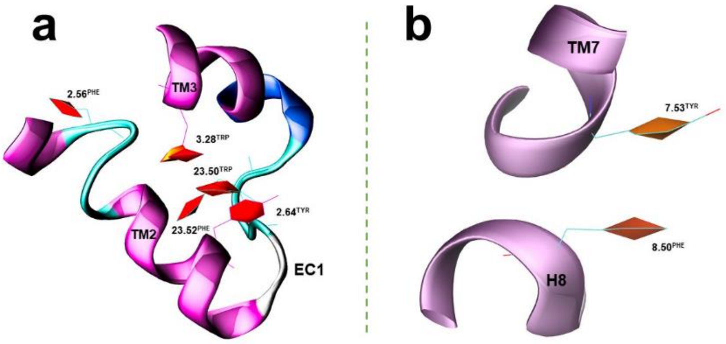


Fig. S11. Hydrophobic clusters formed by aromatic amino acids in the case of the antagonist-H3R complex. a) stacking interactions coming from TM2, TM3 and EC1; b) stacking interaction between two aromatic residues belonging to TM7 and H8.

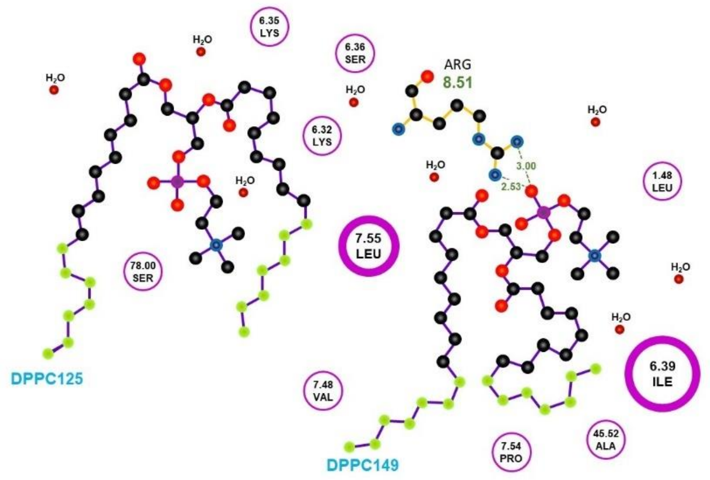


Fig. S12. 2D-binding major mode of DPPC lipids 125 and 149 to the H3 receptor in the antagonist state showing the interacting amino acid residues.

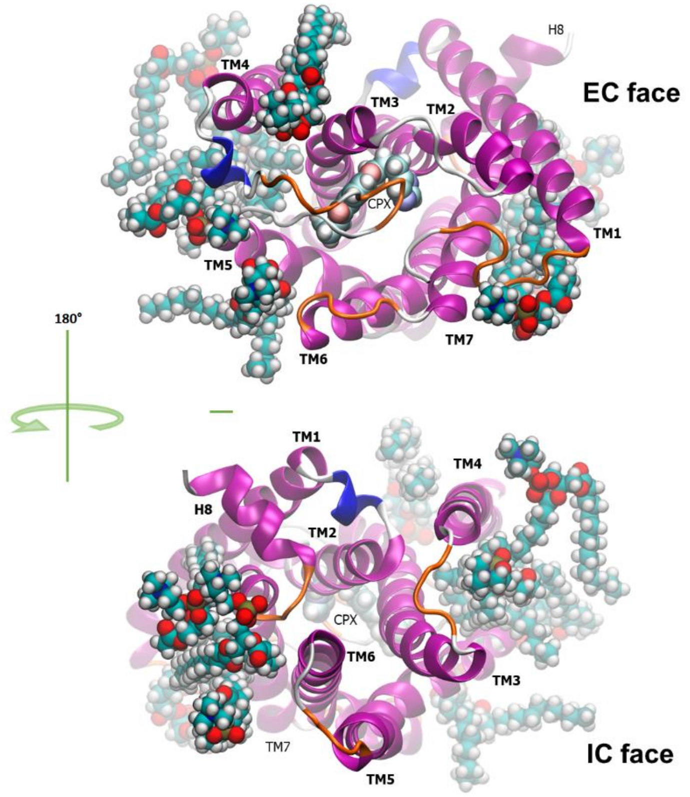

Fig. S13. 3D representation of the average binding mode of DPPC lipids to the H3 receptor in the antagonist state. Above, the extracellular face; below, the intracellular face. The lipids and the CPX ligand are visualized in CPK code.

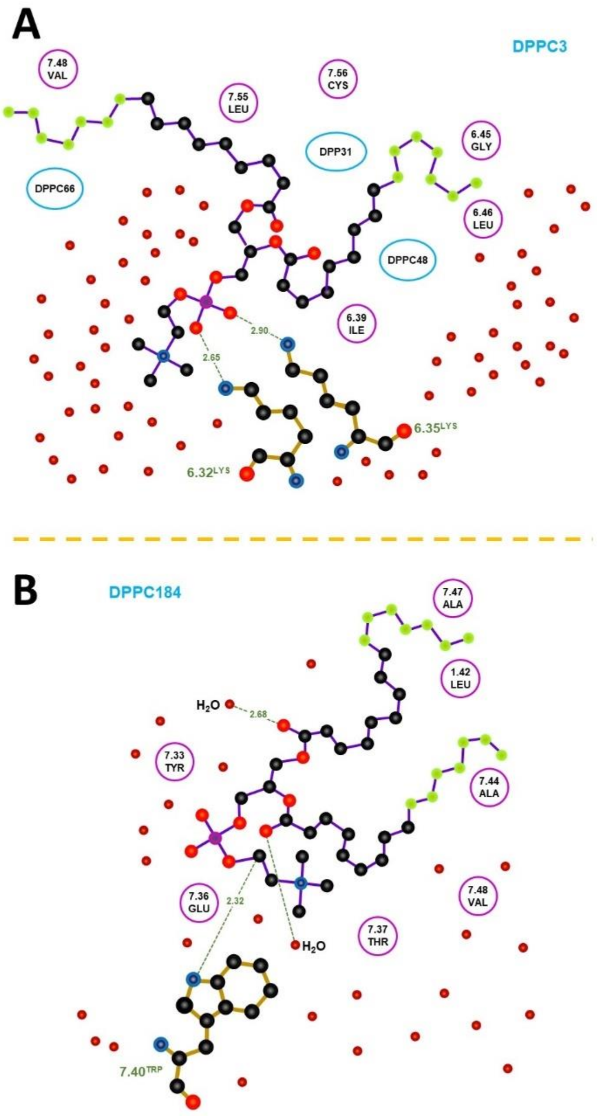


Fig. S14. 2D-binding major mode of DPPC lipids to the H3 receptor in the a) agonist, and b) apo state showing the interaction of amino acid residues and lipids. Multiple DPPC interactions can be observed. Several water molecules (in red) are present.

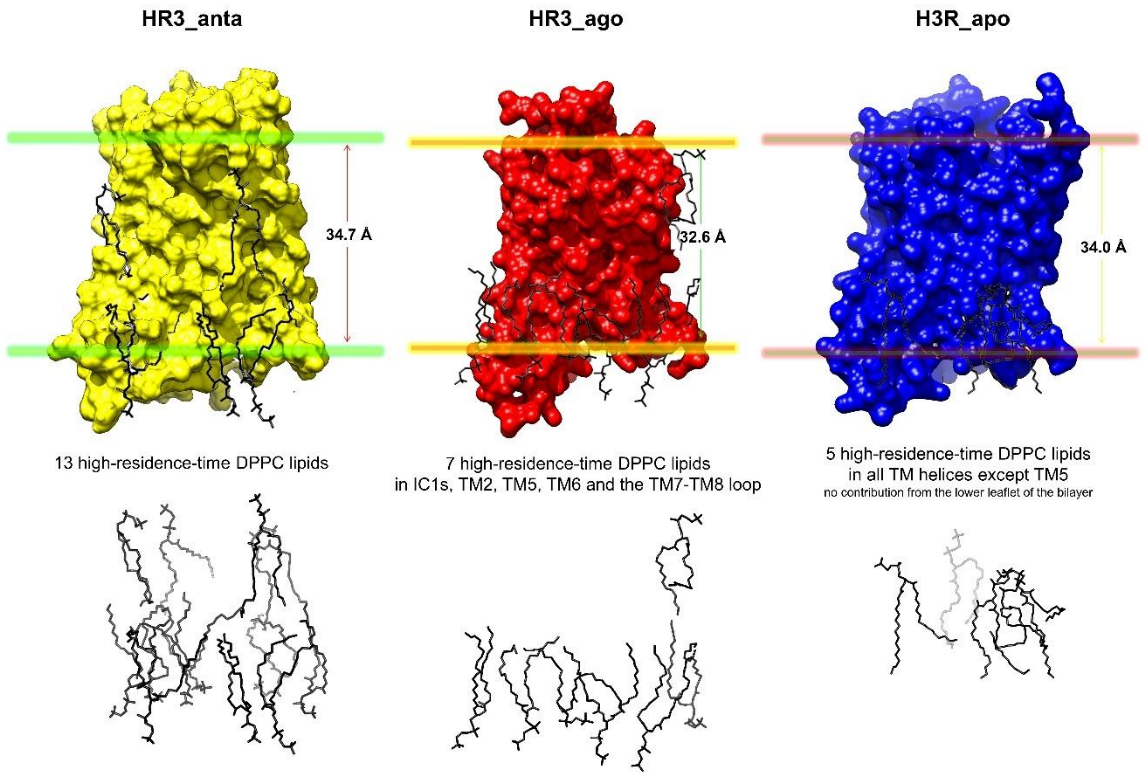


Fig. S15.Lipid-H3R interactions for the three systems. The images are representatives from the whole production phase of the simulation. The upper part shows the thickness of the lipid layer; the lower part view shows just the lipids without the H3R and allows to see the lipids in the backward part of the membrane.

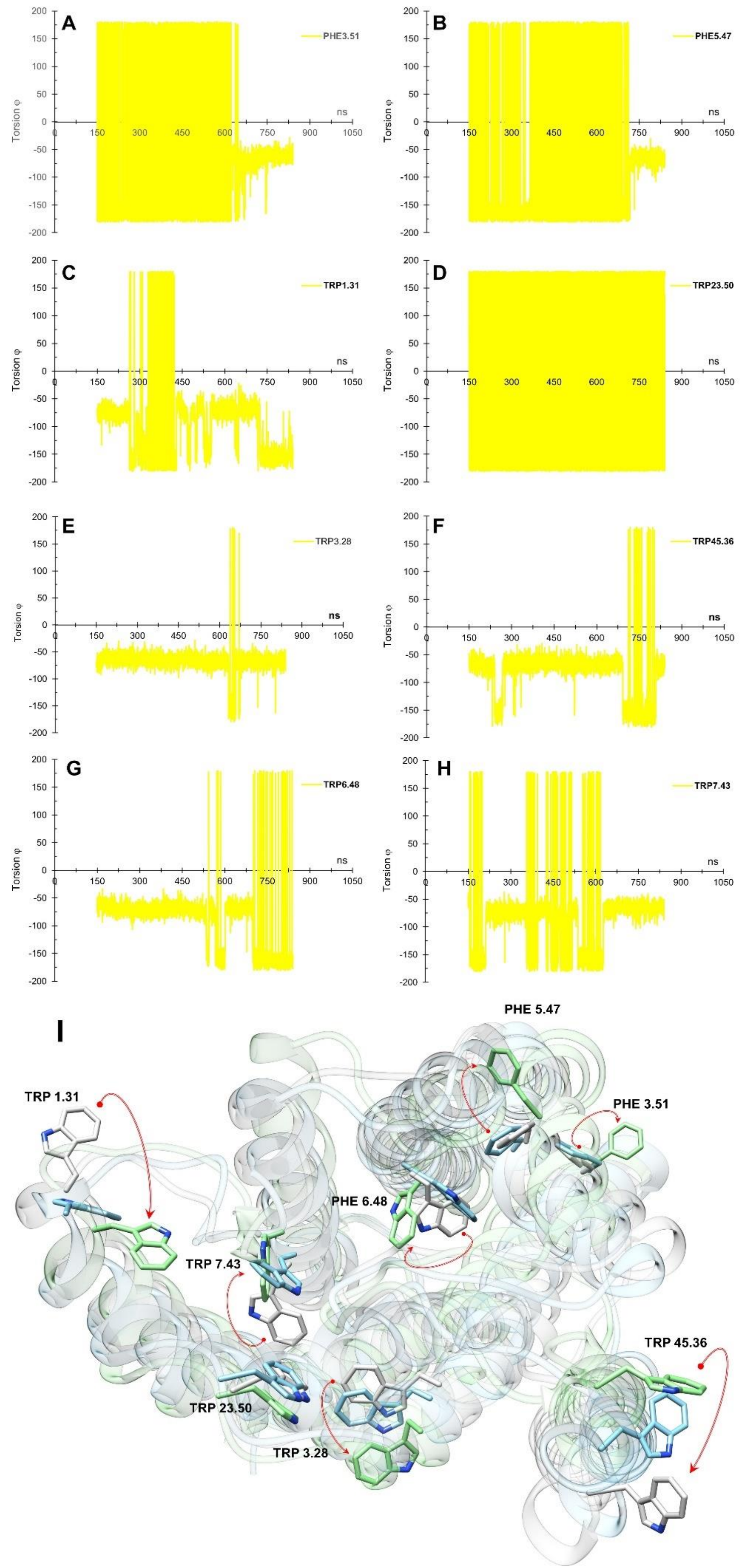
A
PHE3.51
B
PHE5.47
C
TRP1.31
D
TRP23.50
E
TRP3.28
F
TRP45.36
G
TRP6.48
H
TRP7.43
Torsion φ
ns
I
PHE 5.47
TRP 1.31
PHE 3.51
PHE 6.48
TRP 7.43
TRP 45.36
TRP 23.50
TRP 3.28

Fig. S16. χ1 torsion angle time evolution for the side chain of selected residues of the antagonist complex.

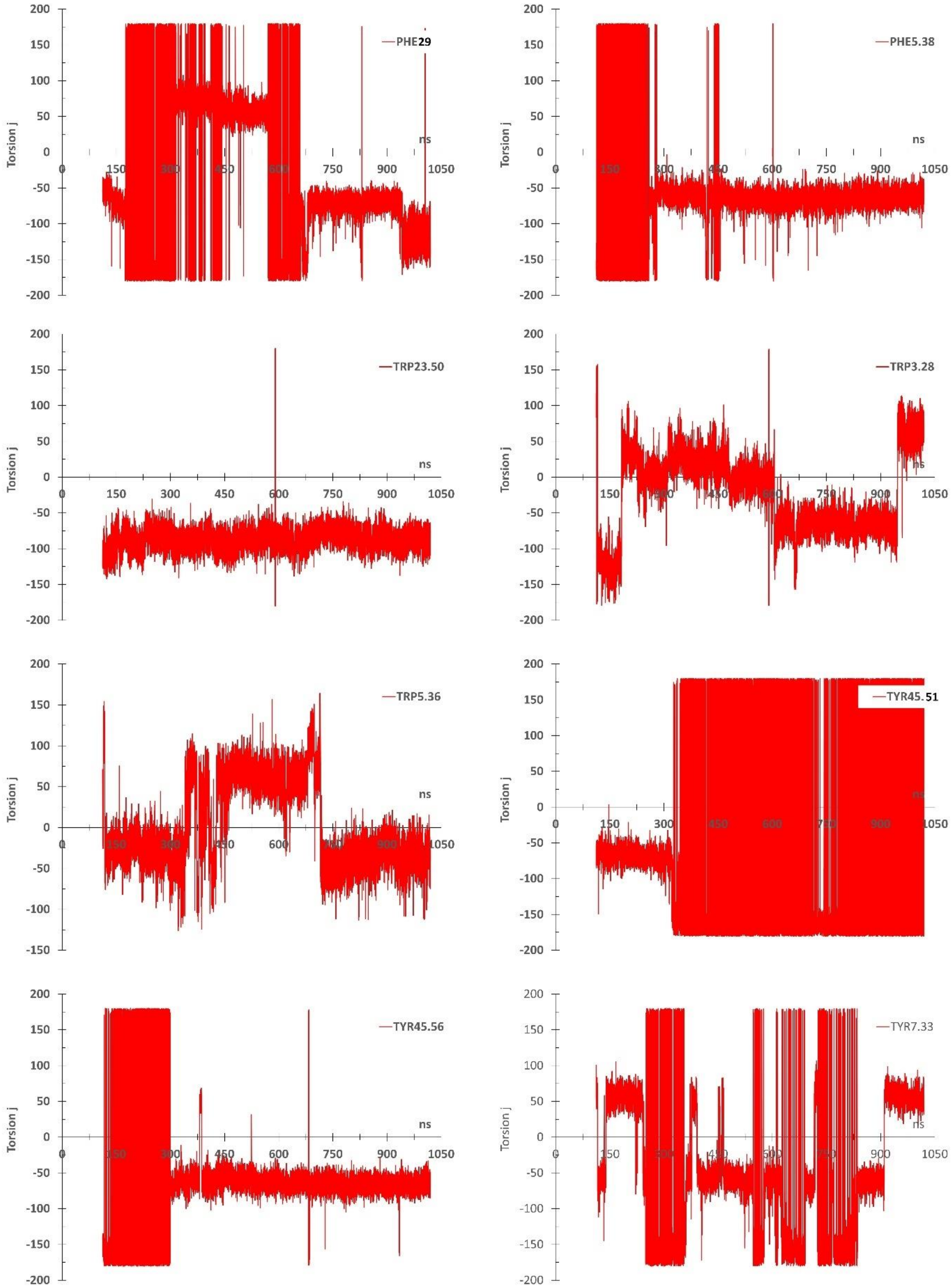

Fig. S17. $\chi 1$ torsion angle time evolution for the side chain of selected residues of the agonist complex.

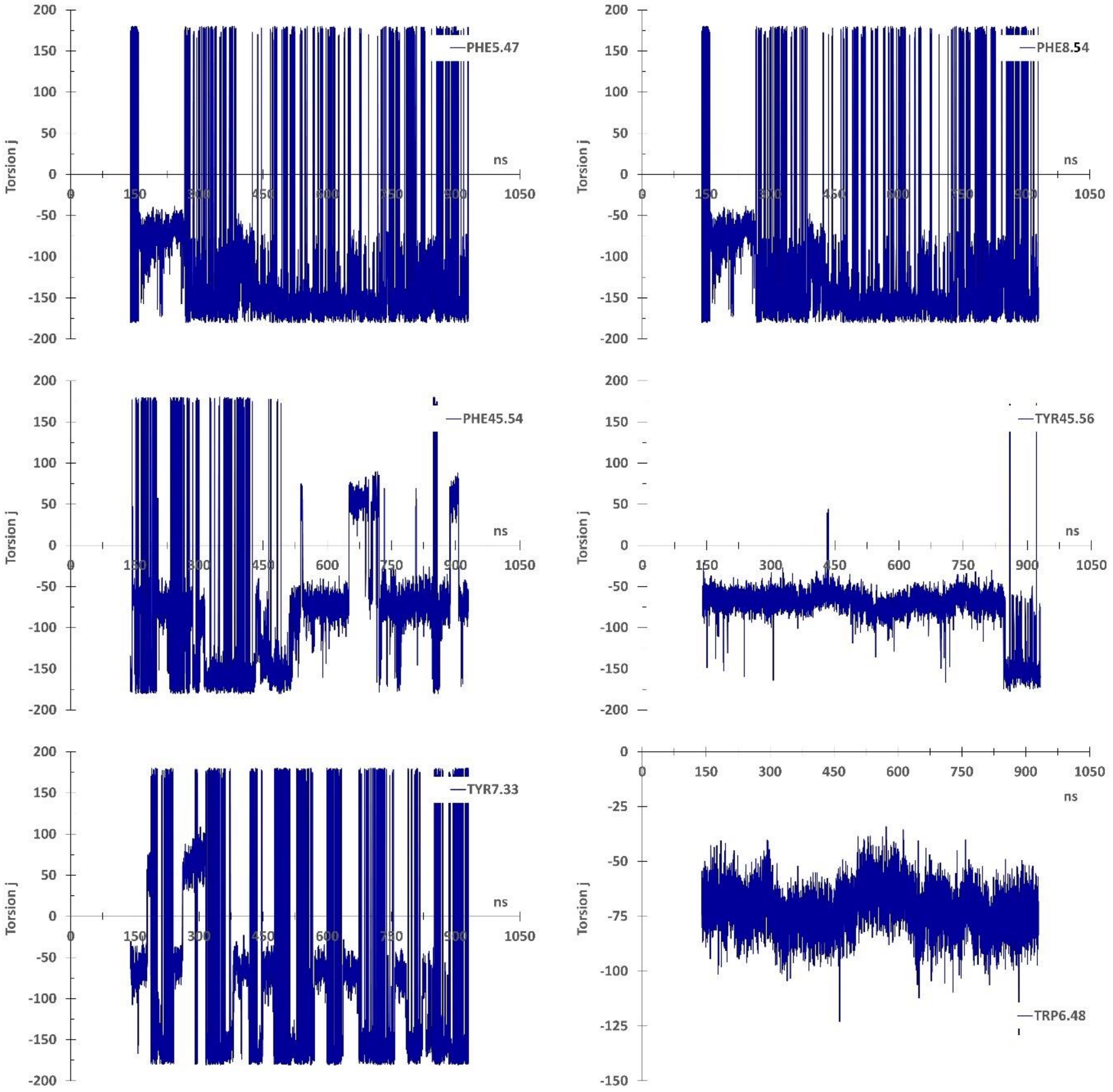


Fig. S18. $\chi_1$ torsion angle time evolution for the side chain of selected residues of the apo receptor.

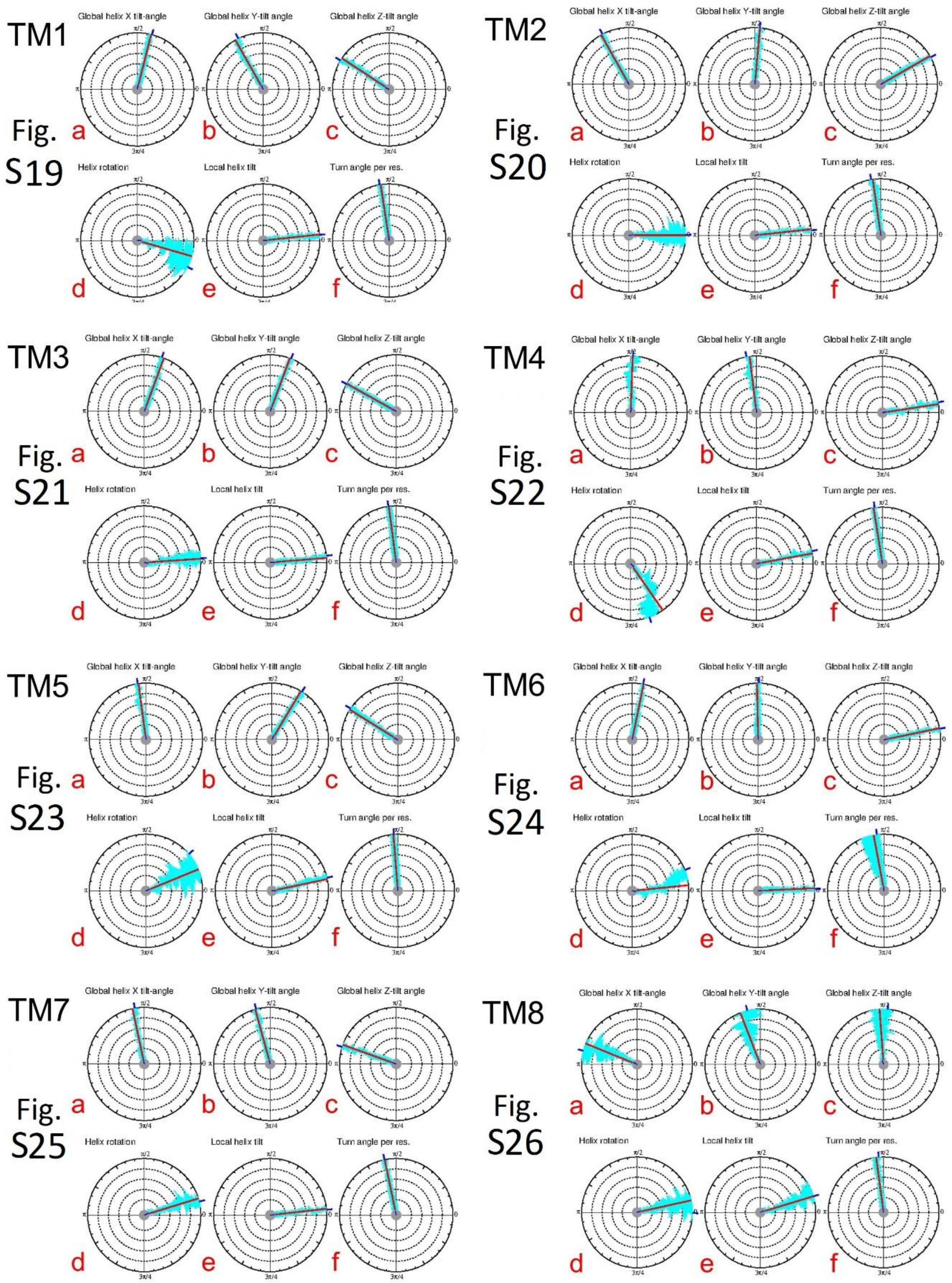


Fig. S19abc to S26abc. The global helix X-, Y- and Z-tilt angles for all eight helices of the antagonist-H3R complex, respectively.

Fig. S19d to S26d. The global helix rotations for all eight helices of the antagonist-H3R complex as a function of trajectory time, respectively.

Fig. S19e to S26e. The local helix tilt angles for all eight helices of the antagonist-H3R complex, respectively.

Fig. S19f to S26f. The turn angle per residue (TPR) for all eight helices of the antagonist-H3R complex, respectively.

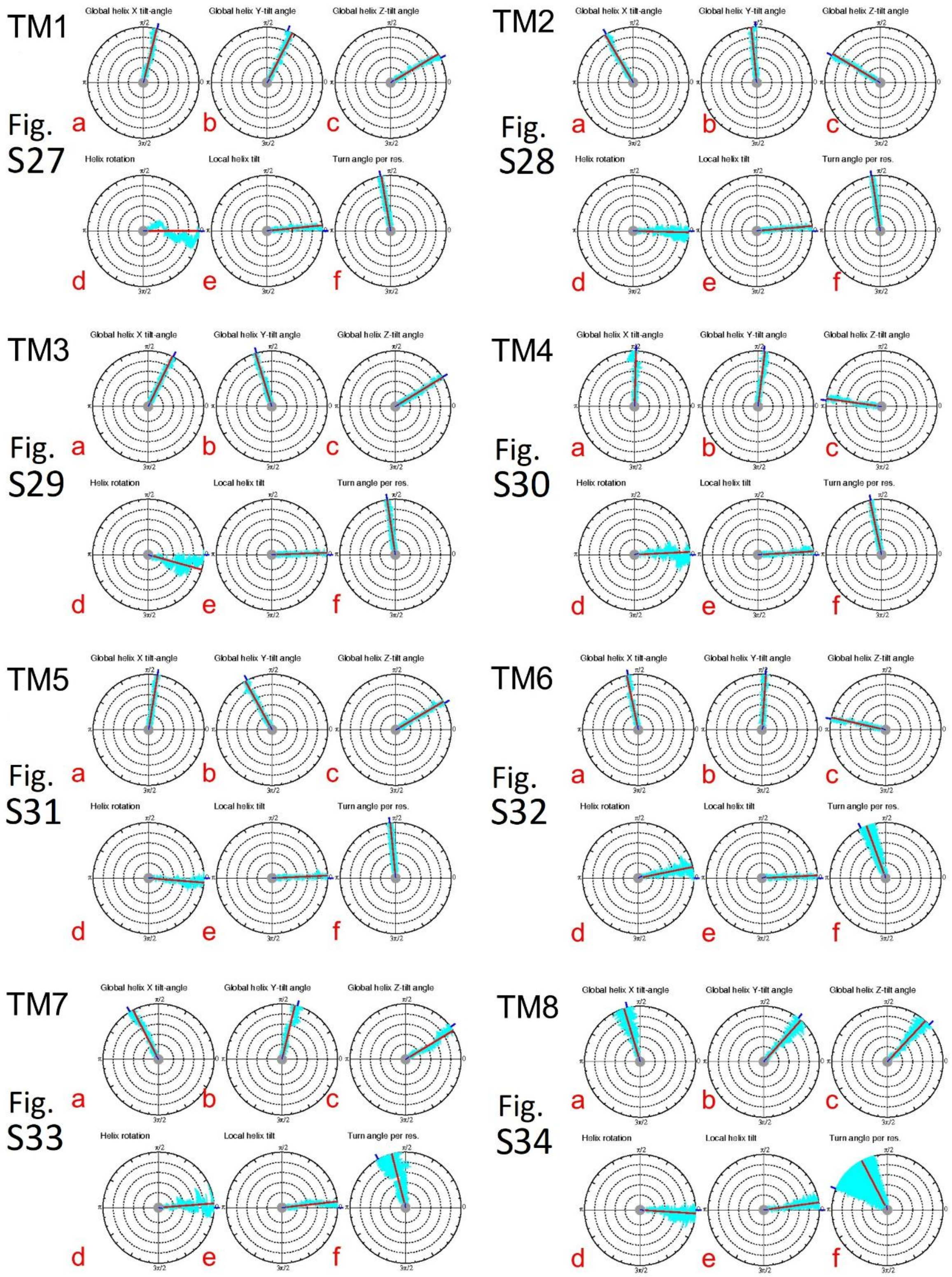


Fig. S27abc to S34abc. The global helix X-, Y- and Z-tilt angles for all eight helices of the agonist-H3R complex, respectively.

Fig. S27d to S34d. The global helix rotations for all eight helices of the agonist-H3R complex, respectively.

Fig. S27e to S34e. The local helix tilt angles for all eight helices of the agonist-H3R complex, respectively.

Fig. S27f to S34f. The turn angle per residue (TPR) for all eight helices of the agonist-H3R complex, respectively.

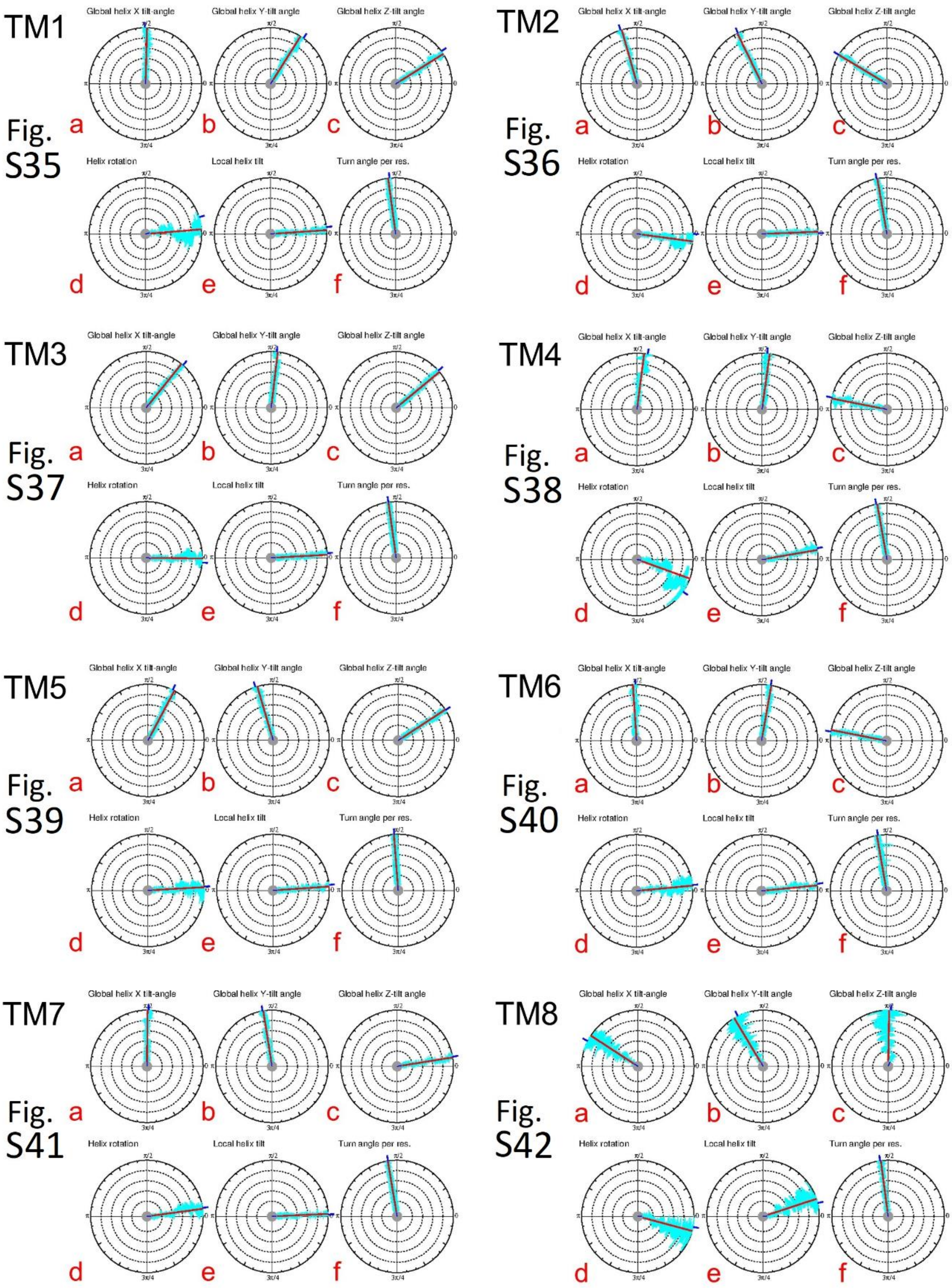


Fig. S35abc to S42abc. The global helix X-, Y- and Z-tilt angles for all eight helices of the apo receptor, respectively.

Fig. S35d to S42d. The global helix rotations for all eight helices of the apo receptor, respectively.

Fig. S35e to S42e. The local helix tilt angles for all eight helices of the apo receptor, respectively.

Fig. S35f to S42f. The turn angle per residue (TPR) for all eight helices of the apo receptor, respectively.

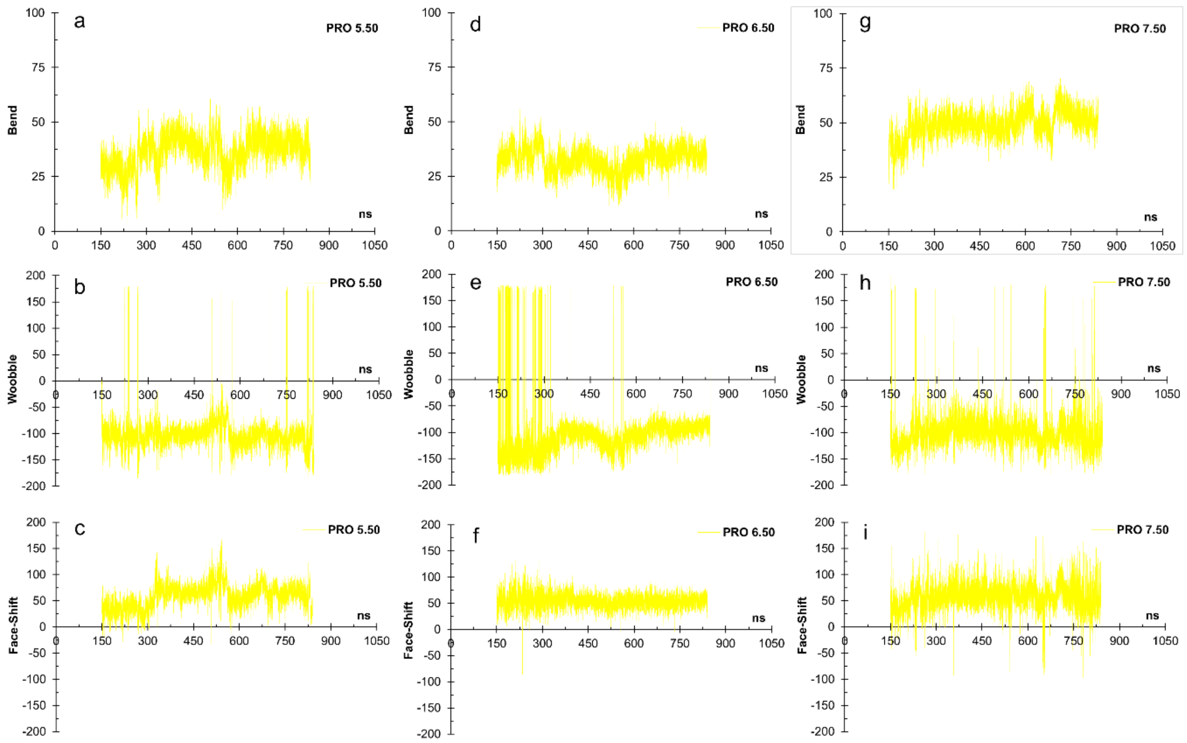


Fig. S43. Bend, wobble, and face-shift angles of the antagonist-H3R complex, around proline residues a) Pro 5.50 (TM5), b) Pro 6.50 (TM6), and c) Pro 7.50 (TM7), respectively.

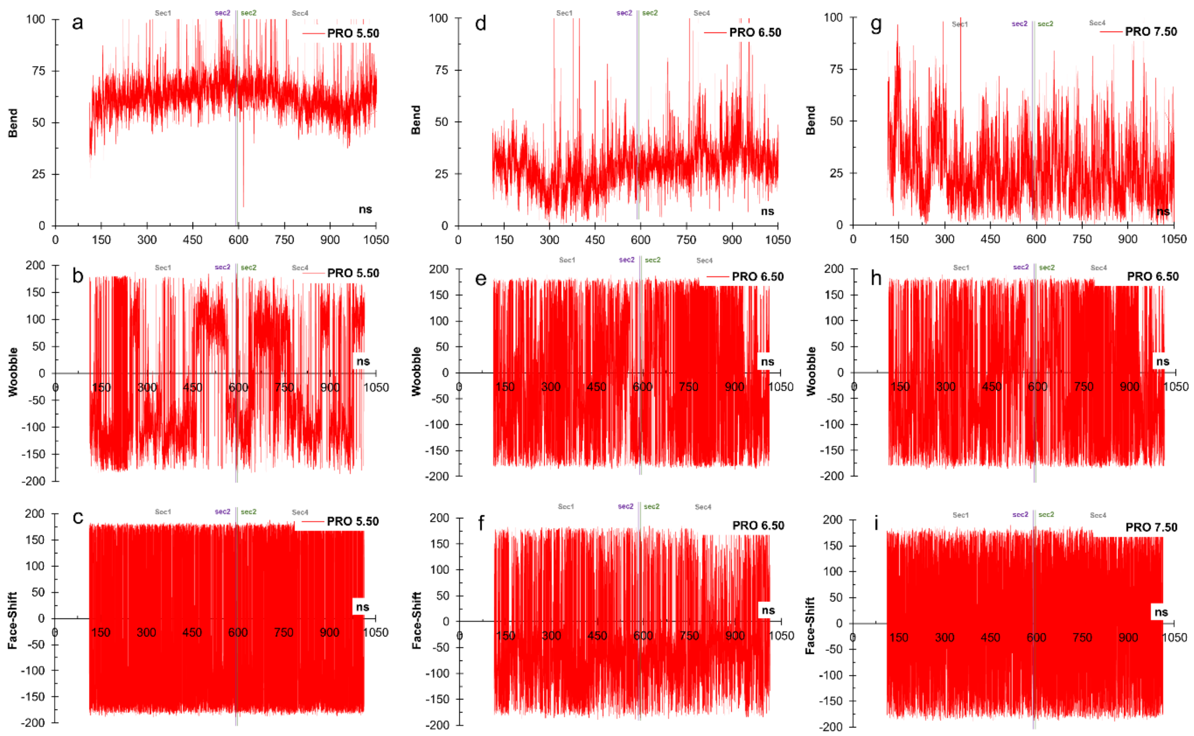


Fig. S44. Bend, wobble, and face-shift angles of the agonist-H3R complex, around proline residues a) Pro 5.50 (TM5), b) Pro 6.50 (TM6), and c) Pro 7.50 (TM7), respectively.

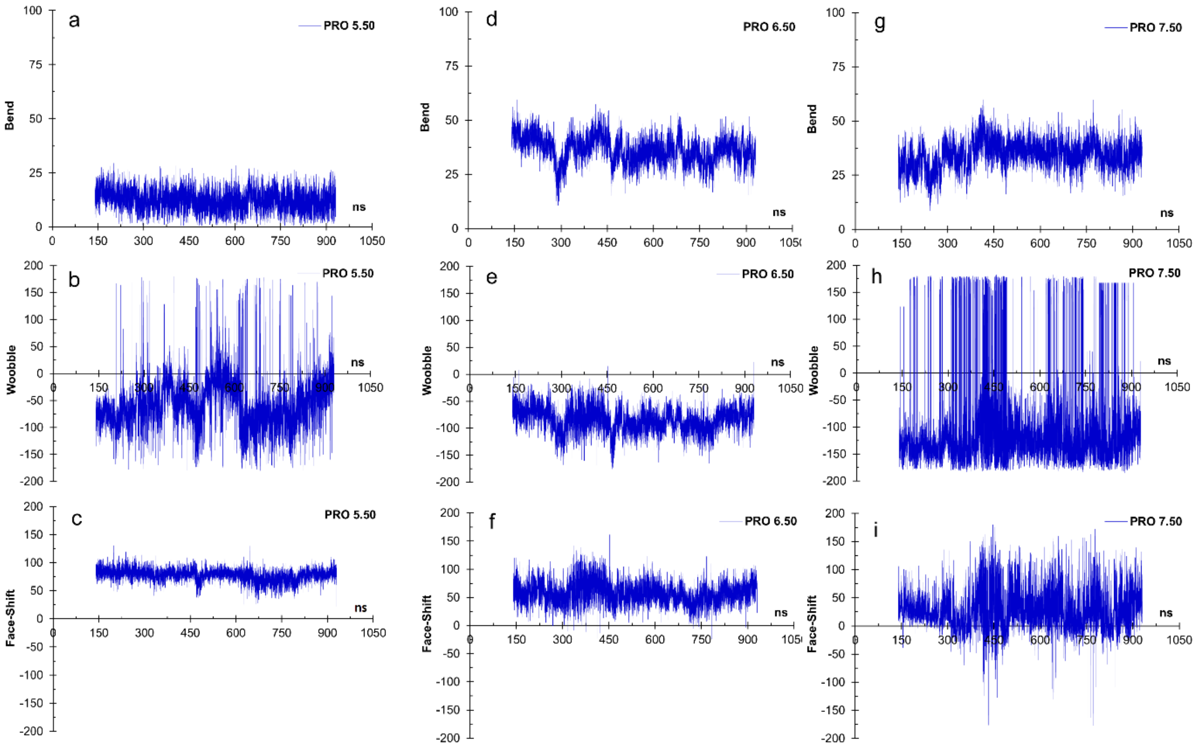

Fig. S45. Bend, wobble, and face-shift angles of the apo receptor, around proline residues a) Pro 5.50 (TM5), b) Pro 6.50 (TM6), and c) Pro 7.50 (TM7), respectively.

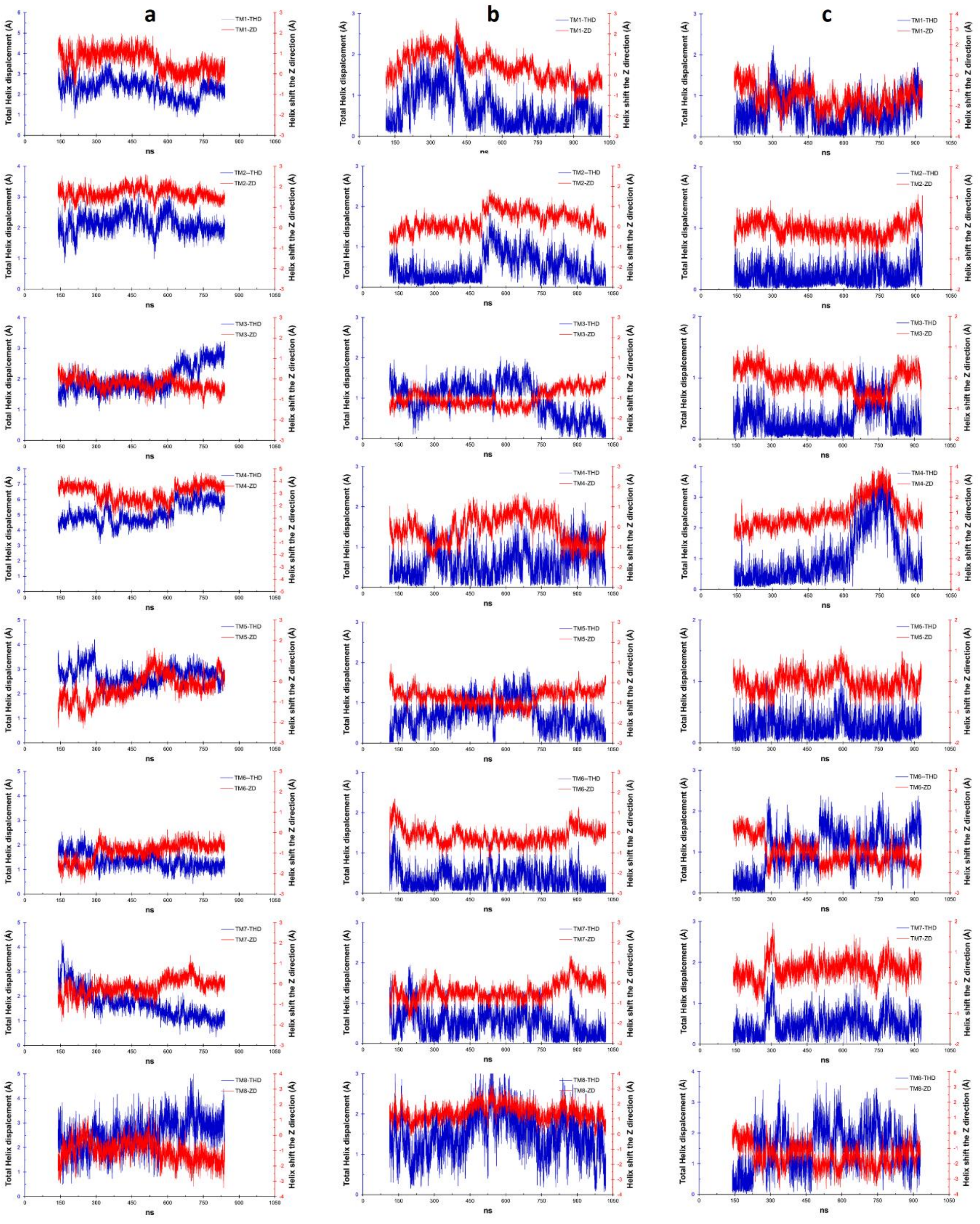

Fig. S46. Total helix displacement (in blue) and helix shift the Z-direction (in red) for a) the antagonist-H3R complex, b) the agonist-H3R complex and c) the apo receptor, respectively.

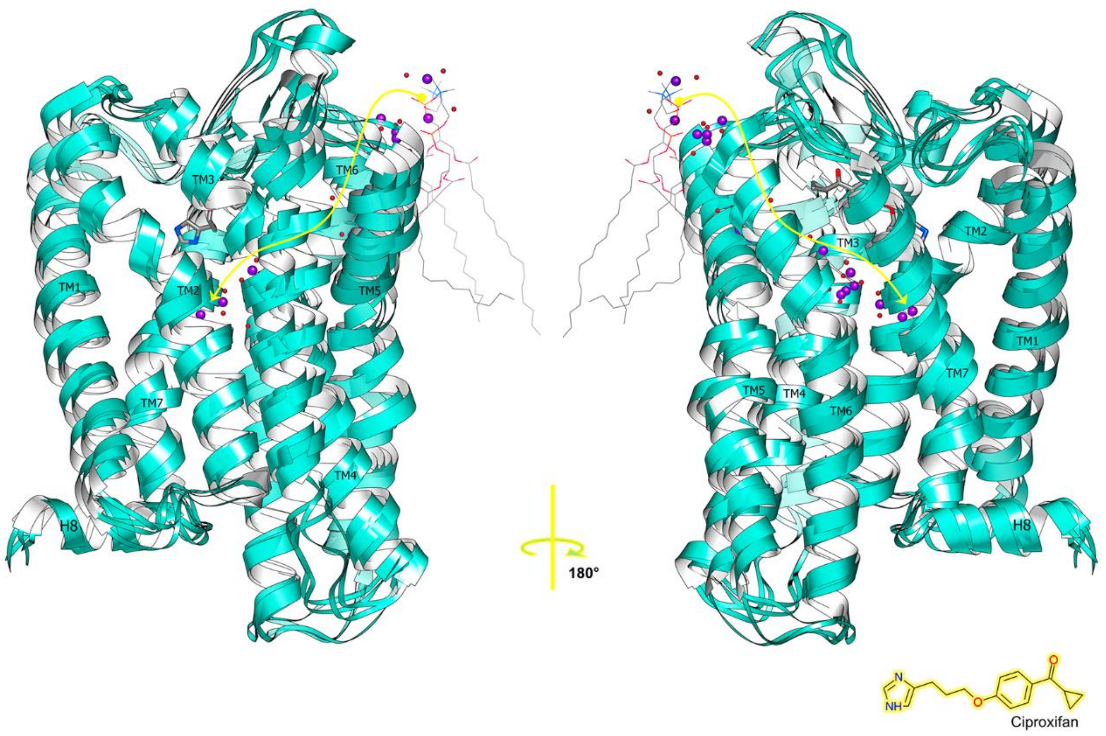


Fig. S47. Schematic illustration of the pathway taken by a $K^+$ metal ion (purple spheres) when penetrating the receptor from the extra-cytoplasmic space and binding to the $Na^+$ allosteric site. The curve in yellow denotes the binding pathway. Red spheres are water molecules and wire lines represent DPPC molecules.

Fig. S48. Video sequence of the spontaneous incorporation of the $K^+$ ion into the orthosteric site. (As a separate file).

### TABLES

**H3Ranta**

| | Hbond | | | | residencetime | Hbond | | | residencetime |
|---|---|---|---|---|---|---|---|---|---|
| | ARG104 | 3.22 | GLU175 | 45.37 | 100.00% | GLU191 | 45.53 | CPX | 47.37% |
| | ARG143 | 34.54 | ASP131 | 3.49 | 100.00% | PHE192 | 45.54 | CPX | 46.10% |
| | GLY186 | 45.48 | GLU191 | 45.32 | 100.00% | | | | |
| | ARG27 | 0.00 | GLU277 | 7.36 | 96.38% | | | | |
| * | THR278 | 7.37 | TYR274 | 7.33 | 91.76% | | | | |
| * | THR306 | 8.55 | ARG302 | 8.51 | 88.02% | | | | |
| * | ILE200 | 5.40 | TRP196 | 5.36 | 87.96% | | | | |
| * | THR201 | 5.41 | TYR197 | 5.37 | 78.94% | | | | |
| | ARG303 | 8.52 | ASP62 | 1.60 | 73.81% | | | | |
| | GLU185 | 45.47 | GLU191 | 45.53 | 73.63% | | | | |
| | TYR189 | 45.51 | GLU191 | 45.53 | 73.59% | | | | |
| * | PHE305 | 8.54 | PHE301 | 8.50 | 72.93% | | | | |
| | HSD298 | 78.00 | SER241 | 6.36 | 72.71% | | | | |
| * | MET41 | 1.39 | LEU37 | 1.35 | 71.89% | | | | |

**H3Rago-sec1**

| | Hbond | | | | residencetime | Hbond | | | residencetime |
|---|---|---|---|---|---|---|---|---|---|
| | ARG27 | 0.00 | GLU191 | 45.32 | 100.00% | HME | ASP114 | 3.32 | 48.07% |
| | ARG310 | 8.52 | ASP62 | 1.60 | 98.50% | | | | |
| * | SER254 | 6.42 | ALA250 | 6.38 | 84.78% | | | | |
| | TRP174 | 45.36 | PRO169 | 4.59 | 81.34% | | | | |
| | ARG310 | 8.52 | TYR306 | 78.00 | 81.32% | | | | |
| | LYS108 | 3.26 | GLU175 | 45.37 | 77.41% | | | | |
| * | ASP80 | 2.50 | LEU76 | 2.46 | 72.77% | | | | |
| * | LYS247 | 6.34 | LYS243 | 6.31 | 71.46% | | | | |
| * | PHE312 | 8.54 | PHE308 | 8.50 | 70.75% | | | | |
| * | THR313 | 8.55 | ARG309 | 8.51 | 70.16% | | | | |

**H3Rago-sec2**

| | | | | | | | | |
|---|---|---|---|---|---|---|---|---|
| ARG3 | 0.00 | GLU19 | 0.00 | 100.00% | HME | ASP280 | 7.32 | 44.37% |
| ARG310 | 8.52 | TYR306 | 78.00 | 100.00% | | | | |
| PHE256 | 6.44 | ILE252 | 6.40 | 99.48% | | | | |
| SER254 | 6.42 | ALA250 | 6.38 | 93.59% | | | | |
| LYS247 | 6.35 | LYS243 | 6.31 | 90.47% | | | | |
| TRP174 | 45.36 | PRO169 | 4.59 | 89.43% | | | | |
| LEU239 | 6.27 | GLN235 | 56.00 | 88.39% | | | | |
| ALA157 | 4.47 | VAL153 | 4.43 | 85.27% | | | | |
| ARG27 | 0.00 | GLU191 | 45.53 | 83.54% | | | | |
| SER30 | 0.00 | GLU284 | 7.36 | 83.19% | | | | |
| ARG228 | 5.68 | GLN235 | 56.00 | 75.91% | | | | |
| ARG104 | 3.22 | GLU185 | 45.47 | 74.87% | | | | |
| PHE86 | 2.56 | LEU82 | 2.52 | 74.52% | | | | |
| THR229 | 5.69 | ILE225 | 5.65 | 74.52% | | | | |
| ASP80 | 2.50 | LEU76 | 2.46 | 74.18% | | | | |
| PHE312 | 8.54 | PHE308 | 8.50 | 73.83% | | | | |
| LYS108 | 3.26 | GLU175 | 45.37 | 71.06% | | | | |
| ALA250 | 6.38 | ALA246 | 6.34 | 70.36% | | | | |

**H3Rago-sec3**

| | | | | |
|---|---|---|---|---|
| SER30 | 0.00 | GLU284 | 7.36 | 100.00% |
| SER254 | 6.42 | ALA250 | 6.38 | 99.39% |
| LYS247 | 6.35 | LYS243 | 6.31 | 98.99% |
| PHE256 | 6.44 | ILE252 | 6.40 | 96.96% |
| LEU239 | 6.27 | GLN235 | 56.00 | 95.33% |
| ARG310 | 8.52 | TYR306 | 78.00 | 91.68% |
| SER248 | 6.36 | LYS244 | 6.32 | 87.63% |
| TRP174 | 45.36 | PRO169 | 4.59 | 86.41% |
| ARG27 | 0.00 | GLU191 | 45.53 | 85.19% |
| ASP80 | 2.50 | LEU76 | 2.46 | 82.76% |
| THR313 | 8.55 | ARG309 | 8.51 | 80.73% |
| ALA157 | 4.47 | VAL153 | 4.43 | 80.73% |
| TYR91 | 2.61 | CYS87 | 2.57 | 80.32% |
| ARG228 | 5.68 | GLN235 | 56.00 | 80.32% |
| LYS108 | 3.26 | ALA170 | 4.60 | 75.05% |

**H3Rago-sec4**

| | | | | |
|---|---|---|---|---|
| ARG3 | 0.00 | GLU191 | 45.53 | 100.00% |
| SER254 | 6.42 | ALA250 | 6.38 | 88.07% |
| ASP80 | 2.50 | LEU76 | 2.46 | 81.14% |
| ARG310 | 8.52 | TYR306 | 78.00 | 78.73% |
| TRP174 | 45.36 | PRO169 | 4.59 | 77.59% |
| PHE312 | 3.50 | PHE308 | 8.50 | 75.65% |

**H3RApo**

| | Hbond | | | | residencetime |
|---|---|---|---|---|---|
| * | SER247 | 6.42 | ALA243 | 6.38 | 86.97% |
| * | SER79 | 2.49 | ASN75 | 2.45 | 86.51% |
| * | ILE78 | 2.48 | LEU74 | 2.44 | 85.91% |
| * | VAL246 | 6.41 | LEU242 | 6.37 | 81.37% |
| | TRP174 | 45.36 | PRO169 | 4.59 | 78.15% |

Table S1. H-bonding residue pairs for residence times greater than 70% for the antagonist, agonist and apo systems, respectively. Includes intra-helical H-bonds (*), except sections 2-4 of the agonist complex. HSD stands for His. Includes main-chain and side-chain H-bonds.

**N-ter**

| H3Ranta | H3Rago | H3Rapo |
|---|---|---|

**TM1**

| H3Ranta | H3Rago | H3Rapo |
|---|---|---|
| | MET41 1.39 | |
| | LEU44 1.42 | |

**ICL1**

| H3Ranta | H3Rago | H3Rapo |
|---|---|---|

**TM2**

| H3Ranta | H3Rago | H3Rapo |
|---|---|---|
| | ASN70 2.4 | |
| SER79 2.49 | | SER79 2.49 |
| ASP80 2.5 | | |
| PHE81 2.51 | | |
| VAL83 2.53 | VAL83 2.53 | VAL83 2.53 |
| | CYS87 2.57 | |
| ILE88 2.58 | ILE88 2.58 | ILE88 2.58 |
| | LEU90 2.6 | |
| TYR91 2.61 | TYR91 2.61 | TYR91 2.61 |
| TYR94 2.64 | TYR94 2.64 | |

**ECL1**

| H3Ranta | H3Rago | H3Rapo |
|---|---|---|

| | No. of waters |
|---|---|
| H3Rapo | 48.9516 |
| H3Ranta | 61.3404 |
| H3Rago | |
| sec1 | 35.1545 |
| sec2 | 19.8312 |
| sec3 | 19.0037 |
| sec4 | 24.9105 |

**TM3**

| H3Ranta | H3Rago | H3Rapo |
|---|---|---|
| CYS107 3.25 | CYS107 3.25 | |
| TRP110 3.28 | | |
| LEU11 3.29 | LEU111 3.29 | LEU111 3.29 |
| ASP114 3.32 | ASP114 3.32 | ASP114 3.32 |
| TYR115 3.33 | TYR115 3.33 | TYR115 3.33 |
| LEU117 3.35 | | LEU117 3.35 |
| CYS118 3.36 | | |
| SER121 3.39 | | SER121 3.39 |
| VAL122 3.4 | | VAL122 3.4 |

**ICL2**

| H3Ranta | H3Rago | H3Rapo |
|---|---|---|

**TM4**

| H3Ranta | H3Rago | H3Rapo |
|---|---|---|

**ECL2**

| H3Ranta | H3Rago | H3Rapo |
|---|---|---|
| | GLU175 45.37 | |
| | TYR189 45.51 | |
| | ALA190 45.52 | |
| GLU191 45.53 | GLU191 45.53 | |
| PHE192 45.54 | PHE192 45.54 | PHE192 45.54 |
| PHE193 45.55 | | |
| TYR194 45.56 | | TYR194 45.56 |

**TM5**

| H3Ranta | H3Rago | H3Rapo |
|---|---|---|
| | PHE198 5.38 | |
| LEU199 5.39 | | LEU199 5.39 |
| ALA202 5.42 | | |
| SER203 5.43 | | |
| GLU206 5.46 | | |
| PHE207 5.47 | | PHE207 5.47 |

**ICL3**

| H3Ranta | H3Rago | H3Rapo |
|---|---|---|

**TM6**

| H3Ranta | H3Rago | H3Rapo |
|---|---|---|
| | LEU249 6.37 | |
| TRP253 6.48 | | |
| TYR256 6.51 | | TYR256 6.51 |
| MET260 6.55 | | MET260 6.55 |
| ARG263 6.58 | TYR263 6.51 | ARG263 6.58 |

**ECL3**

| H3Ranta | H3Rago | H3Rapo |
|---|---|---|

**TM7**

| H3Ranta | H3Rago | H3Rapo |
|---|---|---|
| TYR276 7.35 | | TYR276 7.35 |
| GLU277 7.36 | | |
| SER279 7.38 | | |
| PHE280 7.39 | | PHE280 7.39 |
| LEU283 2.42 | TYR283 7.35 | LEU283 2.42 |
| TRP284 7.43 | GLU284 7.36 | TRP284 7.43 |
| ASN286 7.45 | | ASN286 7.45 |
| SER287 7.46 | PHE287 7.39 | SER287 7.46 |
| ASN290 7.49 | | ASN290 7.49 |
| | TRP291 7.43 | |
| | SER294 7.49 | |

**ICL4**

| H3Ranta | H3Rago | H3Rapo |
|---|---|---|

**H8**

| H3Ranta | H3Rago | H3Rapo |
|---|---|---|

**C-ter**

| H3Ranta | H3Rago | H3Rapo |
|---|---|---|

Table S2. Amino acid residues involved in the formation of the receptor cavity for the antagonist, agonist and apo systems. Small upper left table: water population in the cavity.

**H3Ranta**

| Cluster | 1 | |
|---|---|---|
| TM1 | LEU54 | 1.52 |
| TM1 | ALA58 | 1.56 |
| TM1/TM2 | LEU65 | 12.50 |
| **Cluster** | **2** | |
| Nter/TM1 | ALA23 | 0.00 |
| TM2 | PHE86 | 2.56 |
| TM2 | CYS87 | 2.57 |
| TM2 | LEU90 | 2.60 |
| TM2/TM3 | TRP100 | 23.50 |
| TM3 | LEU106 | 3.24 |
| TM3 | TRP110 | 3.28 |
| **Cluster** | **3** | |
| TM2 | LEU82 | 2.52 |
| TM2 | VAL83 | 2.53 |
| TM3 | VAL113 | 3.31 |
| TM3 | LEU117 | 3.35 |
| **Cluster** | **4** | |
| TM2 | LEU73 | 2.43 |
| TM3 | ILE128 | 3.46 |
| TM5 | ILE221 | 5.61 |
| TM6 | LEU242 | 6.37 |
| TM6 | 2ILE245 | 6.40 |
| TM7 | CYS297 | 7.56 |
| **Cluster** | **5** | |
| TM3 | LEU109 | 3.27 |
| TM3 | VAL112 | 3.30 |
| TM4 | LEU166 | 4.56 |
| **Cluster** | **6** | |
| TM6 | ALA254 | 6.49 |
| TM6 | PRO255 | 6.50 |
| TM6 | ILE262 | 6.57 |
| TM6/TM7 | VAL271 | 67.00 |

**H3Rago – sec1**

| Cluster | 1 | |
|---|---|---|
| TM1 | VAL49 | 1.47 |
| TM1 | LEU54 | 1.47 |
| TM1 | MET56 | 1.54 |
| TM1 | LEU57 | 1.55 |
| TM1 | ALA58 | 1.56 |
| TM1 | PHE59 | 1.57 |
| TM1 | ALA61 | 1.59 |
| TM1/TM2 | LEU65 | 12.50 |
| TM7 | PRO302 | 7.54 |
| **Cluster** | **2** | |
| Nter/TM1 | ALA26 | 0.00 |
| TM2/TM3 | TRP100 | 23.50 |
| TM3 | CYS107 | 3.25 |
| **Cluster** | **3** | |
| TM2 | LEU76 | 2.46 |
| TM3 | ILE125 | 3.43 |
| TM5 | VAL214 | 5.54 |
| TM6 | LEU249 | 6.37 |
| **Cluster** | **4** | |
| TM3 | PHE133 | 3.51 |
| TM3 | VAL136 | 3.54 |
| TM5 | PHE216 | 5.56 |
| **Cluster** | **5** | |
| TM3 | VAL112 | 3.30 |
| TM3 | LEU116 | 3.34 |
| TM3 | ALA119 | 3.37 |
| TM4 | LEU166 | 4.56 |
| TM5 | ALA202 | 5.42 |
| **Cluster** | **6** | |
| TM1 | LEU44 | 1.42 |
| TM3 | VAL122 | 3.40 |
| TM3 | VAL126 | 3.44 |
| TM3 | ILE128 | 3.46 |
| TM5 | PHE207 | 5.47 |
| TM5 | PHE211 | 5.51 |
| TM6 | ILE255 | 6.43 |

**H3Rago – sec2**

| Cluster | 1 | |
|---|---|---|
| TM1 | VAL49 | 1.47 |
| TM1 | LEU54 | 1.52 |
| TM1 | MET56 | 1.54 |
| TM1 | LEU57 | 1.55 |
| TM1 | ALA58 | 1.56 |
| TM1 | PHE59 | 1.57 |
| TM1 | ALA61 | 1.59 |
| TM1/TM2 | LEU65 | 12.50 |
| **Cluster** | **2** | |
| TM2 | LEU82 | 2.53 |
| TM2 | PHE86 | 2.56 |
| TM3 | VAL112 | 3.30 |
| **Cluster** | **3** | |
| TM3 | ALA119 | 3.37 |
| TM/TM5 | LEU177 | 45.39 |
| TM4/TM5 | PHE192 | 45.54 |
| TM5 | PHE193 | 45.55 |
| TM5 | ALA202 | 5.42 |
| TM6 | LEU266 | 6.54 |
| **Cluster** | **4** | |
| TM3 | CYS118 | 3.36 |
| TM3 | VAL122 | 3.40 |
| TM5 | PHE207 | 5.47 |
| TM5 | PHE211 | 5.51 |
| TM6 | ILE255 | 6.43 |
| TM6 | PHE256 | 6.44 |
| **Cluster** | **5** | |
| TM3 | ILE125 | 3.43 |
| TM3 | 1VAL126 | 3.44 |
| TM5 | VAL214 | 5.54 |
| TM5 | PHE216 | 5.56 |
| TM6 | LEU249 | 6.37 |
| TM6 | ILE252 | 6.40 |
| TM7 | CYS304 | 7.56 |
| **Cluster** | **6** | |
| TM6 | LEU265 | 6.53 |
| TM6 | ILE269 | 6.57 |
| TM6/TM7 | VAL278 | 76.00 |
| TM6/TM7 | VAL279 | 7.31 |

**H3Rago – sec3**

| Cluster | 1 | |
|---|---|---|
| TM1 | LEU54 | 1.52 |
| TM1 | LEU57 | 1.55 |
| TM1 | ALA58 | 1.56 |
| TM1 | PHE59 | 1.57 |
| TM1 | ALA61 | 1.59 |
| TM1/TM2 | LEU65 | 12.5 |
| TM7 | PRO302 | 7.54 |
| **Cluster** | **2** | |
| TM2 | LEU82 | 2.52 |
| TM2 | PHE86 | 2.56 |
| TM3 | VAL112 | 3.30 |
| **Cluster** | **3** | |
| TM3 | LEU111 | 3.29 |
| TM3 | ALA119 | 3.37 |
| TM4/TM5 | PHE192 | 45.54 |
| TM5 | ILE200 | 5.40 |
| TM5 | ALA202 | 5.42 |
| TM6 | LEU266 | 6.54 |
| **Cluster** | **4** | |
| TM3 | ILE125 | 3.43 |
| TM2 | LEU76 | 2.46 |
| TM5 | VAL214 | 5.54 |
| TM3 | ILE128 | 3.46 |
| **Cluster** | **5** | |
| TM2 | PHE71 | 2.41 |
| TM2 | PHE72 | 2.42 |
| TM2 | LEU73 | 2.43 |
| TM3 | PHE123 | 3.41 |
| TM3 | LEU127 | 3.45 |
| TM4 | LEU158 | 4.48 |
| **Cluster** | **6** | |
| TM3 | VAL122 | 3.40 |
| TM3 | VAL126 | 3.44 |
| TM5 | PHE207 | 5.47 |
| TM5 | PHE208 | 5.48 |
| TM5 | PRO210 | 5.50 |
| TM5 | PHE211 | 5.51 |

**H3Rago – sec4**

| Cluster | 1 | |
|---|---|---|
| Nter/TM1 | ALA23 | 0.00 |
| Nter/TM1 | ALA26 | 0.00 |
| Nter/TM1 | PHE29 | 0.00 |
| Nter/TM1 | ALA31 | 1.29 |
| **Cluster** | **2** | |
| TM1 | LEU54 | 1.52 |
| TM1 | LEU57 | 1.55 |
| TM1 | ALA58 | 1.56 |
| TM1 | PHE59 | 1.57 |
| TM1/TM2 | LEU65 | 12.50 |
| TM7 | PRO302 | 7.54 |
| **Cluster** | **3** | |
| TM2 | LEU90 | 2.60 |
| TM2/TM3 | TRP100 | 23.50 |
| TM2/TM3 | PHE102 | 23.52 |
| TM3 | CYS107 | 3.25 |
| **Cluster** | **4** | |
| TM2 | LEU76 | 2.46 |
| TM3 | VAL122 | 3.40 |
| TM3 | PHE123 | 3.41 |
| TM3 | ILE125 | 3.43 |
| TM3 | VAL126 | 3.44 |
| TM3 | ILE128 | 3.46 |
| TM5 | PHE207 | 5.47 |
| TM5 | PHE211 | 5.51 |
| TM5 | VAL214 | 5.54 |
| TM6 | LEU249 | 6.37 |
| **Cluster** | **5** | |
| TM3 | LEU111 | 3.29 |
| TM3 | VAL112 | 3.30 |
| TM3 | LEU116 | 3.34 |
| TM4 | ALA163 | 4.53 |
| **Cluster** | **6** | |
| TM3 | ALA119 | 3.37 |
| TM4 | LEU166 | 4.56 |
| TM5 | ALA202 | 5.42 |
| **Cluster** | **7** | |
| TM3 | VAL136 | 3.54 |
| TM5 | ILE221 | 5.61 |
| TM5 | ILE225 | 5.65 |
| **Cluster** | **8** | |
| TM6 | LEU265 | 6.53 |
| TM6 | ILE269 | 6.57 |
| TM6 | ALA272 | 6.60 |

**H3Rapo**

| Cluster | 1 | |
|---|---|---|
| TM1 | MET41 | 1.39 |
| TM1 | ILE45 | 1.43 |
| TM2 | LEU82 | 2.52 |
| TM2 | VAL83 | 2.53 |
| TM3 | VAL113 | 3.31 |
| TM6 | PHE249 | 6.44 |
| TM6 | ILE248 | 6.43 |
| TM7 | PHE280 | 7.39 |
| TM7 | TRP281 | 7.40 |
| TM7 | LEU283 | 7.42 |
| TM7 | VAL289 | 7.48 |
| **Cluster** | **2** | |
| TM1 | VAL49 | 1.47 |
| TM1 | VAL55 | 1.53 |
| TM1 | MET56 | 1.54 |
| TM1 | ALA58 | 1.56 |
| TM1 | PHE59 | 1.57 |
| TM1/MT2 | SER64 | 12.49 |
| TM1/TM2 | LEU65 | 12.50 |
| TM7 | CYS297 | 7.56 |
| **Cluster** | **3** | |
| TM2 | PHE86 | 2.56 |
| TM2 | CYS87 | 2.57 |
| TM2 | LEU90 | 2.60 |
| TM1/TM2 | TRP100 | 23.50 |
| TM1/TM2 | PHE102 | 23.52 |
| TM3 | CYS107 | 3.25 |
| **Cluster** | **4** | |
| TM3 | VAL122 | 3.40 |
| TM3 | 1VAL126 | 3.44 |
| TM5 | PRO210 | 5.50 |
| TM5 | VAL214 | 5.54 |
| **Cluster** | **5** | |
| TM1 | LEU54 | 1.52 |
| TM1 | LEU57 | 1.55 |
| TM7 | PRO295 | 7.54 |

Table S3. Hydrophobic clusters at 90% occupancy formed by at least three side chains for the antagonist-H3R complex, the agonist-H3R complex and the apo receptor, respectively.

**H3Ranta**

| cluster | 1 | |
|---|---|---|
| TM2/TM3 | TRP100 | 23.50 |
| TM2 | PHE86 | 2.56 |
| TM3 | TRP110 | 3.28 |

**H3Rago – sec1**

(no clusters listed)

**H3Rago – sec2**

| cluster | 1 | |
|---|---|---|
| TM5 | PHE207 | 5.47 |
| TM5 | PHE211 | 5.51 |
| TM6 | PHE256 | 6.44 |

**H3Rago – sec3**

| cluster | 1 | |
|---|---|---|
| TM2 | PHE72 | 2.42 |
| TM3 | PHE123 | 3.41 |
| TM2 | PHE71 | 2.41 |
| **cluster** | **2** | |
| TM5 | PHE208 | 5.48 |
| TM5 | PHE207 | 5.47 |
| TM5 | PHE211 | 5.51 |

**H3Rago – sec4**

| cluster | 1 | |
|---|---|---|
| TM3 | PHE123 | 3.41 |
| TM5 | PHE207 | 5.47 |
| TM5 | PHE211 | 5.51 |

**H3Rapo**

| cluster | 1 | |
|---|---|---|
| TM7 | PHE280 | 7.39 |
| TM7 | TRP281 | 7.40 |
| TM6 | PHE249 | 6.44 |
| **cluster** | **2** | |
| TM2 | PHE86 | 2.56 |
| TM1/TM2 | TRP100 | 23.50 |
| TM1/TM2 | PHE102 | 23.52 |

***TMx/TMy = Loop between corresponding TMs**

Table S4. Aromatic residue clusters at 90% occupancy formed by at least three side chains for the antagonist-H3R complex, the agonist-H3R complex and the apo receptor, respectively.

| | | | Residence time | | | Residence time | | | Residence time | | | Residence time | | | Residence time |
|---|---|---|---|---|---|---|---|---|---|---|---|---|---|---|---|
| | **HR3anta** | | | | | | | | | | | | | | |
| DPPC | 125 | | 99.98% | 110 | | 99.96% | 161 | | 99.94% | 79 | | 99.78% | 33 | | 98.64% |
| Amino acid residue | LEU296 | 7.55 | 98.24% | PHE123 | 3.41 | 97.55% | LEU158 | 4.48 | 99.43% | PHE29 | 0.00 | 99.16% | TYR197 | 5.37 | 93.97% |
| | ILE244 | 6.39 | 98.09% | LEU205 | 5.45 | 96.59% | ARG154 | 4.44 | 99.04% | TRP281 | 7.40 | 97.01% | ASN195 | 45.57 | 89.37% |
| | LYS240 | 6.35 | 92.74% | LEU127 | 3.45 | 95.87% | LYS155 | 4.45 | 92.68% | TYR274 | 7.33 | 94.92% | PHE198 | 5.38 | 87.40% |
| | CYS297 | 7.56 | 89.91% | TYR130 | 3.48 | 94.89% | PHE123 | 3.41 | 92.10% | THR278 | 7.37 | 89.16% | TRP196 | 5.36 | 74.68% |
| | LYS237 | 6.32 | 88.37% | VAL126 | 3.44 | 85.24% | ARG151 | 4.41 | 89.66% | GLY26 | 0.00 | 78.49% | LEU165 | 4.55 | 72.77% |
| | HIS298 | 78.00 | 87.19% | LEU134 | 3.52 | 85.11% | ARG150 | 4.40 | 86.99% | SER30 | 0.00 | 77.94% | LEU166 | 4.56 | 56.57% |
| | ILE248 | 6.43 | 86.94% | ARG151 | 4.41 | 75.69% | ALA119 | 3.37 | 84.45% | ALA285 | 7.44 | 77.87% | LEU162 | 4.52 | 52.38% |
| | SER241 | 6.36 | 86.02% | VAL122 | 3.40 | 72.52% | LEU162 | 4.52 | 79.77% | LEU282 | 7.41 | 77.13% | THR201 | 5.41 | 52.16% |
| | LEU293 | 7.52 | 82.05% | ALA119 | 3.37 | 62.11% | SER120 | 3.38 | 77.42% | GLU277 | 7.36 | 73.74% | PRO169 | 4.59 | 51.81% |
| | ARG302 | 8.51 | 79.81% | ARG138 | 3.56 | 58.04% | VAL159 | 4.49 | 76.41% | TRP33 | 1.31 | 73.39% | | | |
| | VAL289 | 7.48 | 76.21% | THR201 | 5.41 | 52.29% | LEU116 | 3.34 | 64.88% | LEU40 | 1.38 | 64.02% | | | |
| | LEU251 | 6.46 | 58.81% | LEU212 | 5.52 | 52.28% | | | | ARG27 | 0.00 | 61.61% | | | |
| | | | | | | | | | | ALA31 | 0.00 | 52.52% | | | |
| | | | | | | | | | | ASP273 | 7.32 | 51.81% | | | |

| | | | Residence time | | | Residence time | | | Residence time | | | Residence time |
|---|---|---|---|---|---|---|---|---|---|---|---|---|
| DPPC | 165 | | 98.13% | 184 | | 85.65% | 168 | | 84.83% | 43 | | 82.97% |
| Amino acid residue | LYS240 | 6.35 | 78.55% | TYR130 | 3.48 | 69.46% | VAL60 | 1.58 | 57.42% | LEU109 | 3.27 | 82.89% |
| | TYR222 | 5.62 | 78.16% | PHE133 | 3.51 | 63.47% | MET56 | 1.54 | 53.52% | TYR167 | 4.57 | 82.23% |
| | ARG231 | 6.26 | 74.41% | LEU212 | 5.52 | 63.44% | LEU57 | 1.55 | 51.13% | GLY105 | 3.23 | 81.49% |
| | LEU251 | 6.46 | 72.63% | THR137 | 3.55 | 62.40% | | | | ALA170 | 4.60 | 77.59% |
| | SER247 | 6.42 | 68.94% | ARG138 | 3.56 | 60.11% | | | | ILE171 | 4.61 | 77.55% |
| | ILE244 | 6.39 | 65.96% | PHE208 | 5.48 | 58.10% | | | | ARG104 | 3.22 | 76.98% |
| | ILE225 | 5.65 | 60.48% | LEU134 | 3.52 | 56.58% | | | | LYS108 | 3.26 | 74.05% |
| | ALA243 | 6.38 | 57.24% | PHE217 | 5.57 | 51.97% | | | | LEU106 | 3.24 | 65.22% |
| | LYS236 | 6.31 | 51.21% | | | | | | | SER173 | 4.63 | 65.11% |
| | GLN226 | 5.66 | 50.87% | | | | | | | TRP160 | 4.50 | 61.88% |
| | ASN218 | 5.58 | 50.20% | | | | | | | PHE164 | 4.54 | 59.13% |
| | | | | | | | | | | ALA163 | 4.53 | 58.24% |
| | | | | | | | | | | VAL113 | 3.31 | 56.30% |
| | | | | | | | | | | GLU175 | 45.37 | 55.41% |

| | | | Residence time | | | Residence time | | | Residence time | | | Residence time |
|---|---|---|---|---|---|---|---|---|---|---|---|---|
| DPPC | 31 | | 82.76% | 149 | | 82.46% | 131 | | 81.48% | 166 | | 81.43% |
| Amino acid residue | TRP196 | 5.36 | 75.41% | VAL292 | 7.51 | 81.81% | PHE216 | 5.56 | 62.39% | VAL153 | 4.43 | 52.00% |
| | ILE200 | 5.40 | 72.68% | PRO291 | 7.50 | 79.11% | THR137 | 3.55 | 61.67% | | | |
| | THR204 | 5.44 | 71.88% | PRO295 | 7.54 | 78.79% | PHE133 | 3.51 | 58.19% | | | |
| | PHE208 | 5.48 | 62.75% | ARG302 | 8.51 | 78.78% | LEU212 | 5.52 | 51.67% | | | |
| | HIS267 | 67.00 | 58.52% | ALA47 | 1.45 | 78.63% | | | | | | |
| | ALA264 | 6.59 | 57.63% | GLY51 | 1.49 | 77.95% | | | | | | |
| | SER203 | 5.43 | 57.39% | LEU54 | 1.52 | 77.17% | | | | | | |
| | PHE207 | 5.47 | 57.37% | PHE305 | 8.54 | 76.24% | | | | | | |
| | LEU199 | 5.39 | 56.78% | LEU296 | 7.55 | 74.25% | | | | | | |
| | | | | ALA288 | 7.47 | 73.66% | | | | | | |
| | | | | THR48 | 1.46 | 70.94% | | | | | | |
| | | | | VAL289 | 7.48 | 70.17% | | | | | | |
| | | | | LEU50 | 1.48 | 67.78% | | | | | | |
| | | | | LEU44 | 1.42 | 52.75% | | | | | | |

Table S5 (see below).

| | | | | | | | | | | | | | | | | | | | | | |
|---|---|---|---|---|---|---|---|---|---|---|---|---|---|---|---|---|---|---|---|---|---|
| | **H3Rago** | | | | | | | | | | | | | | | | | | | | |
| | **sec1** | | | | | | | | | | | | | | | | | | | | |
| | | | Residence time | | | Residence time | | | Residence time | | | Residence time | | | Residence time | | | Residence time | | | Residence time |
| DPPC | 2 | | 100.00% | 47 | | 98.43% | 183 | | 94.05% | 7 | | 93.16% | 65 | | 88.81% | 22 | | 85.25% | 59 | | 85.15% |
| Amino acid residue | ILE251 | 6.39 | 87.11% | TYR222 | 5.62 | 98.43% | VAL92 | 2.62 | 77.96% | VAL299 | 7.51 | 59.14% | LEU303 | 7.55 | 85.18% | ARG66 | 12.51 | 85.21% | TYR222 | 5.62 | 80.87% |
| | LYS247 | 6.35 | 81.72% | ALA250 | 6.38 | 98.18% | ILE88 | 2.58 | 75.50% | ARG309 | 8.51 | 53.02% | ARG309 | 8.51 | 76.62% | VAL60 | 1.58 | 85.21% | GLN226 | 5.66 | 65.76% |
| | ILE255 | 6.43 | 81.36% | ASN218 | 5.58 | 98.07% | PRO93 | 2.63 | 75.20% | THR313 | 8.55 | 52.78% | VAL299 | 7.51 | 59.36% | MET56 | 1.54 | 84.93% | LEU219 | 5.59 | 57.52% |
| | LEU303 | 7.55 | 77.37% | LYS247 | 6.35 | 95.25% | LEU96 | 2.66 | 72.26% | | | | TYR306 | 0.00 | 58.67% | PHE59 | 1.57 | 84.81% | | | |
| | LEU258 | 6.46 | 70.21% | THR215 | 5.55 | 95.16% | PRO89 | 2.59 | 62.70% | | | | | | | LEU74 | 2.44 | 84.65% | | | |
| | ARG309 | 8.51 | 69.49% | LYS243 | 6.31 | 93.09% | THR97 | 2.67 | 60.71% | | | | | | | PHE81 | 2.51 | 81.07% | | | |
| | CYS304 | 7.56 | 68.97% | LEU219 | 5.59 | 93.00% | | | | | | | | | | ILE78 | 2.48 | 80.02% | | | |
| | LEU300 | 7.52 | 67.95% | ALA246 | 6.34 | 91.71% | | | | | | | | | | SER63 | 12.48 | 73.90% | | | |
| | SER254 | 6.42 | 65.25% | VAL253 | 6.41 | 90.88% | | | | | | | | | | LEU57 | 1.55 | 69.68% | | | |
| | ALA250 | 6.38 | 62.41% | SER254 | 6.42 | 88.47% | | | | | | | | | | PHE71 | 2.41 | 64.01% | | | |
| | SER248 | 6.36 | 61.75% | GLY257 | 6.45 | 86.60% | | | | | | | | | | ALA61 | 1.59 | 61.09% | | | |
| | LYS244 | 6.32 | 58.71% | ILE225 | 5.65 | 84.01% | | | | | | | | | | LEU82 | 2.52 | 59.72% | | | |
| | VAL296 | 7.48 | 58.45% | VAL214 | 5.54 | 83.60% | | | | | | | | | | ASP62 | 1.60 | 52.81% | | | |
| | TYR306 | 0.00 | 49.86% | ILE221 | 5.61 | 81.21% | | | | | | | | | | | | | | | |
| | | | | PHE211 | 5.51 | 79.03% | | | | | | | | | | | | | | | |
| | | | | TRP260 | 6.48 | 78.62% | | | | | | | | | | | | | | | |
| | | | | LEU258 | 6.46 | 57.06% | | | | | | | | | | | | | | | |
| | | | | PHE256 | 6.44 | 51.17% | | | | | | | | | | | | | | | |
| | | | | ALA261 | 6.49 | 50.16% | | | | | | | | | | | | | | | |

Table S6 (see below).

| | | | | | | | | | | | | | | |
|---|---|---|---|---|---|---|---|---|---|---|---|---|---|---|
| | **H3Rapo** | | | | | | | | | | | | | | |
| | | | Residence time | | | Residence time | | | Residence time | | | Residence time | | | Residence time |
| DPPC | 184 | | 100.00% | 151 | | 98.25% | 133 | | 96.42% | 155 | | 88.47% | 128 | | 82.44% |
| Amino acid residue | TRP 281 | 7.40 | 100.00% | LEU109 | 3.27 | 98.74% | TRP33 | 1.31 | 73.40% | ARG27 | 0.00 | 79.35% | LEU258 | 6.53 | 80.12% |
| | GLU277 | 7.36 | 99.83% | LYS108 | 3.26 | 97.76% | LEU40 | 1.38 | 72.31% | PHE29 | 0.00 | 53.83% | ILE262 | 6.57 | 75.91% |
| | TYR274 | 7.33 | 99.27% | VAL112 | 3.3 | 95.42% | LEU37 | 1.35 | 59.57% | | | | PRO255 | 6.50 | 65.46% |
| | ALA285 | 7.44 | 98.93% | TYR167 | 4.57 | 94.57% | TYR274 | 7.33 | 58.91% | | | | TRP275 | 7.34 | 64.76% |
| | THR278 | 7.37 | 96.32% | VAL113 | 3.31 | 92.42% | | | | | | | CYS266 | 6.61 | 55.80% |
| | LEU44 | 1.42 | 95.96% | LEU116 | 3.34 | 92.05% | | | | | | | ALA254 | 6.49 | 53.68% |
| | LEU37 | 1.35 | 95.44% | ALA163 | 4.53 | 91.62% | | | | | | | LEU259 | 6.54 | 53.05% |
| | VAL95 | 2.65 | 95.14% | GLY105 | 3.23 | 89.95% | | | | | | | | | |
| | LEU40 | 1.38 | 90.32% | TRP160 | 4.5 | 82.08% | | | | | | | | | |
| | TYR91 | 2.61 | 90.18% | VAL159 | 4.49 | 82.04% | | | | | | | | | |
| | LEU282 | 7.41 | 88.36% | ARG104 | 3.22 | 78.15% | | | | | | | | | |
| | GLY98 | 2.68 | 86.11% | ALA170 | 4.6 | 66.44% | | | | | | | | | |
| | ALA288 | 7.47 | 74.47% | LEU82 | 2.52 | 66.34% | | | | | | | | | |
| | VAL289 | 7.48 | 73.43% | PHE164 | 4.54 | 61.74% | | | | | | | | | |
| | ASP273 | 7.32 | 70.32% | TYR176 | 45.38 | 61.73% | | | | | | | | | |
| | TRP33 | 1.31 | 65.56% | ILE171 | 4.61 | 61.59% | | | | | | | | | |
| | TYR94 | 2.64 | 63.54% | MET156 | 4.46 | 59.43% | | | | | | | | | |
| | LEU96 | 2.66 | 61.77% | LEU117 | 3.35 | 56.34% | | | | | | | | | |
| | LEU43 | 1.41 | 57.98% | | | | | | | | | | | | |
| | TRP284 | 7.43 | 53.40% | | | | | | | | | | | | |
| | THR97 | 2.67 | 52.68% | | | | | | | | | | | | |

Table S7 (see below).

Tables S5, S6 and S7. Lipid-binding sites on the inactive-state (antagonist-bound), active-state (agonist-bound), and constitutive-state receptor (apo), respectively.

| | H3Ranta | | H3Rago | | H3Rapo | |
|---|---|---|---|---|---|---|
| | No .aa | % | No. aa | % | No. aa | % |
| non-polar, aliphatic | | | | | | |
| GLY | 3 | 2.88% | 1 | 1.69% | 2 | 4.00% |
| ALA | 0 | 0.00% | 5 | 8.47% | 5 | 10.00% |
| VAL | 9 | 8.65% | 7 | 11.86% | 4 | 8.00% |
| LEU | 26 | 25.00% | 10 | 16.95% | 14 | 28.00% |
| ILE | 5 | 4.81% | 6 | 10.17% | 2 | 4.00% |
| MET | 1 | 0.96% | 1 | 1.69% | 1 | 2.00% |
| polar, non-charged | | | | | | |
| SER | 5 | 4.81% | 4 | 6.78% | 0 | 0.00% |
| THR | 6 | 5.77% | 3 | 5.08% | 2 | 4.00% |
| CYS | 1 | 0.96% | 1 | 1.69% | 1 | 2.00% |
| PRO | 3 | 2.88% | 2 | 3.39% | 1 | 2.00% |
| ASN | 2 | 1.92% | 1 | 1.69% | 0 | 0.00% |
| GLN | 1 | 0.96% | 0 | 0.00% | 0 | 0.00% |
| aromatic | | | | | | |
| PHE | 11 | 10.58% | 5 | 8.47% | 1 | 2.00% |
| TYR | 6 | 5.77% | 3 | 5.08% | 6 | 12.00% |
| TRP | 4 | 3.85% | 1 | 1.69% | 6 | 12.00% |
| charged | | | | | | |
| LYS | 6 | 5.77% | 4 | 6.78% | 1 | 2.00% |
| ARG | 11 | 10.58% | 4 | 6.78% | 2 | 4.00% |
| HIS | 1 | 0.96% | 0 | 0.00% | 0 | 0.00% |
| ASP | 1 | 0.96% | 1 | 1.69% | 1 | 2.00% |
| GLU | 2 | 1.92% | 0 | 0.00% | 1 | 2.00% |
| | 104 | 100.00% | 59 | 100.00% | 50 | 100.00% |

Table S8. Residue population in contact with the lipids grouped according to amino acid classes (non-polar aliphatic, uncharged polar, aromatic, and charged) for the antagonist, agonist and apo systems, respectively.

**Antagonist complex**

| Resid | Number of residues | With respect to all hydrophobic | Number of residues | With respect to all hydrophylic | Number of residues | With respect to all charged |
|---|---|---|---|---|---|---|
| Ala | 6 | 11.76% | | | | |
| Arg | | | | | 3 | 5.88% |
| Asn | | | | | | |
| Asp | | | | | | |
| Cys | | | | | | |
| Gln | | | | | | |
| Glu | | | | | | |
| Gly | | | | | | |
| Ile | 1 | 1.96% | | | | |
| Leu | 14 | 27.45% | | | | |
| Lys | | | | | 1 | 1.96% |
| Met | 1 | | | | | |
| Phe | 2 | 3.92% | | | | |
| Pro | 1 | | | | | |
| Ser | | | 1 | 1.96% | | |
| Thr | | | 6 | 11.76% | | |
| Trp | 3 | 5.88% | | | | |
| Tyr | | | 3 | 5.88% | | |
| Val | 8 | 15.69% | | | | |
| | | With respect to all exposed residues 70.59% | | With respect to all exposed residues 21.57% | | With respect to all exposed residues 7.84% |

**Agonist complex**

| Resid | Number of residues | With respect to all hydrophobic | Number of residues | With respect to all hydrophylic | Number of residues | With respect to all charged |
|---|---|---|---|---|---|---|
| Ala | 7 | 16.67% | | | | |
| Arg | | | | | 4 | 9.52% |
| Asn | | | | | | |
| Asp | | | | | | |
| Cys | | | | | | |
| Gln | | | | | | |
| Glu | | | | | | |
| Gly | | | | | | |
| Ile | 2 | 4.76% | | | | |
| Leu | 12 | 28.57% | | | | |
| Lys | | | | | 3 | 7.14% |
| Met | | | | | | |
| Phe | 4 | 9.52% | | | | |
| Pro | | | | | | |
| Ser | | | 2 | 4.76% | | |
| Thr | | | 3 | 7.14% | | |
| Trp | 1 | 2.38% | | | | |
| Tyr | | | | | | |
| Val | 4 | 9.52% | | | | |
| | | With respect to all exposed residues 71.43% | | With respect to all exposed residues 11.90% | | With respect to all exposed residues 16.67% |

**Apo receptor**

| Resid | Number of residues | With respect to all hydrophobic | Number of residues | With respect to all hydrophylic | Number of residues | With respect to all charged |
|---|---|---|---|---|---|---|
| Ala | 10 | 14.93% | | | | |
| Arg | | | | | 6 | 8.96% |
| Asn | | | | | | |
| Asp | | | | | 2 | 2.99% |
| Cys | | | | | | |
| Gln | | | | | | |
| Glu | | | 1 | 1.49% | | |
| Gly | | | 1 | 1.49% | | |
| Ile | 1 | 1.49% | | | | |
| Leu | 18 | 26.87% | | | | |
| Lys | | | | | 2 | 2.99% |
| Met | 1 | 1.49% | | | | |
| Phe | 3 | 4.48% | | | | |
| Pro | 1 | 1.49% | | | | |
| Ser | | | 1 | 1.49% | | |
| Thr | | | 6 | 8.96% | | |
| Trp | 3 | 4.48% | | | | |
| Tyr | | | 3 | 4.48% | | |
| Val | 8 | 11.94% | | | | |
| | | With respect to all exposed residues 67.16% | | With respect to all exposed residues 17.91% | | With respect to all exposed residues 14.93% |

***Hydrophobic class:*** *phe, met, trp, ile, val, leu, pro, ala*
***Hydrophylic class:*** *asn, cys, gln, gly, ser, thr, tyr*
***Charged class: arg****, asp, glu, his, lys*
*(Biophys J. 2007 Jul 1; 93(1): 225–231*

Table S9. Average composition of transmembrane surface amino acid residues. There are no exposed His residues.

| Anta | | Ago | | Apo | |
|---|---|---|---|---|---|
| MET41:TRP281 | 1.39:7.40 | MET41:TRP288 | 1.39:7.40 | MET41:TYR91 | 1.39:2.61 |
| | | | | MET41:TRP281 | 1.39:7.40 |
| MET41:TRP284 | 1.39:7.43 | | | MET41:TRP284 | 1.39:7.43 |
| | | | | ARG150:PRO291 | 1.50:7.50 |
| ASN69:ASP131 | 2.39:3.49 | | | MET56:PHE81 | 1.54:2.51 |
| ASN69:ARG132 | 2.39:3.50 | | | ASN69:ASP131 | 2.39:3.49 |
| | | | | ASN69:ARG132 | 2.39:3.50 |
| | | | | ASP80:PRO291 | 2.50:7.50 |
| ARG132:ASP131 | 3.50:3.49 | | | ASP114:TRP284 | 3.32:7.43 |
| ARG132:ASN224 | 3.50:5.64 | | | | |
| ARG132:ASP235 | 3.50:6.30 | | | | |
| PHE151:TRP253 | 5.51:6.48 | | | | |
| | | | | PHE207:TRP253 | 5.47:6.48 |
| MET260:TYR256 | 6.55:6.51 | MET267:PHE208 | 6.55:5.48 | TRP253:SER287 | 6.48:7.46 |
| | | MET267:TYR115 | 6.55:3.33 | MET260:TYR256 | 6.55:6.51 |
| MET260:PHE280 | 6.55:7.39 | MET267:TYR263 | 6.55:6.51 | | |
| | | TYR301:PHE308 | 7.53:8.50 | | |
| | | | | TYR294:PHE301 | 7.53:8.50 |
| | | | | TYR294:PHE305 | 7.53:8.54 |

Table S10. Selected pairs of residues establishing contacts according to inter-residue distances criteria of 6 Å between COMs of side chains. Includes ionic locks Arg3.50-Asp3.49, and Arg3.50-Asp6.30.
